# Subfamily C7 Raf-like kinases MRK1, RAF26, and RAF39 regulate immune homeostasis and stomatal opening in *Arabidopsis thaliana*

**DOI:** 10.1101/2023.11.29.569073

**Authors:** Márcia Gonçalves Dias, Bassem Doss, Anamika Rawat, Kristen R. Siegel, Tharika Mahathanthrige, Jan Sklenar, Paul Derbyshire, Thakshila Dharmasena, Emma Cameron, Cyril Zipfel, Frank L.H. Menke, Jacqueline Monaghan

## Abstract

The calcium-dependent protein kinase CPK28 is a regulator of immune homeostasis in multiple plant species. Here, we used a proteomics approach to uncover CPK28-associated proteins. We found that CPK28 associates with subfamily C7 Raf-like kinases MRK1, RAF26, and RAF39, and trans-phosphorylates RAF26 and RAF39. Metazoan Raf kinases function in mitogen-activated protein kinase (MAPK) cascades as MAPK kinase kinases (MKKKs). Although Raf-like kinases share some features with MKKKs, we found that MRK1, RAF26, and RAF39 are unable to trans-phosphorylate any of the 10 Arabidopsis MKKs. We show that MRK1, RAF26, and RAF39 localize to the cytosol and endomembranes, and we define redundant roles for these kinases in stomatal opening, immune-triggered reactive oxygen species (ROS) production, and resistance to a bacterial pathogen. Overall, our study suggests that C7 Raf-like kinases associate with and are phosphorylated by CPK28, function redundantly in stomatal immunity, and possess substrate specificities distinct from canonical MKKKs.

## Introduction

Plants encounter a variety of stressors in the environment that can negatively impact their growth and survival. The ability of plants to respond to danger signals such as drought, heat, cold, salinity, or pathogen attack, is critical to optimizing growth and reproduction in a changing environment. Sensing, integrating, and responding to stress is achieved through cellular signaling pathways that ultimately result in temporary genetic reprogramming. Signal transduction is largely achieved by protein kinases that phosphorylate target proteins to regulate their activity, localization, and binding partners. Protein kinases are highly diverse in terms of their substrate specificity and cellular localization, and their roles in plant stress pathways have been extensively documented. Many cell surface receptors involved in stress signaling are transmembrane receptor kinases (RKs) or receptor proteins (RPs) that bind ligands via their extracellular domain (DeFalco & Zipfel, 2021) and associate closely with several classes of intracellular protein kinases including receptor-like cytoplasmic kinases (RLCKs) (Liang & Zhou, 2018), mitogen-activated protein kinases (MAPKs) (Taj *et al.*, 2010), and calcium-dependent protein kinases (CDPKs) (Yip Delormel & Boudsocq, 2019). Although several targets of RLCKs, MAPKs, and CDPKs have been documented, a key challenge is to identify context-specific substrates of protein kinases and points of regulation in signaling pathways.

CDPKs participate in a broad range of cellular processes, including stomatal movement, hormone signaling, cell cycle and differentiation, seed development and germination, metabolic regulation, and pathogen defense (Yip Delormel & Boudsocq, 2019). The diversity and complexity of CDPK-mediated signaling pathways highlights their importance in plant growth, development, and adaptation to changing environmental conditions. There are 42 CDPKs encoded in *Arabidopsis thaliana* (hereafter: Arabidopsis), that can be separated into five subfamilies (I, II, III, IV, and CRK (CDPK-related kinases)) based on their phylogenetic relationships (Chen *et al.*, 2017; Yip Delormel & Boudsocq, 2019). CDPKs consist of a variable N-terminal domain, a protein kinase domain, an autoinhibitory junction domain (AIJ), and a C-terminal calmodulin-like domain. In the inactive state, CDPKs adopt a closed conformation in which the AIJ occupies the active site. Ca^2+^ binding to the calmodulin-like domain results in a drastic conformational change that derepresses the kinase by exposing the active site and allowing CDPKs to phosphorylate targets (Liese & Romeis, 2013).

The subfamily IV CDPK CPK28 is multi-functional, playing roles in plant growth and development (Matschi *et al.*, 2013), stress responses (Jin *et al.*, 2017; Hu *et al.*, 2021; Ding *et al.*, 2022b,a), and defense against pathogens (Monaghan *et al.*, 2014, 2015; Matschi *et al.*, 2015). CPK28 phosphorylates the E3 ubiquitin ligases PLANT U-BOX 25 (PUB25) and PUB26, enhancing their ability to poly-ubiquitinate the RLCK BOTRYTIS INDUCED KINASE 1 (BIK1), a common substrate of multiple receptors and a critical signaling node in plant immunity (Monaghan *et al.*, 2014; Wang *et al.*, 2018; DeFalco & Zipfel, 2021). The CPK28-PUB25/26 regulatory module buffers BIK1 protein accumulation to optimize immune output (Gonçalves Dias *et al.*, 2022). In the current study, we aimed to identify additional CPK28 binding partners in Arabidopsis using a proteomics approach. We found that CPK28 co-purifies with many protein kinases, including MIXED LINEAGE KINASE/RAF-RELATED KINASE 1 (MRK1). Because limited genome sequences were available at the time of discovery in 1997, MRK1 was named according to its possible relationship to mammalian mixed-lineage kinases (MLKs) or Raf kinases (Ichimura *et al.*, 1997). Metazoan rapidly accelerated fibrosarcoma (Raf) kinases are dual-specificity serine/threonine and tyrosine protein kinases that function in MAPK cascades. In mammals, the Ras-Raf-MEK-ERK pathway has been intensely studied and serves as a paradigm for membrane-to-nucleus signal transduction. In this pathway, binding of epidermal growth factor (EGF) to the EGF receptor at the plasma membrane results in activation and phosphorylation of its cytoplasmic tyrosine kinase domain. This activates the GTPase Ras, which then binds to and activates Raf, which serves as a MAPK kinase kinase (MKKK), phosphorylating and activating a MAPK kinase MEK, which then phosphorylates and activates a MAPK (originally named extracellular signal regulated kinase; ERK) (Terrell & Morrison, 2019). Reflecting the expansion of the protein kinase family in the plant kingdom, there are 20 MAPKs, 10 MKKs, and 80 MKKKs in Arabidopsis (González-Coronel *et al.*, 2021) – many more than in mammals. Despite their number, very little is known about MKKKs. Sequence homology defines three distinct subclasses known as MKKK, ZIK, and Raf-like kinases. There are 48 Raf-like kinases in Arabidopsis, divided into eleven subfamilies: B1-B4 and C1-C7 (Jonak *et al.*, 2002; González-Coronel *et al.*, 2021). Phylogenetic analyses indicate that plant Raf-like kinases do not cluster with metazoan MKKK or Raf kinases (Tang & Innes, 2002; Champion *et al.*, 2004) and are considered a plant (Pl)-specific family of tyrosine kinase-like (TKL) proteins (TKL-Pl-4) (Lehti-Shiu & Shiu, 2012). Despite this divergence, TKL-Pl-4 kinases share sequence features with metazoan Rafs and MLKs and therefore may function biochemically as MKKKs in MAPK cascades (Champion *et al.*, 2004; Lehti-Shiu & Shiu, 2012; González-Coronel *et al.*, 2021), however this has not been comprehensively studied.

MRK1 belongs to the C7 subfamily of Raf-like kinases, together with RAF26, RAF39, CONVERGENCE OF BLUE LIGHT AND CO_2_ 1 (CBC1), and CBC2 (Hiyama *et al.*, 2017). CBC1 and CBC2 are highly expressed in guard cells and have established roles in light-induced stomatal opening (Hiyama *et al.*, 2017). While stomatal pores play a critical role in controlling gas exchange and water transpiration, they also represent a point of entry for microbial pathogens (Melotto *et al.*, 2006), and immune-induced stomatal closure is a well-documented antimicrobial defense response (Melotto *et al.*, 2017). Here, we define redundant roles for MRK1, RAF26, and RAF39 in the inhibition of immune-triggered production of reactive oxygen species (ROS), and also demonstrate that MRK1, RAF26, and RAF39 function in stomatal opening which correlates to enhanced resistance to a bacterial pathogen. We show that MRK1, RAF26, and RAF39 localize intracellularly to the cytosol and endomembranes. We confirm that MRK1, RAF26, and RAF39 associate with CPK28 and that CPK28 can trans-phosphorylate RAF26 and RAF39 *in vitro.* We further show that MRK1, RAF26, and RAF39 are active kinases that are able to auto-phosphorylate *in vitro*. However, they are unable to trans-phosphorylate any of the 10 Arabidopsis MKKs *in vitro,* suggesting that they possess substrate specificities distinct from canonical MKKKs. Overall, our study reveals that C7 Raf-like kinases are CPK28 substrates that function redundantly in immune-triggered ROS production and light-induced stomatal opening, and provide evidence that they do not function as MKKKs.

## Materials and Methods

### Germplasm and plant growth conditions

*Arabidopsis thaliana* insertion mutant lines were obtained from the Arabidopsis Biological Resource Centre (ABRC) and genotyped to homozygosity using gene-specific primers in standard polymerase chain reactions (PCR). Double and triple mutants were generated by crossing, genotyped to homozygosity, and confirmed in subsequent generations. The T-DNA insertion site for the *mrk1-1* mutant was confirmed by Sanger sequencing a PCR-generated amplicon (Centre for Applied Genomics, Toronto). To assess if the insertion mutations resulted in lower gene expression, target genes were amplified using quantitative reverse transcription (qRT)-PCR. For this, leaf tissue was ground in liquid N_2_ and total RNA was extracted using the Aurum Total RNA Mini Kit (BioRad) according to the manufacturer’s instructions. Superscript III reverse transcriptase (Invitrogen) was used with oligo dT18 to generate cDNA according to the manufacturer’s instructions. cDNA was diluted and target gene expression assessed by qRT-PCR using gene-specific primers and SsoAdvanced Universal SYBR Green Supermix (BioRad). Detailed information regarding all germplasm generated or used in this study, including primers used for genotyping and qRT-PCR, is available in **Table S1**.

The experimental conditions used to grow and harvest samples from *cpk28-1/35S:CPK28-YFP, nsl1-1/35S:NSL1-YFP* and Col-0/*35S:Lti6B-GFP* for the proteomics screen was previously described (Bender *et al.*, 2017). Briefly, plants were grown on soil for 22 days under 10-h-light/8-h-dark cycle at 22°C in controlled environmental chambers at the John Innes Centre (Norwich Research Park). All other plants were grown in the Queen’s University Phytotron. For aseptic growth, Arabidopsis seeds were surface-sterilized with 40% bleach, sown on petri plates containing 0.5x Murashige and Skoog (MS) media (Cedarlane) and 0.8% agar, and stratified for 3-5 days at 4°C in the dark prior to exposure to light. For soil-grown plants, seeds were sown directly on potting soil (Sungro Sunshine Mix 1 or Fafard’s Agro G6 w. Coco) and seedlings were transplanted into pots as individual plants in 3” pots or as 6 plants/pot in 8” pots two weeks after sowing. Plants were grown in controlled growth chambers (BioChambers) with a 10-h-light/8-h-dark cycle at 22°C, with 30% relative humidity and a light intensity of 150 µE m^2^ s^-1^, top-watered when needed (typically every other day), and fertilized biweekly with a solution of 1.5 g/L 20:20:20 N:P:K. *Nicotiana benthamiana* seeds were sown directly on potting soil as above, transplanted as individual seedlings per pot, and grown in a dedicated growth chamber (Conviron) under similar conditions, except with a 16-h-light/8-h-dark cycle and fertilized weekly. Mite bags containing *Amblyseius swirskii* (Koppert) were added to each tray of plants bi-weekly to prevent pest infestations.

### Molecular cloning

Clones were generated using various methods as outlined in detail in **Table S1**. The 1,173 bp coding sequence of *MRK1* was amplified from DKLAT3G63260 (Popescu *et al.*, 2007) with Q5 *Taq* Polymerase (NEB) and Gateway-compatible pENTR-MRK1 clones were generated the using Gibson Assembly Master Mix (NEB) according to the manufacturer’s instructions. Gateway-compatible pTwistENTR vectors containing the coding sequences for *RAF26* (1,092 bp) or *RAF39* (1,137 bp) were synthesized by Twist BioSciences. An additional guanine was added to the inserts to maintain the first coding frame for C-terminal fusions following recombination into destination vectors. Recombination into the binary pK7FWG2 destination vector (Karimi *et al.*, 2002) for expression of *MRK1, RAF26*, and *RAF39* driven by the cauliflower mosaic virus (CaMV) *35S* promoter and C-terminally tagged with green fluorescent protein (GFP) was achieved using Gateway LR Clonase II (Invitrogen) according to the manufacturer’s instructions.

Vectors suitable for split-luciferase complementation were generated either by traditional digestion-ligation cloning or Gateway LR reactions. For digestion-ligation cloning, engineered 5’ and 3’ endonuclease sites flanking the target genes facilitated ligation into pCAMBIA1300-nLuc for *35S*-driven expression of recombinant proteins C-terminally tagged with the N-terminal 416 amino acids of firefly luciferase, or into pCAMBIA1300-cLuc for *35S*-driven expression of recombinant proteins N-terminally tagged with the C-terminal 153 amino acids of firefly luciferase (Chen *et al.*, 2008). The 2,685 bp coding sequence of *FER* was amplified from DKLAT3G51550 (Popescu *et al.*, 2007) with Q5 *Taq* Polymerase (NEB) and desalted using GenepHlow PCR Cleanup Kit (GeneAid) according to the manufacturer’s instructions. *MRK1, RAF26, RAF39,* and other coding sequences were synthesized by Twist BioSciences and rehydrated in pure water. Fragments and vector backbones were digested with appropriate endonucleases (NEB), desalted using GenepHlow PCR Cleanup Kit (GeneAid) and ligated with T4 DNA ligase (NEB), each step according to manufacturer’s directions. For Gateway-compatible vectors, we used pGWB-nLuc or pGWB-cLuc vectors, engineered from pCAMBIA1300-nLuc and pCAMBIA1300-cLuc (Yu *et al.*, 2020), obtained from Addgene. Entry vectors were either synthesized by Twist Biosciences or obtained from the ABRC, as outlined in **Table S1**. Recombination into destination vectors was achieved using Gateway LR Clonase II (Invitrogen) according to the manufacturer’s instructions. Whenever entry and destination vectors had the same antibiotic resistance markers, the entry vector backbone was linearized by endonuclease digestion prior to the LR reaction. Vectors suitable for expression and purification of His_6_-and/or glutathione S-transferase (GST)-tagged recombinant proteins in *Escherichia coli* were either cloned by Twist Biosciences into the pET28a+ backbone (EMD Biosciences) or cloned in-house into the pGex6p.1 backbone (GE Healthcare). When needed, mutations were incorporated directly at the synthesis stage.

Plasmids were transfected into *E. coli* Top10 cells, selected on petri plates with 1% agar and Luria-Bertani (LB) media (BioShop Canada) supplemented with appropriate antibiotics. Single colonies were used to inoculate liquid cultures and plasmids were extracted using the Presto Mini Plasmid Kit (GeneAid) according to manufacturer’s instructions. Successful assemblies were confirmed either by Sanger sequencing (Centre for Applied Genomics, Toronto ON, Canada) or by whole-plasmid sequencing (Plasmidosaurus, Eugene OR, USA). Information about all vectors used in this study, including previously published vectors, can be found in **Table S1.**

### *Agrobacterium*-mediated transient expression in *N. benthamiana*

Binary vectors were transfected into *Agrobacterium tumefaciens* strain GV3101 cells and grown on LB plates containing appropriate antibiotics. A single colony was transferred to liquid LB media with appropriate antibiotics, and grown for 12-16 h at 28°C. Cells were pelleted gently at 600 x *g*, resuspended in induction buffer (10 mM MgCl_2_, 10 mM MES pH 6.3), incubated for 2-3 h at room temperature on an orbital shaker, and normalized to OD_600_=0.2 using a microplate reader (SpectraMax Paradigm). Fully expanded upper leaves were selected from 4-week-old *N. benthamiana* plants for transformation. All constructs were co-transformed with viral suppressor P19 (Voinnet *et al.*, 2003), and leaves were infiltrated on the abaxial side using a 1 mL needleless syringe. Tissue for confocal imaging or split-luciferase complementation was harvested three days after infiltration.

### Split-luciferase complementation

*A. tumefaciens* carrying plasmids suitable for split-luciferase complementation assays (either pCAMBIA1300-n/cLuc or pGWB-n/cLuc; see **Table S1**) were used to transiently express proteins of interest in *N. benthamiana* as described above. Three days post infiltration, leaf disks (n=12) were collected using a 4 mm biopsy punch and placed in 100 μL of double-distilled water (ddH_2_O) in a white 96 well plate. Once all samples were collected, the water was replaced with 50 μL 1 mM D-Luciferin (Gold Biotechnology), incubated in the dark for 15 minutes, and luminescence recorded in a plate reader with an integration time of 1 s/well (SpectraMax Paradigm).

### Confocal microscopy

Leaf samples were collected with a 4 mm biopsy punch and mounted abaxial-side-up on a glass slide in a drop of water. Fluorescent proteins were excited with a 488 Argon laser and imaged using separate channels to detect emission of GFP (510-540 nm) or RFP (635-680 nm). For co-localization we used ER-mCherry, which was created by translationally fusing mCherry with an N-terminal secretion signal and a C-terminal HDEL sequence (Nelson *et al.*, 2007), and BRI1-mRFP (Saile *et al.*, 2021). Images were taken using a Zeiss LSM710 confocal microscope in the Biology Department at Queen’s University and processed using Zeiss Zen Software.

### Immune assays

Immunogenic flg22, elf18, and AtPep1 peptides were synthesized by EZ Biotech (Indiana USA). Immune-induced ROS production was performed on 4-to 5-week-old soil-grown plants as previously described (Bredow *et al.*, 2019). Immune-induced activation of MAPKs was performed on 2-week-old sterile seedlings as previously described (Monaghan *et al.*, 2014). Bacterial infections were performed on 4-to 5-week-old soil-grown plants. For spray-inoculation, *Pseudomonas syringae* pv. *tomato* (*Pst*) DC3000 was cultured at 28°C in LB media supplemented with rifampicin. Cells were gently pelleted and diluted to OD_600_=0.02 (10^7^ cfu/mL) in 10 mM MgCl_2_ and spray-inoculated onto 4-week-old plants until run-off. Right before spraying, 0.04% Silguard was added as a surfactant (Mireault *et al.*, 2014). Three days after inoculation, leaf tissue was harvested using a 4-mm biopsy punch and homogenized in 10 mM MgCl_2_. Four samples per genotype were collected by combining leaf discs from three different plants, serially diluted in 10 mM MgCl_2_, and bacterial growth was determined by expressing the number of colony forming units (cfu) per leaf area. Syringe-inoculations were performed similarly, however *Pst* DC3000 was diluted to OD_600_=0.0002 (10^5^ cfu/mL) in 10 mM MgCl_2_ with no surfactant.

Stomatal apertures were measured across four middle-aged leaves of 4-to 5-week-old soil-grown plants. The middle portion of each leaf was cut into three squares, avoiding the petiole, midrib, leaf base, tip, and margins. The leaf samples were placed in a buffer containing 50 mM KCl and 10 mM MES, pH ∼6 in a sterile 12 well plate, covered with a transparent lid, and placed in a growth chamber for 3 h. Following this stomatal opening period, the leaf squares were separated into T0, T1, and T3 sampling groups and incubated with 1 μM flg22 for 60 min (T1) or 180 min (T3) to induce stomatal closing and re-opening. At the appropriate time point, the abaxial sides of the leaf squares were mounted to a piece of double-sided tape attached to a microscope slide and carefully scraped using a razor blade until only the epidermal layer was left. Multiple fields-of-view of the epidermal tissue layer, including stomata, were imaged using a Zeiss Axioplan microscope with a 40X oil immersion lens objective. Images and scales were converted to JPEG files using Zeiss Zen software. The width and length of 120 individual stomata per time point for each genotype were measured using Image J software (Schindelin *et al.*, 2015) and converted to aperture values in R.

### Protein purification

#### Proteins expressed in plant tissue

Relatively equal amounts of *N. benthamiana* tissue (twelve 4 mm leaf discs per sample) were flash-frozen and ground to a fine powder in liquid N_2_, and proteins were extracted in standard Laemmli Buffer at 80°C for 10 minutes prior to SDS-PAGE and immunoblotting.

#### Proteins purified from E. coli

All proteins were expressed and purified from *E. coli* strain BL21 using the constructs outlined in **Table S1**. The cultures were grown at 37°C in LB containing appropriate antibiotics until the OD_600_ reached 0.7-0.8. Protein expression was induced by adding 0.5 mM or 1 mM of β-D-1-thio-galactopyranoside (IPTG) with shaking for 20 h at 28°C. Bacterial cells were harvested at 3,234 x *g* for 25 min at 4°C. The His_6_-tagged proteins were resuspended in extraction buffer consisting of 50 mM Tris·HCl (pH 7.5), 100 mM NaCl, and 1 mM phenylmethylsulfonyl fluoride (PMSF). The GST-tagged proteins were resuspended in phosphate-buffered saline (PBS) (Thermo Fisher) containing 1 mM dithiothreitol (DTT) and 1 mM PMSF. Cells were lysed by passing the resuspended pellets three times through a French Press G-M® High Pressure Cell Disruption (Clifton, NJ, USA). Lysates were clarified by centrifugation at 15,400 × *g* for 40 min at 4°C. The supernatants were loaded into a conical tube containing either nickel-nitriloacetic acid (HisPur™ 25215, Thermo Fisher Scientific) or glutathione agarose beads (G4510, Sigma Aldrich) with shaking for 1-2 hours at 4°C. His_6_ proteins were eluted from Ni-NTA beads by sequential washes with extraction buffer containing different imidazole concentrations (10 mM, 25 mM, 50 mM, 250 mM and 500 mM). Elution fractions were dialyzed in 2,000 volumes of 25 mM Tris⋅HCl (pH 7.5), 50 mM NaCl, and 1 mM DTT overnight at 4°C. GST proteins were eluted from glutathione agarose beads by washing with the elution buffer (50 mM Tris⋅HCl (pH 7.5), 10 mM reduced glutathione, 5 mM DTT). All proteins were concentrated using Amicon Ultra-15 centrifugal filter unit (10 or 30 KDa MWCO, MilliporeSigma). Protein concentrations were determined using Bradford reagent (23200, Thermo Fisher) and aliquots were flash frozen in liquid N_2_ and stored at −80°C until use.

### *In vitro* kinase assays

Auto-phosphorylation assays were performed using 2 μg kinase in a buffer containing 50 mM Tris-HCl (pH 8.0), 25 mM MgCl_2_, 25 mM MnCl_2_, 5 mM DTT, 5 μM ATP and 0.5-2 μCi γP^32^-ATP. Trans-phosphorylation assays used 2 μg kinase and 4 μg substrate in the same buffer. The buffer used in the trans-phosphorylation assays with His_6_-MBP-CPK28 contained 500 μM CaCl_2_ and no MnCl_2_. All reactions were incubated for 60 minutes at 30°C. Reactions were stopped by adding 6× Laemmli buffer and heating at 80 °C for 5 min. Proteins were separated in 10% SDS-PAGE gel at 80 V for 30 min followed by 150 V for 1 h in 1x SDS running buffer (25 mM Tris-HCl pH 6.8, 190 mM glycine, 0.1% (w/v) SDS). The gels were sandwiched between two sheets of transparency film, exposed to a storage phosphor screen (Molecular Dynamics) overnight and visualized using a Typhoon 8600 Imager (Molecular Dynamics/Amersham). Gels were stained with Coomassie Brilliant Blue (CBB) R-250 (MP Biomedicals) or SimplyBlue SafeStain (Invitrogen; CBB G-250) and scanned using an HP Officejet Pro 8620.

### SDS-PAGE and immunoblotting

Samples were loaded on a 10% SDS polyacrylamide mini-gel using a Bio-Rad PROTEAN III system and separated at 75 V for 30 min followed by 150 V for 1 h in 1x SDS running buffer (25 mM Tris-HCl pH 6.8, 190 mM glycine, 0.1% (w/v) SDS). For immunoblots, proteins were then transferred to an EtOH-activated polyvinylidene difluoride (PVDF) membrane at 100 V for 1.5 h at 4°C in a wet transfer buffer (25 mM Tris-HCl pH 6.8, 190 mM glycine, 20% EtOH). Membranes were blocked in a 5% skim milk/TBST (20 mM Tris-HCl pH 6.8, 150 mM NaCl, 0.1% Tween-20) solution for 1 h at room temperature, and incubated in the appropriate primary antibody for 12-16 h at 4°C. If secondary antibodies were required, the membrane was washed with TBST prior to secondary incubation. All membranes were washed twice in TBST and once in TBS (20 mM Tris-HCl pH 6.8, 150 mM NaCl) for 10 min prior to enhanced chemiluminescence (ECL) detection of horseradish peroxidase (HRP)-conjugated antibodies. Membranes were incubated with ECL Clarity Substrate (BioRad) and visualized on a ChemiDoc Touch Imaging System (BioRad). Antibodies and titers used: 1:5,000 mouse anti-GFP (Roche 1814460001); 1:10,000 goat anti-mouse-HRP (Sigma A0168); 1:5,000 rabbit anti-His (Cell Signaling 2365); 1:5,000 mouse anti-GST (Sigma SAB4200237); 1:2,000 rabbit anti-p44/42 MAPK (Erk1/2) (Cell Signaling 9102S); 1:10,000 goat anti-rabbit-IgG (Sigma A0545). Depending on the experiment, gels or membranes were stained with Coomassie Brilliant Blue (CBB) R-250 (MP Biomedicals) or SimplyBlue SafeStain (Invitrogen; CBB G-250) to assess protein levels or verify loading.

### Proteomics

Plant growth conditions, protein purification, immunoprecipitation, sample preparation, liquid chromatography followed by tandem mass spectrometry (LC-MS/MS), and data analysis were previously described in full detail (Bender *et al.*, 2017). Proteins identified in immunoaffinity-enriched samples were measured with data dependent method on high resolution LC-MS systems, Orbitrap Fusion (Thermo Fisher Scientific). The acquired spectra were peak-picked and searched by Mascot search engine (Matrix Science Ltd.) to identify the peptide sequences from the search space defined by the background proteome. The peptides were combined into proteins based on the principle of parsimony by the search engine. Resulting proteins were further described by quantitative values based on the number of spectra that identified them. The individual runs were combined in the Scaffold program (Proteome Software Inc.), where the data were evaluated and filtered to contain less than 1% false positives (FDR) and the resulting matrix was exported as a spreadsheet. The matrix of proteins detected in different samples served as the input for an R script for further processing and visualization (**File S1**).

### Statistics

GraphPad Prism 8 or R were used to perform statistical tests on all quantitative data.

## Results

### Identification of CPK28-associated proteins

To identify potential CPK28 interacting partners, we affinity-purified CPK28 C-terminally tagged with yellow fluorescent protein (YFP) from complementing *cpk28-1/35S:CPK28-YFP* transgenic lines (Matschi *et al.*, 2013; Monaghan *et al.*, 2014). We similarly affinity-purified the plasma membrane-localized protein NSL1-YFP from *nsl1-1/35S:NSL1-YFP* (Holmes *et al.*, 2021) and transmembrane protein Lti6B-GFP from Col-0/*35S:Lti6B-GFP* (Cutler *et al.*, 2000) lines to serve as comparative controls. Following immunoprecipitation with anti-GFP microbeads, we performed liquid chromatography followed by tandem mass spectrometry (LC-MS/MS) to identify peptides associated with CPK28, NSL1, or Lti6B. We considered peptides that reliably co-immunoprecipitated with CPK28-YFP across independent trials, but did not co-immunoprecipitate reliably with NSL1-YFP or Lti6B-GFP, as potential CPK28-associated proteins (**Table S2**).

Notably, we identified peptides corresponding to experimentally-validated CPK28-associated proteins including the NADPH oxidase RESPIRATORY OXIDASE HOMOLOG D (RBOHD) (Monaghan *et al.*, 2014) and calmodulin (Bender *et al.*, 2017). We also identified peptides corresponding to other known CPK28-associated proteins, including multiple isoforms of methionine adenosyltransferase (MAT) (Jin *et al.*, 2017), ascorbate peroxidase (APX) (Hu *et al.*, 2021) and glutamine synthase (GS) (Hu *et al.*, 2021); however, peptides for all of these proteins were also observed in the NSL1-YFP and Lti6B-GFP controls (**Table S2**). It is important to consider that context-specific associations between CPK28 and binding partners may not be captured from co-immunoprecipitation-based proteomics reflecting only a single time point during the plant growth cycle. Indeed, we did not recover peptides corresponding to several other experimentally-validated CPK28 binding partners. While we did not identify peptides corresponding to the ARABIDOPSIS TOXICOS EN LEVADURA E3 ubiquitin ligases ATL6 or ATL31, which polyubiquitinate the active form of CPK28 resulting in its proteasomal degradation (Liu *et al.*, 2022, 2023), nor any peptides corresponding to the E3 ligases PUB25 or PUB26, which are phosphorylated and partially activated by CPK28 (Wang *et al.*, 2018), we did identify several components of the ubiquitin-proteasome machinery (**Table S2**). Similarly, although we did not identify peptides corresponding to the RLCK BIK1, which associates with and reciprocally phosphorylates CPK28 (Monaghan *et al.*, 2014; Bredow *et al.*, 2021), we did identify five related RLCKs: BRASSINOSTEROID SIGNALING KINASE 1 (BSK1), CYTOSOLIC ABA RECEPTOR KINASE 7 (CARK7), MAZZA (MAZ/CARK5), PROLINE-RICH EXTENSIN-LIKE KINASE 1 (PERK1) and PERK15 (**Table S2**). In tomato (*Solanum lycopersicum*; Sl), SlCPK28 associates with the phytosulfokine receptor SlPSKR1 (Ding *et al.*, 2022a), and although we did not identify peptides corresponding to AtPSKR1 in our dataset, we did identify twelve other RKs as putative CPK28 binding partners: SUPPRESSOR OF BAK1-INTERACTING KINASE 1 (SOBIR1), BAK1-ASSOCIATING RECEPTOR KINASE 1 (BARK1), LEUCINE-RICH REPEAT RECEPTOR-LIKE KINASE WITH EXTRACELLULAR MALECTIN-LIKE DOMAIN 1 (LMK1), NEMATODE-INDUCED LRR-RLK 2 (NILR2), LYSM RLK1-INTERACTING KINASE 1 (LIK1), MDIS1-INTERACTING RECEPTOR LIKE KINASE 2 (MIK2), FERONIA (FER), MEDOS 1 (MDS1), HERCULES RECEPTOR KINASE 4 (HERK4), WALL-ASSOCIATED KINASE 1 (WAK1), WAK2, and L-TYPE LECTIN RECEPTOR KINASE IV.1 (LECRK-IV.1) (**Table S2**). CPK28 is a multi-functional protein with roles in immune signaling (Monaghan *et al.*, 2014, 2015; Wang *et al.*, 2018), vegetative-to-reproductive stage transition (Matschi *et al.*, 2013, 2015), temperature stress responses (Hu *et al.*, 2021; Ding *et al.*, 2022b), and more (Jin *et al.*, 2017; Ding *et al.*, 2022a). The potential for CPK28 to associate with so many RKs and RLCKs at the plasma membrane may reflect this broad functionality.

### CPK28 associates with subfamily C7 Raf-like protein kinases

We identified 8 unique peptides corresponding to the Raf-like protein kinase MRK1/RAF48 as a putative CPK28-associated protein (**Table 1; Table S2**). To confirm that MRK1 associates with CPK28, we performed split-luciferase complementation assays. In this method, one protein of interest is C-terminally tagged with the N-terminus of firefly luciferase (nLuc) and the other protein of interest is N-terminally tagged with the C-terminus of firefly luciferase (cLuc). If the two proteins associate, they reconstitute luciferase catalytic activity and emit light when provided with the substrate luciferin (Chen *et al.*, 2008). We found that transiently co-expressing CPK28-nLuc and cLuc-MRK1 in *N. benthamiana* reconstitutes the enzymatic function of luciferase, while co-expressing cLuc-MRK1 with another plasma-membrane localized protein, FER-nLuc, does not (**Figure 1A**). MRK1 belongs to the Raf-C7 subfamily and is closely related to four other proteins, sharing 64-65% sequence identity at the amino acid level with RAF26 and RAF39 (78% identical), and CBC1 and CBC2 (Hiyama *et al.*, 2017) (78% identical). Because of their similarity, we were curious if RAF26, RAF39, or CBC1 could also associate with CPK28. Interestingly, we found that co-expressing CPK28-nLuc with cLuc-RAF26, cLuc-RAF39, or cLuc-CBC1 similarly reconstituted luciferase function (**Figure 1B-C; Figure S1**). We conclude that CPK28 is able to associate with C7 Raf-like kinases *in vivo*.

**Figure 1.**
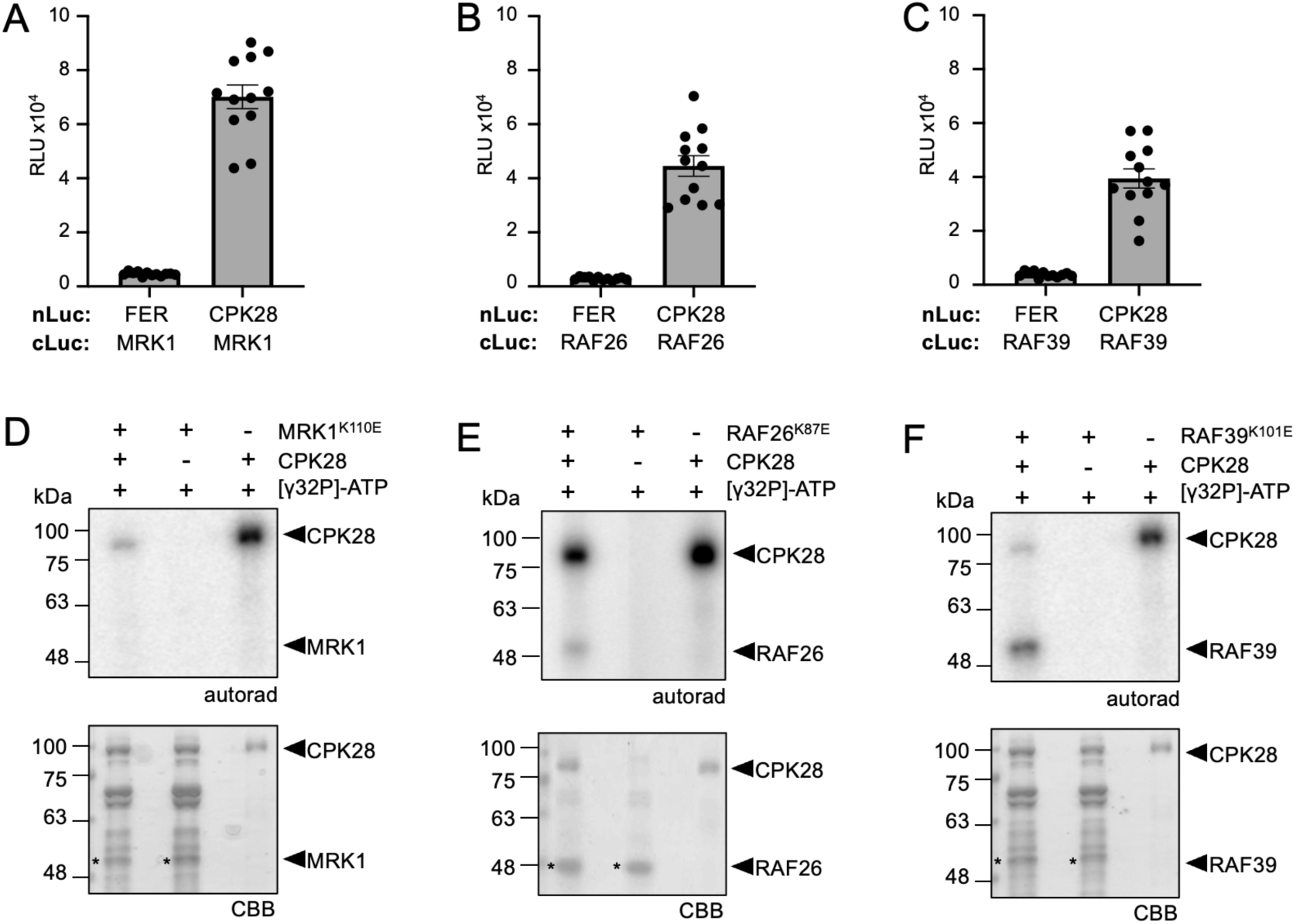
CPK28 associates with C7 Raf-like kinases and phosphorylates RAF26 and RAF39. **(A-C)** Split-luciferase complementation assays with FER-nLuc or CPK28-nLuc and cLuc-MRK1 **(A)**, cLuc-RAF26 **(B)**, and cLuc-RAF39 **(C)**. Total photon counts are plotted as relative light units (RLU) after co-expression of the respective proteins in *N. benthamiana*. Individual values are plotted from a representative experiment (n=12) and are significantly different from each control (Student’s unpaired t-test; *p*<0.0001). These assays were repeated over 4 times each by BD over a 12 month period with similar results; representative data are shown. **(D-F)** *In vitro* kinase assays using His_6_-MBP-CPK28 as the kinase and catalytically-inactive His_6_-MRK1^K110E^ **(D)**, His_6_-RAF26^K87E^ **(E)**, or His_6_-RAF39^K101E^ **(F)** as substrates. Autoradiographs (autorad) indicate incorporation of γP^32^ and protein loading is indicated by post-staining the membranes with Coomassie Brilliant Blue (CBB). Assays were performed more than 3 times each by MGD over a 6 month period with similar results; representative data are shown. Cloning credits are provided in **Table S1**.

**Table 1.**
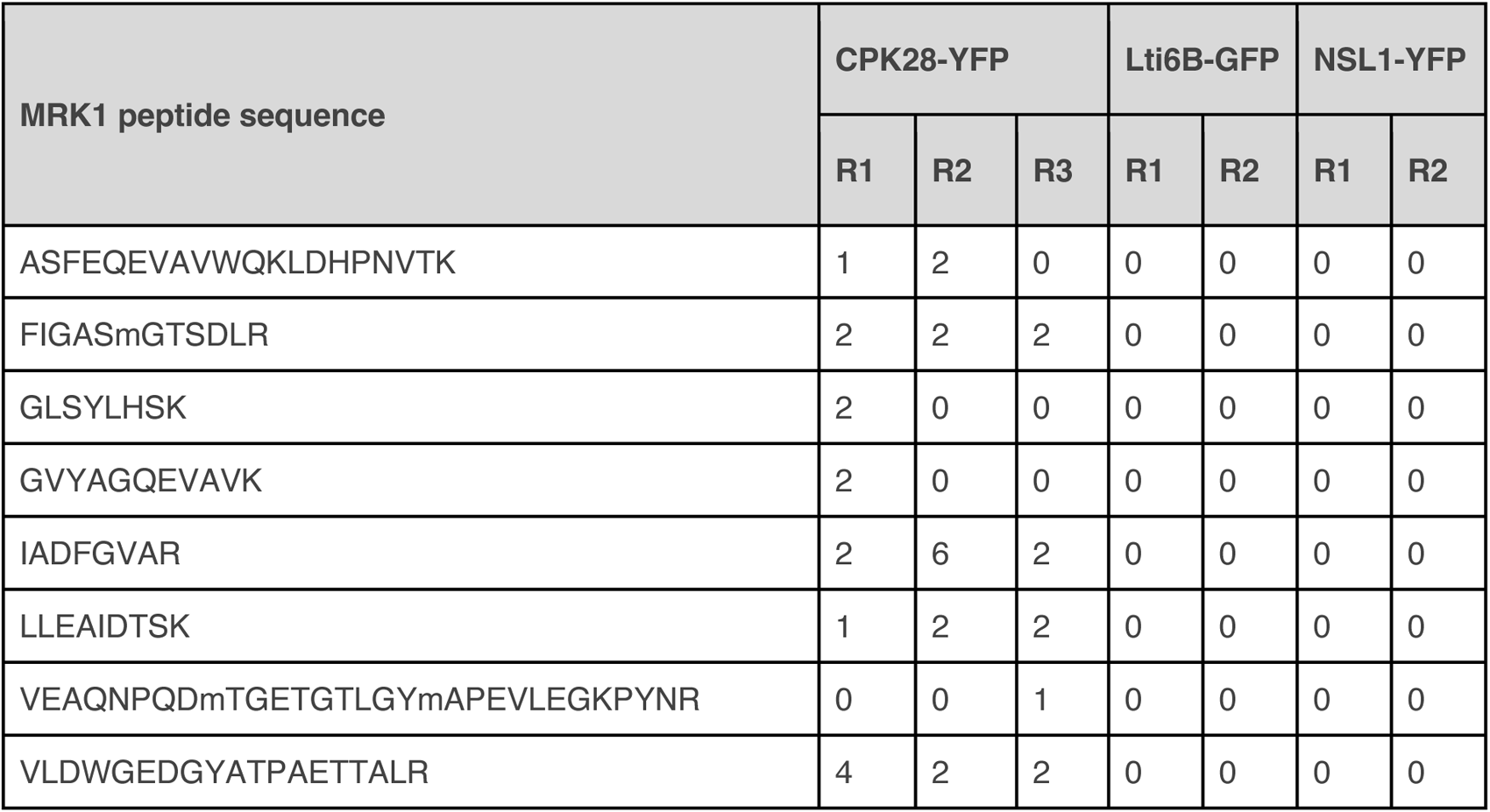
MRK1 peptides identified following affinity-purification of CPK28-YFP. Total spectral counts for MRK1 in each of the biological replicates (R1-R3) for the bait CPK28-YFP and both negative controls Lti6B-GFP and NSL1-GFP. Mascot search files were imported into Scaffold (2.5.1) and filtered with a 1% FDR protein threshold. See **Table S2** for more details.

| MRK1 peptide sequence | CPK28-YFP |  |  | Lti6B-GFP |  | NSL1-YFP |  |
| --- | --- | --- | --- | --- | --- | --- | --- |
|  | R1 | R2 | R3 | R1 | R2 | R1 | R2 |
| ASFEQEVAVWQKLDHPNVTK | 1 | 2 | 0 | 0 | 0 | 0 | 0 |
| FIGASmGTSDLR | 2 | 2 | 2 | 0 | 0 | 0 | 0 |
| GLSYLHSK | 2 | 0 | 0 | 0 | 0 | 0 | 0 |
| GVYAGQEVAVK | 2 | 0 | 0 | 0 | 0 | 0 | 0 |
| IADFGVAR | 2 | 6 | 2 | 0 | 0 | 0 | 0 |
| LLEAIDTSK | 1 | 2 | 2 | 0 | 0 | 0 | 0 |
| VEAQNPQDmTGETGTLGYmAPEVLEGKPYNR | 0 | 0 | 1 | 0 | 0 | 0 | 0 |
| VLDWGEDGYATPAETTALR | 4 | 2 | 2 | 0 | 0 | 0 | 0 |

### CPK28 phosphorylates RAF26 and RAF39

CPK28 displays strong kinase activity both *in vivo* (Matschi *et al.*, 2013; Monaghan *et al.*, 2015) and *in vitro* (Monaghan *et al.*, 2014; Bender *et al.*, 2017). Because they are able to associate, we hypothesized that trans-phosphorylation may occur between CPK28 and C7 Raf-like kinases. As protein kinases have well-defined structures with a high level of conservation, it is possible to predict the location of the ATP-binding lysine in the active site. We therefore generated lysine (K)-to-glutamate (E) variants for MRK1, RAF26, and RAF39 to render them catalytically inactive in order to differentiate auto-from trans-phosphorylation events. We then expressed and purified recombinant MRK1^K110E^, RAF26^K87E^, or RAF39^K101E^ N-terminally tagged with His_6_, as well as CPK28 N-terminally tagged with His_6_ and maltose-binding protein (MBP) from *E. coli* and performed *in vitro* kinase assays using γP^32^-ATP. While we were unable to detect CPK28-mediated phosphorylation of MRK1^K110E^ (**Figure 1D**), CPK28 was able to phosphorylate both RAF26^K87E^ (**Figure 1E**) and RAF39^K101E^ (**Figure 1F**), as indicated by the incorporation of γP^32^. We conclude that while CPK28 can associate with MRK1, RAF26, and RAF39 *in planta*, it is only able to trans-phosphorylate RAF26 and RAF39 *in vitro*, suggesting that CPK28 possesses a high level of specificity for substrate choice.

### MRK1, RAF26, and RAF39 auto-phosphorylate *in vitro*

C7 Raf-like kinases MRK1, RAF26, RAF39, CBC1, and CBC2 all contain the TKL-Pl-4 consensus sequence G-T-x-x-[W/Y]-M-A-P-E in the kinase domain (**Figure 2A**). Alphafold2 (Jumper *et al.*, 2021) predictions suggest that C7 Raf-like kinases adopt typical bilobal protein kinase structures; however, we noted the presence of an extended intrinsically disordered loop between the β4 and β5 sheets in the N-lobe (**Figure 2B**). A multiple sequence alignment of this area in all members of the Arabidopsis MKKK, Raf, and ZIK/WNK (With No Lysine) families revealed that while this loop typically contains 2-3 amino acids, it is uniquely extended to 21-26 residues in the C7-Raf subfamily (**Figure 2B**). Furthermore, this extension contains 3-5 phosphorylatable residues which may confer regulatory functions specific to the C7-Raf subfamily.

**Figure 2.**
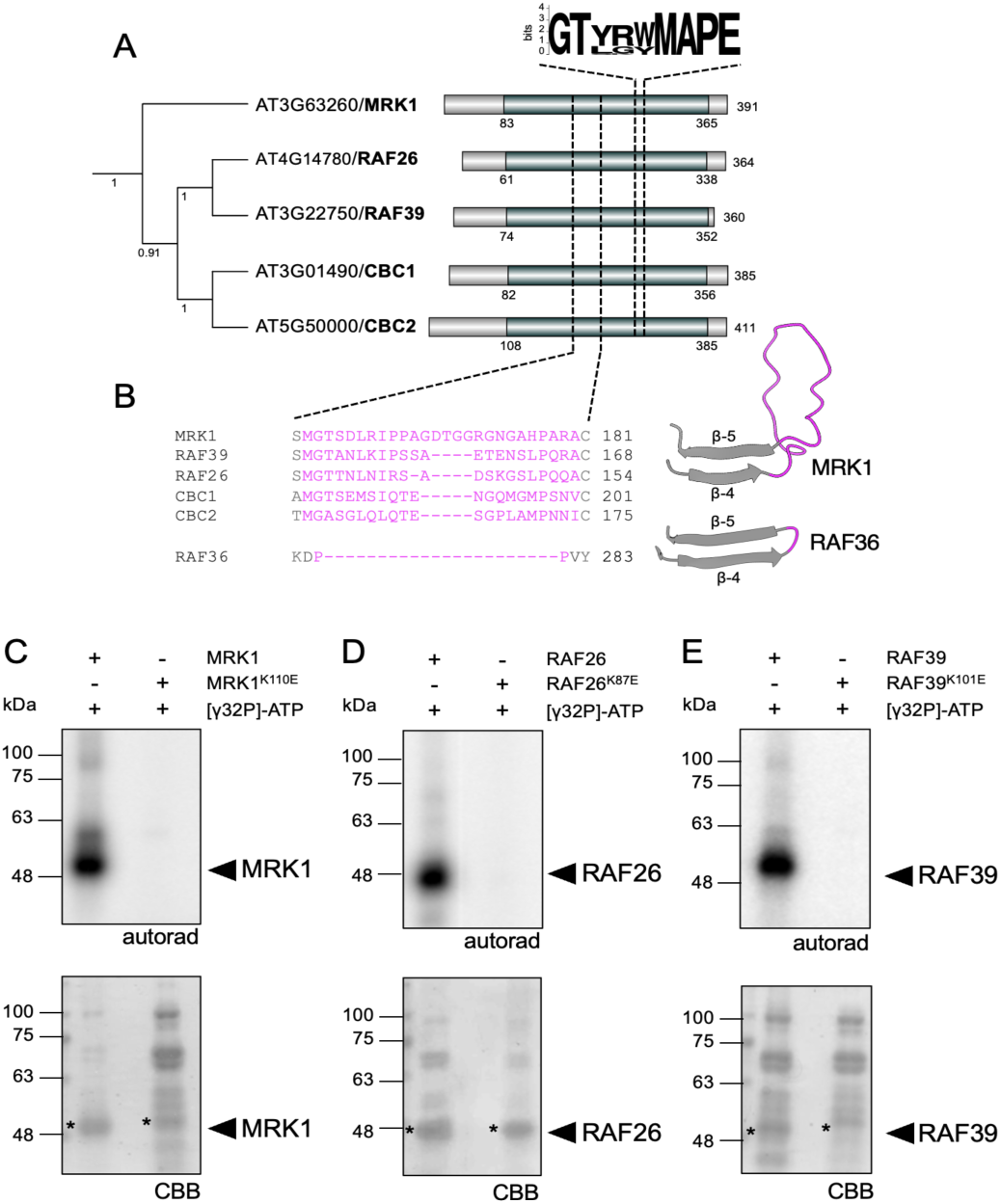
Subfamily C7 Raf-like kinases have a unique extended loop in the N-lobe of the kinase domain and auto-phosphorylate *in vitro*. **(A)** Protein sequences from the subfamily C family of Raf-like kinases were retrieved from The Arabidopsis Information Resource and a multiple sequence alignment was generated using the Muscle algorithm in MEGAX (Kumar *et al.*, 2018). The alignment was used to generate a neighbour-joining tree with 1000 bootstraps; the tree shown here is just the C7 subfamily. The full C-Raf family alignment was used to analyze the consensus signature motif G-T-x-x-[W/Y]-M-A-P-E and visualized here using Weblogo (Crooks *et al.*, 2004). The protein kinase domains are labeled based on the Uniprot database; the protein lengths are indicated on the far right. **(B)** Multiple sequence alignment of the C7 Raf-like kinases compared to RAF36 to illustrate the unique extension identified in C7 Raf-like kinases, which forms an extended disordered loop between the β4 and β5 sheets of the N-lobe (predictions shown for MRK1 and RAF36). Predicted protein structures were downloaded from Alphafold2 (Jumper *et al.*, 2021)and visualized using ChimeraX (Pettersen *et al.*, 2021). **(C-E)** *In vitro* kinase assays indicate that His_6_-MRK1 **(C)**, His_6_-RAF26 **(D)**, and **(E)** His_6_-RAF39 are able to auto-phosphorylate. Each assay included catalytically-inactive variants as controls. Autoradiographs (autorad) indicate incorporation of γP^32^ and protein loading is indicated by post-staining the membranes with Coomassie Brilliant Blue (CBB). JM performed the analysis in A and B; MGD performed the kinase assays in C, D, E at least three times over a 6 month period with similar results and representative data is shown. Cloning credits are provided in **Table S1**.

**Figure 3.**
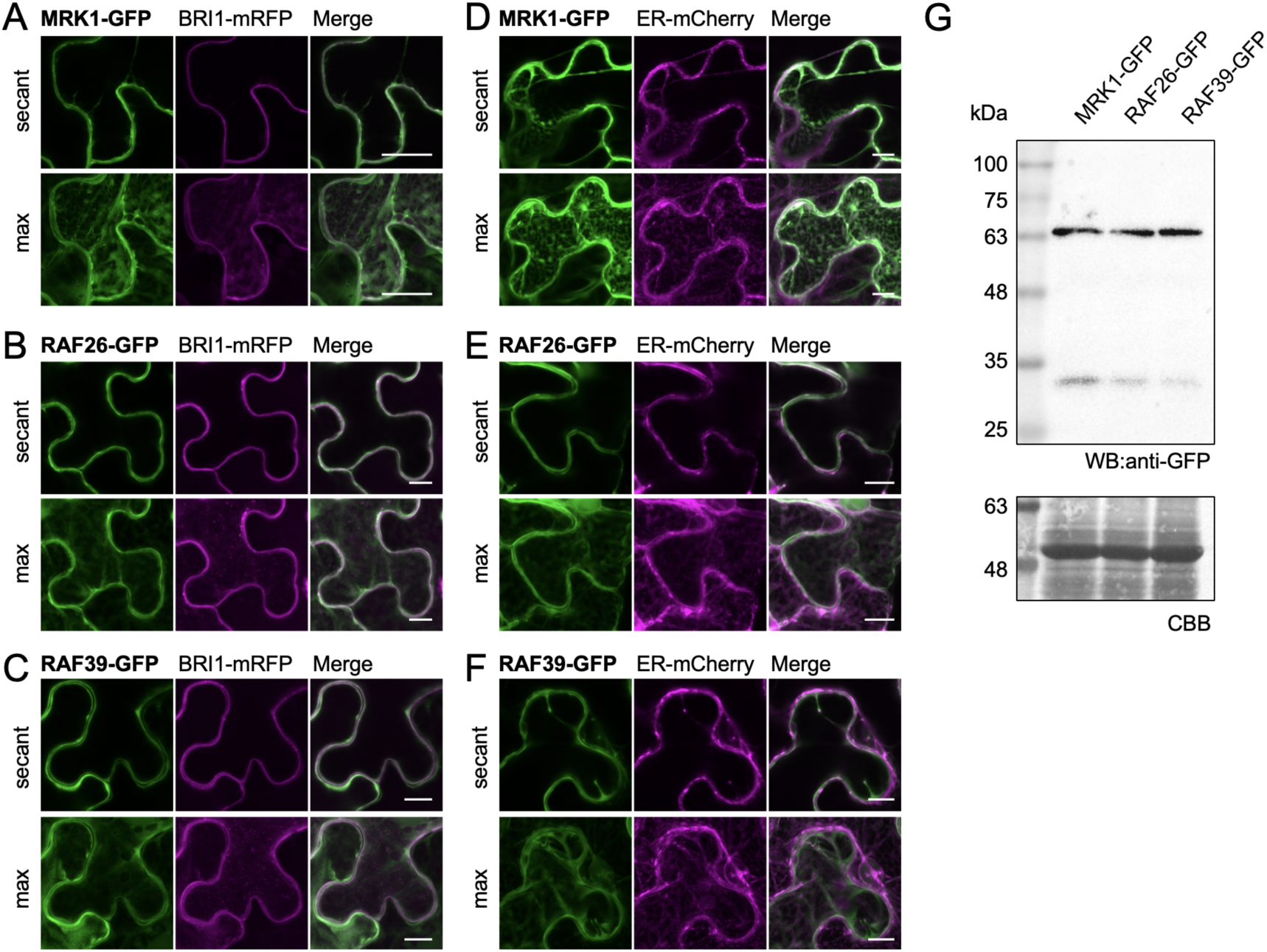
MRK1, RAF26, and RAF39 localize to endomembranes and the cytosol. **(A-F)** Confocal micrographs of MRK1-GFP, RAF26-GFP, and RAF39-GFP co-expressed with either BRI1-mRFP **(A-C)** or ER-mCherry **(D-F)** in *N. benthamiana.* Maximum projections (max) are shown in the lower panels and single-plane sections (secant) are shown in the upper panels. Scale bars are 20 μm (A) or 10 μm (B-F). These assays were repeated 3 times by AR over a 6 month period with similar results. **(G)** MRK1-GFP, RAF26-GFP, and RAF39-GFP were expressed in *N. benthamiana,* proteins extracted and a western blot using anti-GFP antibodies was performed. MRK1-GFP (∼69.6 kDa) RAF26-GFP (∼67.7 kDa), and RAF39-GFP (∼69.7 kDa) migrated to their expected sizes. Coomassie Brilliant Blue (CBB) of RuBisCO indicates loading. This experiment was repeated twice with identical results by MGD. Cloning credits are provided in **Table S1**.

Several MKKKs and Raf-like kinases can auto-phosphorylate *in vitro* (Ma *et al.*, 2022). Indeed, recombinantly purified CBC1 and CBC2 N-terminally tagged with GST are both capable of *in vitro* auto-phosphorylation (Hiyama *et al.*, 2017). To determine if the other C7 Raf-like kinases are similarly capable of auto-phosphorylation, we expressed and purified recombinant MRK1, RAF26, and RAF39 N-terminally tagged with His_6_ from *E. coli* and performed auto-phosphorylation assays *in vitro* using γP^32^-ATP. As controls, we included the catalytically inactive variants His_6_-MRK1^K110E^, His_6_-RAF26^K87E^, and His_6_-RAF39^K101E^ to rule out the possibility of trans-phosphorylation by co-purified proteins. We found that the wild-type variants of MRK1, RAF26, and RAF39 readily incorporated γP^32^, while the catalytically-inactive variants did not, indicating that they possess kinase activity *in vitro* and can auto-phosphorylate (**Figure 2C-E**).

### MRK1, RAF26, and RAF39 localize to the cytosol and endomembranes

Peptides matching MRK1, RAF26, RAF39, CBC1, and CBC2 have been identified in multiple Arabidopsis plasma membrane proteomes (Nelson *et al.*, 2006; Benschop *et al.*, 2007; Niittylä *et al.*, 2007; Marmagne *et al.*, 2007; Mitra *et al.*, 2009; Kamal *et al.*, 2020). Recently, CBC1-GFP and CBC2-GFP were found to localize to the cytosol in Arabidopsis guard cells, but can associate with another Raf-like kinase, HIGH TEMPERATURE 1 (HT1) at the cell periphery in bimolecular fluorescence complementation experiments (Hiyama *et al.*, 2017). To determine their subcellular localization, we cloned MRK1, RAF26, and RAF39 as C-terminal translational fusions with green fluorescent protein (GFP), transiently expressed them in *N. benthamiana,* and visualized cellular fluorescence using confocal microscopy. Co-expression with the plasma membrane marker BRASSINOSTEROID INSENSITIVE 1 (BRI1)-mRFP suggested that pools of MRK1-GFP, RAF26-GFP, and RAF39-GFP localize to the plasma membrane, however these proteins also localize throughout the cytosol (**Figure 4A-C**). We found that MRK1-GFP, RAF26-GFP, and RAF39-GFP co-localized strongly with endomembrane marker ER-mCherry (**Figure 4D-F**). Importantly, all proteins migrated to expected sizes in a western blot (**Figure 4G**). Taken together, these results suggest that MRK1, RAF26, and RAF39 broadly localize throughout the cytosol and the endomembrane system – locations that make it possible to associate with CPK28 at the plasma membrane. Because we were unable to identify classical secretory pathway sorting sequences or endoplasmic reticulum (ER) retention signals in any of these proteins using the signal prediction tools WoLFPSORT (Horton *et al.*, 2007) or SignalP 5.0 (Almagro Armenteros *et al.*, 2019), we hypothesize that MRK1, RAF26, and RAF39 localize to endomembranes via additional binding partners.

**Figure 4.**
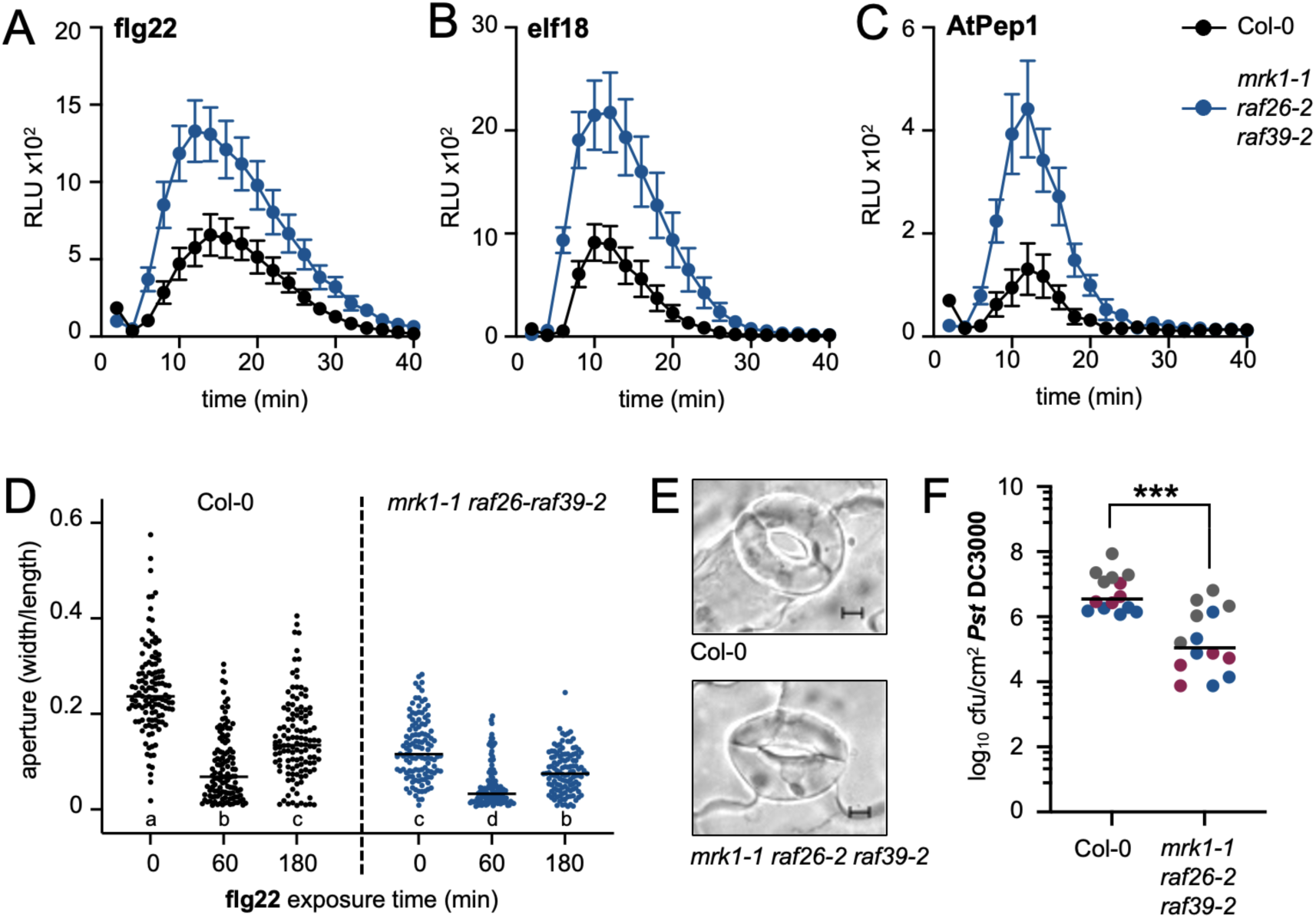
MRK1, RAF26, and RAF39 are genetically redundant regulators of immune homeostasis and stomatal opening. **(A-C)** ROS production measured in relative light units (RLUs) after treatment with 100 nM flg22 **(A)**, 100 nM elf18 **(B)**, or 500 nM AtPep1 **(C)**. Values represent means +/-standard error (n=6-12). Data presented in A was collected by BD; data presented in B and C was collected by MGD. These assays were repeated several times by BD, MGD, EC, and JM over multiple years. **(D)** Stomatal apertures prior to (0 min) and following exposure to 1 μM flg22 (60, 180 min). Individual values are plotted and represent ratios of stomatal width:length. The straight line represents the mean (n=120). Lower case letters indicate statistically significant groups, determined by a one-way ANOVA followed by Tukey’s post-hoc test (*p*<0.005). **(E)** Representative micrographs of stomata prior to flg22 treatment, showing visibly smaller apertures in *mrk1-1 raf26-2 raf39-2* compared to Col-0. Scale bar is 5 μm. Experiments in D and E were repeated 5 times by BD and AR over a 12 month period; representative data collected by BD is shown. **(F)** Growth of *Pseudomonas syringae* pv. *tomato* (*Pst*) isolate DC3000 3 days after spray-inoculation. Data from 3 independent biological replicates are plotted together, denoted by gray, blue, and magenta dots. Values are colony forming units (cfu) per leaf area (cm^2^) from 4-5 samples per genotype (each sample contains 3 leaf discs from 3 different infected plants). The line represents the mean (n=14). Asterisks indicate significantly different groups, determined by a Student’s unpaired t-test (*p*<0.0001). Data was collected by AR over a 12 month period. Credits for genetic crosses and genotyping are provided in **Table S1**.

### MRK1, RAF26, and RAF39 are genetically redundant regulators of immune-triggered ROS

Lacking a humoral system, plants rely on innate and cell-autonomous immune responses to fight against disease. Plant cell membranes contain high-affinity transmembrane pattern recognition receptors (PRRs) that detect highly conserved microbial molecules known as microbe-associated molecular patterns (MAMPs) or endogenous damage-associated molecular patterns (DAMPs). Small peptides known as phytocytokines can also be secreted into the extracellular space, bind PRRs, and potentiate immune signaling (Gust *et al.*, 2017; Segonzac & Monaghan, 2019). In plants, PRRs are typically RKs or RPs. RKs contain a ligand-binding ectodomain, a transmembrane domain, and an intracellular protein kinase domain, allowing them to both detect M/DAMPs and transduce the signal. In contrast to RKs, RPs lack a kinase domain, relying on regulatory RKs to relay the signal (DeFalco & Zipfel, 2021). The largest group of plant PRRs are the leucine-rich repeat (LRR)-containing RKs, which preferentially bind protein-based M/DAMPs. The LRR-RK FLAGELLIN SENSING 2 (FLS2) binds flg22, a 22-amino acid epitope from the N-terminus of bacterial flagellin, while the LRR-RKs EF-Tu RECEPTOR (EFR) and PEP-RECEPTOR 1 and 2 (PEPR1/2) bind the 18-amino acid epitope of elongation factor Tu (elf18) or endogenous peptide AtPep1, respectively (Zipfel *et al.*, 2006; Chinchilla *et al.*, 2007; Yamaguchi *et al.*, 2010; Krol *et al.*, 2010). Both RKs and RPs form heteromeric complexes with regulatory co-receptors at the plasma membrane that typically engage in reciprocal trans-phosphorylation ultimately leading to receptor complex activation and intracellular signaling, including changes in ion flux, defense gene expression, and ROS production (Couto & Zipfel, 2016).

Because of their association with immune regulator CPK28, we hypothesized that C7 Raf-like kinases may function in plant immune signaling. To test if C7 Raf-like kinases are genetically required for plant immune responses, we obtained homozygous insertional mutants in *MRK1 (mrk1-1), RAF26 (raf26-1, raf26-2), RAF39 (raf39-1, raf39-2), CBC1 (cbc1-1, cbc1-2),* and *CBC2 (cbc2-3)* (**Figure S2A**). We noted that leaf and rosette morphology in all mutants was comparable to wild-type Col-0 plants grown over multiple years in controlled environment chambers (**Figure S2B**), although we did note slightly smaller growth in the *cbc1-1 cbc2-3* mutant as previously reported (Hiyama *et al.*, 2017). Following the detection of immunogenic peptides by PRRs, RLCKs and CDPKs phosphorylate and activate the NADPH oxidase RBOHD, which catalyzes the production of a burst of apoplastic ROS within minutes (Yu *et al.*, 2017). We found that the flg22-induced ROS burst was not affected in *mrk1-1, raf26-1, raf26-2, raf39-1, raf39-2, cbc1-1, cbc1-2,* or *cbc2-3* single mutants (3/3 independent biological replicates; **Figure S3A-D**). As genetic redundancy was previously shown between *CBC1* and *CBC2* in blue light-mediated stomatal opening (Hiyama *et al.*, 2017), we also generated *cbc1-1 cbc2-3* and *raf26-2 raf39-2* double mutants. The flg22-induced ROS burst was not affected in the *cbc1-1 cbc2-3* double mutant (12/13 biological replicates; **Figure S3E**), nor in the *raf26-2 raf39-2* double mutants (8/10 biological replicates; **Figure S3F**). We next generated a *mrk1-1 raf26-2 raf39-2* triple mutant, as well as *mrk1-1 raf26-2* and *mrk1-1 raf39-2* double mutants. We consistently observed enhanced flg22-triggered ROS in both the *mrk1-1 raf26-2* and *mrk1-1 raf39-2* double mutants (14/15 and 14/16 biological replicates, respectively), as well as the *mrk1-1 raf26-2 raf39-2* triple mutant (13/14 replicates; **Figure 4A, Figure S3G-H**). This response is not specific to flg22, as we also observed enhanced elf18-and AtPep1-triggered ROS production in *mrk1-1 raf26-2 raf39-2* (4/4 biological replicates; **Figure 4B,C**). Importantly, we confirmed that these alleles result in lower expression of their target genes (**Figure S2C**). To test if enhanced ROS confers enhanced disease resistance in *mrk1-1 raf26-2 raf39-2*, we infected plants with the virulent bacterial pathogen *Pseudomonas syringae* pv. *tomato* (*Pst*) DC3000 and counted *in planta* bacterial growth 3 days after syringe-infiltration. We observed similar bacterial growth in Col-0 and *mrk1-1 raf26-2 raf39-2* (4/5 biological replicates; **Figure S3I**). Although we did not generate a *mrk1-1 cbc1-1 cbc2-3* triple mutant, our results suggest that *MRK1* plays a key role in regulating immune-triggered ROS, sharing unequal genetic redundancy with at least *RAF26* and *RAF39*.

### C7 Raf-like kinases regulate light-induced stomatal opening that correlates with enhanced resistance to a bacterial pathogen

The production of ROS is thought to provide direct antimicrobial activity in the apoplast, and also acts as a signaling molecule (Melotto *et al.*, 2017). In guard cells, immune-triggered ROS production has been linked to stomatal closure. While stomatal pores play a critical role in controlling gas exchange for photosynthesis, open stomata can be seized as a point of entry for microbial pathogens; stomatal closure thus restricts access (Melotto *et al.*, 2017). C7 Raf-like kinases are expressed broadly throughout plant tissues, including in guard cells (Hayashi *et al.*, 2017). CBC1 and CBC2 are particularly strongly expressed in guard cells and have been shown to function redundantly in blue light and CO_2_-mediated stomatal opening (Hiyama *et al.*, 2017; Takahashi *et al.*, 2022). With this in mind, we were interested to assess if MRK1, RAF26, or RAF39 similarly inhibit stomatal opening. To test this, we first confirmed altered stomatal aperture in the *cbc1-1 cbc2-3* double mutant under bright light compared to Col-0 (**Figure S4A**). Similar to *cbc1-1 cbc2-3,* light-induced stomatal opening was impaired in *raf26-2 raf39-2, mrk1-1 raf26-2,* and *mrk1-1 raf39-2* double mutants (**Figure S4B-D**), as well as *mrk1-1 raf26-2 raf39-2* triple mutants (**Figure 4D-E**), as stomatal apertures were much smaller than in Col-0. We did not observe any differences in stomatal apertures between Col-0 and the *mrk1-1, raf26-2,* or *raf39-2* single mutants (**Figure S4E**). These results suggest that MRK1, RAF26, and RAF39 function redundantly in light-induced stomatal opening.

During an immune response, stomata remain closed for some time but will reopen after the threat has passed. Because of this, we were interested to assess if immune-triggered stomatal closure and re-opening is regulated by the C7 Raf-like kinases. We thus treated plants with flg22 and measured stomatal apertures after 1h and 3h, to reflect the ‘closed’ and ‘reopening’ states in Col-0. In all the double mutants, we observed strong flg22-induced stomatal closure (**Figure S4A-D**). Interestingly, stomata were closed more ‘tightly’ in the triple *mrk1-1 raf26-2 raf39-2* mutant than in Col-0 (3/3 replicates; **Figure 4D**). These data are congruent with previous work that indicated tighter stomatal closure in *cbc1 cbc2* mutants in response to abscisic acid (ABA) (Hiyama *et al.*, 2017), and together support the model that C7 Raf kinases promote stomatal opening by derepressing stomatal closure. When we measured apertures after 3h of exposure to flg22, we observed partial re-opening in Col-0 as well as the double and triple mutants (**Figure 4D, Figure S4A-D**), suggesting that additional components regulate stomatal re-opening following immune-mediated closure.

We reasoned that smaller stomatal apertures capable of closing very tightly in response to an immune trigger might restrict pathogen entry to plant tissue. We therefore spray-inoculated plants with *Pst* DC3000 to better mimic a natural infection and assessed *in planta* growth after three days. Here, we found that bacterial growth was reduced ∼10-fold in *mrk1-1 raf26-2 raf39-2* compared to Col-0 plants (3/3 biological replicates; **Figure 4F**). Interestingly, we also observed reduced bacterial growth in *cbc1-1 cbc2-3* mutants when spray-inoculated with *Pst* DC3000 (2/3 biological replicates; **Figure S4F**). This suggests that the smaller stomatal aperture observed in both *mrk1-1 raf26-2 raf39-2* and *cbc1-1 cbc2-3* is capable of providing enhanced resistance to *Pst* DC3000.

### MRK1, RAF26, and RAF39 do not trans-phosphorylate MKKs *in vitro*

The phosphorylation and activation of MAPKs occurs within minutes of PRR activation and in parallel with the apoplastic ROS burst (Yu *et al.*, 2017). In Arabidopsis, at least two MAPK cascades are activated following MAMP perception, consisting of MAPKKK5-MKK4/MKK5-MPK3/6 (Asai *et al.*, 2002; Yamada *et al.*, 2016; Bi *et al.*, 2018) or MEKK1-MKK1/MKK2-MPK4 (Ichimura *et al.*, 2002, 2006; Nakagami *et al.*, 2006; Suarez-Rodriguez *et al.*, 2007; Gao *et al.*, 2008). MPK4 and MPK3/6 have diverse targets including WRKY transcription factors that drive expression of immune-related genes and contribute to genetic reprogramming of the cell to combat infection (Mao *et al.*, 2011; Guan *et al.*, 2014). Because C7 Raf-like kinases are predicted to function as MKKKs, and since *mrk1-1 raf26-2 raf39-2* mutants displayed enhanced immune-triggered ROS, we were interested to test if MRK1, RAF26, and RAF39 are involved in immune-triggered MAPK activation. We thus assessed the phosphorylation status of MPK6, MPK3 and MPK4/MPK11 in Col-0 compared to the *mrk1-1 raf26-2 raf39-2* triple mutants following flg22 perception. Our results indicate that MAPK activation occurs similarly in Col-0, *mrk1-1 raf26-2 raf39-2,* and *cbc1-1 cbc2-3* mutants (**Figure S5A-B**).

Although phylogenetically considered a subfamily of MKKKs, it is unclear if Raf-like kinases function biochemically as kinases that phosphorylate MKKs in a MAPK cascade. Both the Raf-like and ZIK/WNK subfamilies are divergent from canonical MKKKs, and neither cluster well with metazoan MKKK, Raf, or MLK proteins (**Figure 5A**) (Champion *et al.*, 2004). To clarify if MRK1, RAF26, and RAF39 can function as MKKKs, we tested if they can trans-phosphorylate MKKs *in vitro.* There are 10 MKKs encoded in Arabidopsis that cluster into four subfamilies: subfamily A contains MKK1, MKK2, and MKK6; subfamily B contains MKK3; subfamily C contains MKK4 and MKK5; and subfamily D contains MKK7, MKK8, MKK9, and MKK10 (Jiang & Chu, 2018). We cloned and purified all 10 MKK proteins as catalytically inactive variants (replacing the ATP-binding lysine with glutamate), N-terminally tagged with GST. Kinase assays using γP^32^-ATP indicate that none of MRK1, RAF26, or RAF39 are able to trans-phosphorylate any of the 10 Arabidopsis MKKs *in vitro* (**Figure 5B-D**). This suggests that they do not function biochemically as MKKKs in MAPK cascades.

**Figure 5.**
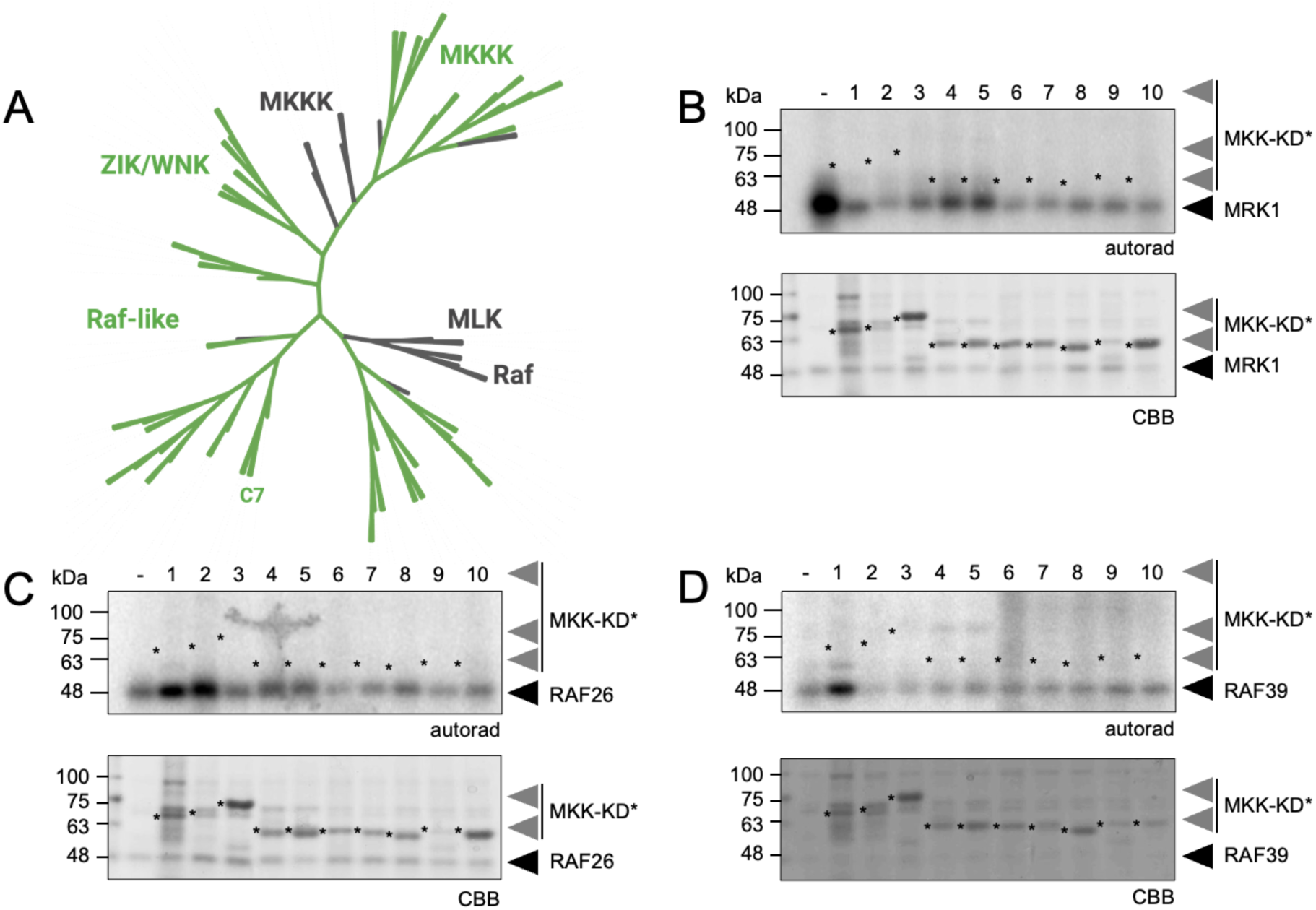
MRK1, RAF26, and RAF39 do not trans-phosphorylate MKKs. **(A)** An unrooted phylogenetic tree of the Arabidopsis MKKK, ZIK/WNK, and Raf-like subfamilies (green) together with human MKKK, MLK, and Raf kinases (gray). Subfamily C7 Raf-like kinases are indicated. A multiple sequence alignment using the full-length sequences of all proteins in the subfamilies was performed using Clustal Omega and the resulting neighbour-joining phylogenetic tree was visualized using iTOL (Letunic & Bork, 2021); subfamilies are collapsed at the ends of nodes. Analysis performed by JM. **(B-D)** *In vitro* kinase assays indicate that His_6_-MRK1 **(B)**, His_6_-RAF26 **(C)**, and **(D)** His_6_-RAF39 are unable to trans-phosphorylate any of the 10 Arabidopsis MKKs N-terminally tagged with GST. Catalytically inactive MKK variants were used and are numbered as 1-10 for MKK1^K97E^, MKK2^K108E^, MKK3^K112E/K113E^, MKK4^K108E^, MKK5^K99E^, MKK6^K99E^, MKK7^K74E^, MKK8^K82E/K83E^, MKK9^K76E^, MKK10^K77E^, and are indicated by asterisks. Autoradiographs (autorad) indicate incorporation of γP^32^ and protein loading is indicated by post-staining the membranes with Coomassie Brilliant Blue (CBB). MGD and TD performed the assays three times over a 3 month period with similar results and representative data is shown. Cloning credits are provided in **Table S1**.

## Discussion

Raf-like kinases are a plant-specific family with documented roles in ethylene signaling, osmotic stress, stomatal movement, and immunity (Fàbregas *et al.*, 2020; González-Coronel *et al.*, 2021; Ma *et al.*, 2022). Here, we focus on subfamily C7 Raf-like kinases and provide evidence that they function in the regulation of stomatal aperture and immune signaling. Previous studies have described the redundant roles of CBC1 and CBC2 in stomatal opening (Hiyama *et al.*, 2017; Hayashi *et al.*, 2020; Takahashi *et al.*, 2022), and we demonstrate similar function for the remaining C7 subfamily members MRK1, RAF26, and RAF39. Stomatal pores are formed between two guard cells that allow gas exchange and water transpiration to optimize plant growth, but can also be co-opted by pathogens to gain entry to plant tissues. The aperture of stomatal pores can adopt ‘open’ or ‘closed’ conformations, depending on environmental conditions that include both abiotic and biotic factors. For example, stomata adopt an open conformation under bright light or when levels of CO2 are limiting, thus driving photosynthesis. Conversely, stomata adopt a closed conformation in response to stress signals such as an increase in ABA or cytosolic Ca^2+^, or when levels of CO_2_ are sufficient (Shimazaki *et al.*, 2007; Melotto *et al.*, 2017). While there are pathway-specific signaling mechanisms in place, opening and closing of stomata is ultimately controlled by changes in water potential that affect turgor pressure and membrane polarization/depolarization in guard cells.

In the presence of blue light, activated PHOTOTROPIN 1 and 2 (PHOT1/2) receptors facilitate H^+^-ATPase-mediated plasma membrane hyperpolarization, which results in stomatal opening and increased gas exchange at the stomatal pore (Kinoshita *et al.*, 2001). Anion channels such as SLOW ANION CHANNEL ASSOCIATED 1 (SLAC1) are deactivated following blue light perception to inhibit membrane depolarization (Inoue & Kinoshita, 2017). In the presence of high intracellular CO_2_, the SnRK protein kinase OPEN STOMATA 1 (OST1) activates SLAC1 to trigger anion efflux and ultimately cause stomatal closure. To increase carbon uptake under low CO_2_, the subfamily C5 Raf-like kinase HIGH LEAF TEMPERATURE 1 (HT1) inhibits OST1 activation and facilitates SLAC1 inactivation (Tian *et al.*, 2015), which in turn enhances water uptake in guard cells and results in stomatal opening. Thus, exposure to blue light or high levels of CO_2_ results in stomatal opening in wild-type plants. However, these responses are defective in *cbc1 cbc2* double mutants, where stomata remain closed (Hiyama *et al.*, 2017; Takahashi *et al.*, 2022). Genetic, biochemical, and electrophysiological assays indicate that this phenotype is due to a break in the signaling pathway that enables blue light-induced inhibition of S-type anion channels such as SLAC1 (Hiyama *et al.*, 2017). HT1 activates CBC1 by phosphorylation on several sites, including critical residues in the activation loop (Hiyama *et al.*, 2017; Takahashi *et al.*, 2022). In addition, CBC1 can be phosphorylated by PHOT1 *in vitro* and is rapidly phosphorylated in response to blue light *in vivo* (Hiyama *et al.*, 2017), suggesting that CBC1/2 integrate signals from both blue light and CO_2_ pathways. Here, we show that *mrk1-1 raf26-2 raf39-2* mutants display smaller stomatal aperture similar to *cbc1-1 cbc2-3,* suggesting that all C7 Raf-like kinases participate in stomatal opening. While several phosphorylation sites on MRK1, RAF39, CBC1, and CBC2 have been curated from shotgun phosphoproteomics studies (Wang *et al.*, 2013a,b; Hoehenwarter *et al.*, 2013; Wu *et al.*, 2013; Roitinger *et al.*, 2015; Marondedze *et al.*, 2016; Nukarinen *et al.*, 2016; Bhaskara *et al.*, 2017; Al-Momani *et al.*, 2018; Song *et al.*, 2018) as well as targeted studies (Hiyama *et al.*, 2017; Takahashi *et al.*, 2022), functional roles have so far only been assigned for Ser43 and Ser45 located at the N-terminus of CBC1 (Hiyama *et al.*, 2017). Notably, the majority of phosphosites on MRK1, RAF39, and CBC2 map to their N-termini in areas of low sequence conservation and low intrinsic order (**Figure S6**). Other phosphosites map to areas well known to be involved in kinase activation, including in the Gly-rich and activation loops (**Figure S6**). It will be of interest to assess the functional role of N-terminal phosphorylation on C7 Raf-like kinases, as these are likely to represent areas of isoform-specific regulation.

Here we show that CPK28 associates with C7 Raf-like kinases *in vivo* and is able to phosphorylate RAF26 and RAF39 *in vitro*. Intriguingly, despite high sequence identity with RAF26 and RAF39 (78%), we could not detect CPK28-mediated phosphorylation of MRK1. While it remains possible that CPK28 may phosphorylate MRK1 *in vivo,* these results could reflect different regulatory mechanisms between highly similar proteins. We scrutinized a multiple sequence alignment comparing MRK1, RAF26, and RAF39 to identify differences that could explain these observations. We identified only two areas of sequence divergence in phosphorylatable residues (Ser, Thr, or Tyr) between MRK1 and RAF26 or RAF39. One area is in the C7-specific intrinsically disordered loop connecting the β4 and β5 sheets in the N-lobe (**Figure 2B; Figure S6**), where MRK1 has three phosphorylatable residues while RAF26 and RAF39 each have five, and the other constitutes a 16-amino acid α-helix close to the C-terminal end of the protein, where MRK1 lacks phosphorylatable residues and RAF26 and RAF39 each contain two (**Figure S6**). Mapping CPK28-mediated phosphosites on RAF26 and RAF39 will be of interest, as will testing if CPK28 can phosphorylate CBC1 and CBC2.

Publicly-available gene expression data indicates that *CPK28* and all C7 Raf-like genes are expressed in guard cells (Yang *et al.*, 2008), but *CBC1* and *CBC2* are the most highly expressed (Yang *et al.*, 2008; Hiyama *et al.*, 2017). A role for CPK28 in stomatal aperture has not yet been described, and previous work found no differences in flg22-induced stomatal closure in two *CPK28-OE* lines compared to wild-type plants (Monaghan *et al.*, 2014). Here, we show that both *cbc1-1 cbc2-3* and *mrk1-1 raf26-2 raf39-2* mutants are more resistant to spray-inoculation of the bacterial pathogen *Pst* DC3000, which we consider may be a consequence of their smaller stomatal aperture. In addition, we found that while immune-triggered ROS was unchanged in *cbc1-1 cbc2-3* mutants, the *mrk1-1 raf26-2 raf39-2* mutants displayed enhanced ROS which suggests both unique and overlapping functions within this gene family. Neither *cbc1-1 cbc2-3* nor *mrk1-1 raf26-2 raf39-2* displayed differences in flg22-induced MAPK activation, which occurs in parallel to immune-triggered ROS. Interestingly, viral-induced gene silencing of the wheat (*Triticum aestivum*) ortholog of RAF39, *Ta*Raf46, similarly results in enhanced ROS accumulation, defense gene expression, and protection against the rust stripe pathogen *Puccinia striiformis* f. sp. *tritici* (*Pst*) isolates CYR23 and CYR31 (Wan *et al.*, 2022). Conversely, overexpression of *Ta*Raf46 results in a loss of immune responses and enhanced susceptibility to *Pst* CYR23 (Wan *et al.*, 2022). In addition, viral-induced gene silencing of the cotton ortholog of RAF39, *Gh*MAP3K65, similarly results in enhanced defense gene expression and resistance to both the fungal pathogen *Rhizoctonia solani* and the bacterial pathogen *Ralstonia solanacerum* (Zhai *et al.*, 2017). Although the functional relationship between CPK28 and MRK1, RAF26, and RAF39 is yet to be determined, it would be interesting to know if orthologs of CPK28 phosphorylate RAF39 in wheat and cotton.

To counteract immune responses and enable disease, pathogens secrete effector proteins that target key components of the immune system, including many protein kinases. In resistant plants, pathogen effectors are detected by intracellular nucleotide-binding LRR receptors (NLRs) that trigger localized programmed cell death when activated (El Kasmi, 2021). Interestingly, the *N. benthamiana* ortholog of RAF39 was identified in an *in planta* biotin ligase labeling assay as a protein in close proximity to the *Pst* DC3000 effector AvrPto at the plasma membrane (Conlan *et al.*, 2018). Although a direct protein:protein association between AvrPto and *Nb*RAF39 was not confirmed, this raised the possibility that C7 Raf-like kinases may be recruited or targeted by pathogen effectors. Recently, Pst27791, a serine-rich effector protein from the stripe rust pathogen *Puccinia striiformis* f. sp. *tritici* isolate CYR23 (*Pst* CYR23) was shown to interact with and stabilize the accumulation of *Ta*Raf46 when heterologously expressed in *N. benthamiana* (Wan *et al.*, 2022). Transgenic overexpression of Pst27791 in wheat results in enhanced susceptibility to *Pst* CYR23 only when *Ta*Raf46 is expressed, suggesting that Pst27791 requires *Ta*Raf46 for its virulence (Wan *et al.*, 2022). In Arabidopsis, MRK1 is ubiquitinated on residue K342 (Grubb *et al.*, 2021) and its protein abundance decreases by 50% following flg22 treatment (Benschop *et al.*, 2007), which could reflect a derepression mechanism to enable immune signaling. In this scenario, effector-mediated stabilization of C7 Raf-like kinases could result in sustained repression of immune signaling to further pathogen spread. All of this evidence supports a role for C7 Raf-like kinases as regulators of stomatal aperture and immune homeostasis in multiple plant species, and may therefore be of interest to breeders.

Although Raf-like kinases are considered a subfamily of MKKKs, their *bona fide* role as MKKKs has been debated (Champion *et al.*, 2004; Ma *et al.*, 2022). Canonical MKKKs phosphorylate MKKs at specific Ser/Thr residues located within a conserved S/T-X_3-5_-S/T motif in the activation loop, which activates MKKs and allows them to phosphorylate MAPK targets (Rodriguez *et al.*, 2010). Thus, to behave as a canonical MKKK in a MAPK cascade, a kinase would need to phosphorylate this consensus motif in an MKK. Some Raf-like kinases can phosphorylate MKKs, but there is limited evidence that this phosphorylation occurs within the S/T-X_3-5_-S/T motif. Recently, the subfamily C1 Raf-like kinase RAF27 (also known as BLUE LIGHT-DEPENDENT H+-ATPASE PHOSPHORYLATION; BHP, or INTEGRIN-LIKE KINASE 5; ILK5) was shown to associate with and phosphorylate MKK5, and that mutation of Thr215 and Ser221 in the S/T-X_3-5_-S/T motif to non-phosphorylatable Ala residues reduced trans-phosphorylation by RAF27/BHP/ILK5 (Kim *et al.*, 2023). This suggests that RAF27/BHP/ILK5 phosphorylates MKK5 at the consensus motif as well as at other sites. Something similar was demonstrated for the subfamily B3 Raf-like kinase MKKK δ-1 (MKD1), which can trans-phosphorylate both MKK1 and MKK5 *in vitro* (Asano *et al.*, 2020). Mass spectrometry analysis indicated that while MKD1 phosphorylates MKK5 at Thr215 and Ser221 (within the S/T-X_3-5_-S/T motif), it additionally phosphorylates MKK5 at Thr83 and MKK1 at Ser46 – N-terminal residues that are not found within the activation loop consensus motif (Asano *et al.*, 2020). In rice, the subfamily C2 Raf-like kinase OsILA1 phosphorylates OsMKK4 on multiple N-terminal residues including the key site Thr34, and not in the consensus motif (Chen *et al.*, 2021). Additional evidence that Raf-like kinases are atypical MKKKs comes from studies indicating that some can phosphorylate substrates that are not MKKs. For example, the subfamily B3 Raf-like kinase CONSTITUTIVE TRIPLE RESPONSE 1 (CTR1), a well-known kinase involved in ethylene signaling, phosphorylates ETHYLENE INSENSITIVE 2 (EIN2) at multiple residues (Ju *et al.*, 2012). In addition, several other subfamily B Raf-like kinases phosphorylate members of the sucrose nonfermenting-1-related protein kinase (SnRK) family in osmotic stress signaling (Saruhashi *et al.*, 2015; Takahashi *et al.*, 2020; Lin *et al.*, 2020; Soma *et al.*, 2020; Katsuta *et al.*, 2020; Fàbregas *et al.*, 2020), and the C5 Raf-like kinase HT1 phosphorylates multiple sites on CBC1 including Thr256 and Ser280 in the activation loop (Hiyama *et al.*, 2017; Takahashi *et al.*, 2022). The B3 Raf-like kinase ENHANCED DISEASE SUSCEPTIBILITY 1 (EDR1) negatively regulates immune signaling (Frye *et al.*, 2001; Ma *et al.*, 2022) and has been shown to associate with MKK4 and MKK5 (Zhao *et al.*, 2014). Interestingly, *edr1* mutants accumulate less MKK4/MKK5 and MPK6/MPK3 proteins (Zhao *et al.*, 2014), and EDR1 associates with E3 ligases KEEP ON GOING (KEG) (Wawrzynska *et al.*, 2008; Gu & Innes, 2011) and ATL1 (Serrano *et al.*, 2014). While it is unknown if any of these proteins are EDR1 substrates, KEG ubiquitinates MKK4/MKK5 resulting in their proteasomal turnover (Gao *et al.*, 2021), suggesting that EDR1 regulates MKK accumulation via modulation of E3 ligases. The rice ortholog of EDR1 also negatively regulates immunity (Kim *et al.*, 2003; Shen *et al.*, 2011) and associates with but does not phosphorylate OsMKK10.2 (Ma *et al.*, 2021). All together, these data suggest that EDR1 acts as a noncanonical MKKK in both rice and Arabidopsis. Notably, even some MEKK-like MKKKs play noncanonical roles in signaling pathways. For example, MKKK7 is differentially phosphorylated in response to flg22 and attenuates flg22-induced immune signaling including the activation of MPKs (Mithoe *et al.*, 2016). Thus, it seems that the expansion of the MKKK family in plants has allowed for the evolution of novel functions. While it remains possible that certain Raf-like kinases may operate as canonical MKKKs, it is evident that some Raf-like kinases accept alternative substrate proteins. Our finding that MRK1, RAF26, and RAF39 cannot phosphorylate any of the 10 Arabidopsis MKKs *in vitro* suggests that they likely do not function as canonical MKKKs *in vivo*. An important next step will be to identify biologically relevant substrates for C7 Raf-like kinases, of which currently none are known.

## Acknowledgements

We acknowledge the importance of diversity, equity and inclusion in the sciences and thank all members of the Monaghan Lab for their commitment to fostering a welcoming and collaborative research environment. Queen’s University is situated on the territory of the Haudenosaunee and Anishinaabek and we are grateful to live, work, and play on these lands. We are grateful to all members of our labs, past and present, for engaging discussions over the course of this project, and for reviewing our manuscript before submission. We thank Madison Giroux for assistance with preliminary split-luciferase complementation assays; Saied Mobini for managing the Queen’s University Phytotron Facility; and Tony Papanicolaou for managing the Microscopy Facility in the Department of Biology.

## Competing Interests

None declared.

## Author Contributions

JM and KRS designed the project. MGD, BD, AR, KRS, TM, TD, EC and JM generated materials, performed experiments, and analyzed results. JS and PD processed and analyzed CPK28-associated proteins identified by proteomics, supervised by FM, JM, and CZ. Individual credits are included wherever possible in the figure captions and table legends. JM guided the work, secured funding, and wrote the paper with input from all authors.

## Data Availability

Any materials described in this article will be made freely available upon request. The person responsible for sharing materials is the author of correspondence. Proteomics data has been deposited to the ProteomeXchange Consortium via the PRIDE (Perez-Riverol *et al.*, 2019) partner repository.

## Funding

This work was funded by the following grants awarded to JM: Canadian Natural Sciences and Engineering Research Council of Canada (NSERC) Discovery and Discovery Accelerator Programs [grant numbers RGPIN-2016-04787 and RGPAS-492902-2016], the Canada Research Chair (CRC) Program [JM is CRC-II in Plant Immunology], the Ontario Ministry of Colleges and Universities Early Researcher Award Program [grant number ER21-16-100], and the UK Biotechnology and Biological Sciences Research Council (BBSRC) Anniversary Future Leaders Fellowship Program 2015. Additional funding for proteomics work was provided by core funding from the Gatsby Charitable Foundation for The Sainsbury Laboratory in Norwich, UK. MGD was supported by a Research Internship Abroad fellowship (BEPE) from the São Paulo Research Foundation (FAPESP) [grant number 2021/06835-3]. BD was supported by the Queen’s University Summer Work Experience Program (SWEP 2022) and the Queen’s University Faculty of Arts and Science Undergraduate Research Fund (ASURF 2023). KRS was supported by an NSERC Undergraduate Summer Research Award (USRA 2017), NSERC Canada Graduate Scholarship for MSc students (CGS-M 2017-2018) and an Ontario Graduate Scholarship (OGS 2018-2019).

## Supporting Information

### Supplemental Figures

**Figure S1.**
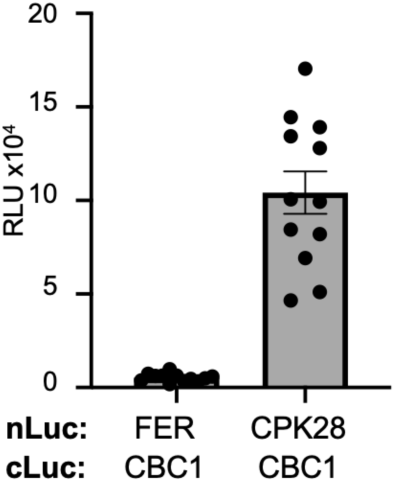
CPK28 associates with CBC1. Split-luciferase complementation assays with FER-nLuc or CPK28-nLuc and cLuc-CBC1. Total photon counts are plotted as relative light units (RLU) after co-expression of the respective proteins in *N. benthamiana*. Individual values are plotted from a representative experiment (n=12) and are significantly different from each control (Student’s unpaired t-test; *p*<0.0001). These assays were repeated over 4 times each by TM over a 6 month period with similar results. Cloning credits available in **Table S1**.

**Figure S2.**
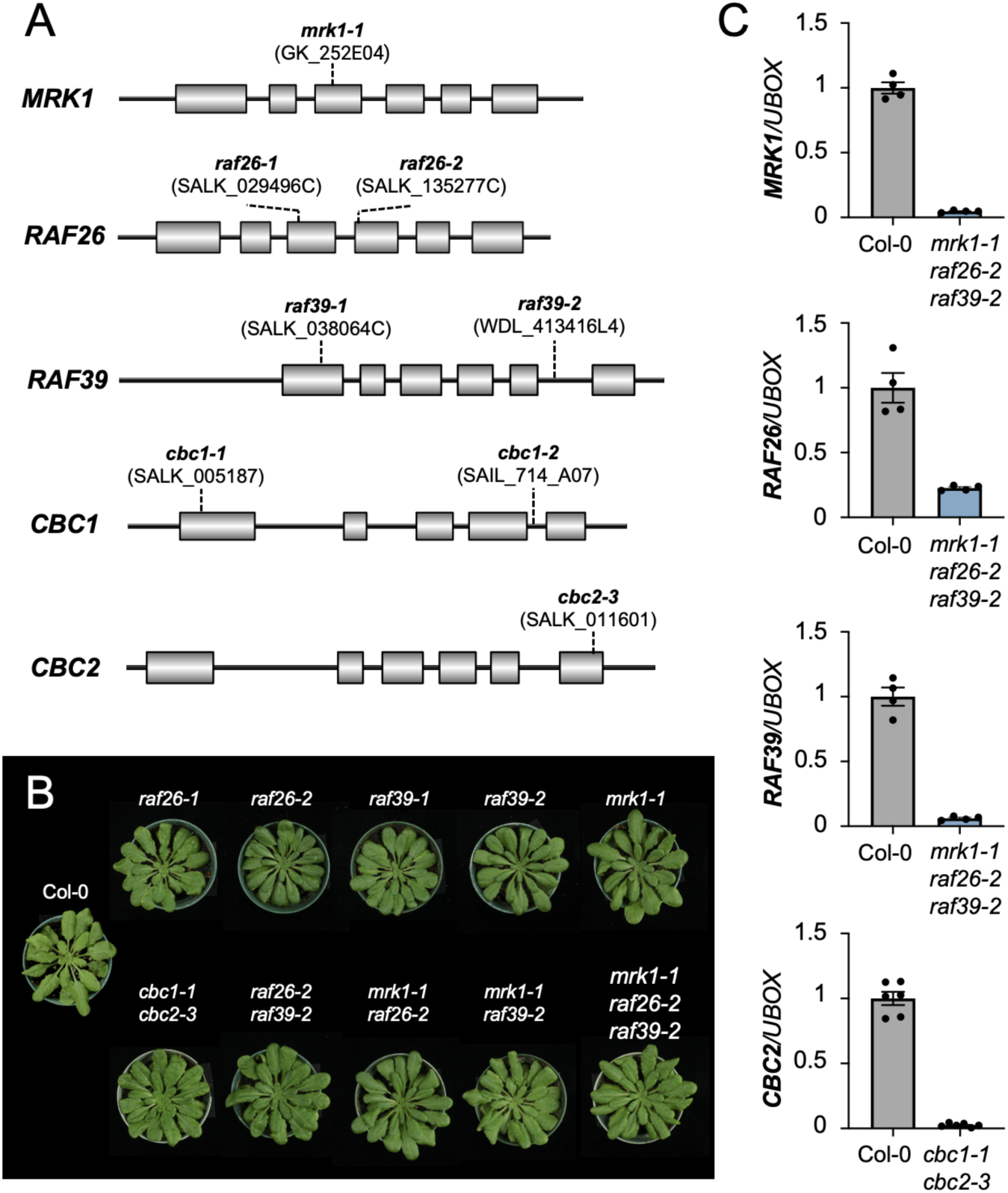
Genetic characterization of C7-Raf loss-of-function mutants. **(A)** Schematic representation, drawn to scale, of subfamily C7 genes, indicating exons (boxes), untranslated regions (lines), and the location of T-DNA insertion alleles. Genomic information was retrieved from The Arabidopsis Information Resource by KRS and JM. Lines were genotyped to homozygosity by KRS, EC, JM, BD, and AR as described in **Table S1**. **(B)** Photographs of representative plants of each genotype after 5 weeks of growth on soil under short-day conditions. Photographs taken by MGD. **(C)** Quantitative real-time qRT-PCR of target genes relative to *UBOX.* Means for 3-4 independent biological replicates are shown +/-standard error of the mean. Data for *MRK1, RAF26,* and *RAF39* expression in *mrk1-1 raf26-2 raf39-2* was collected by AR, while data for *CBC2* expression in *cbc1-1 cbc2-3* was collected by KRS. Lower expression of *CBC1* has already been confirmed for the *cbc1-1* allele (SALK_005187) (Hayashi *et al*., 2020). Primers for genotyping and qRT-PCR are provided in **Table S1**.

**Figure S3.**
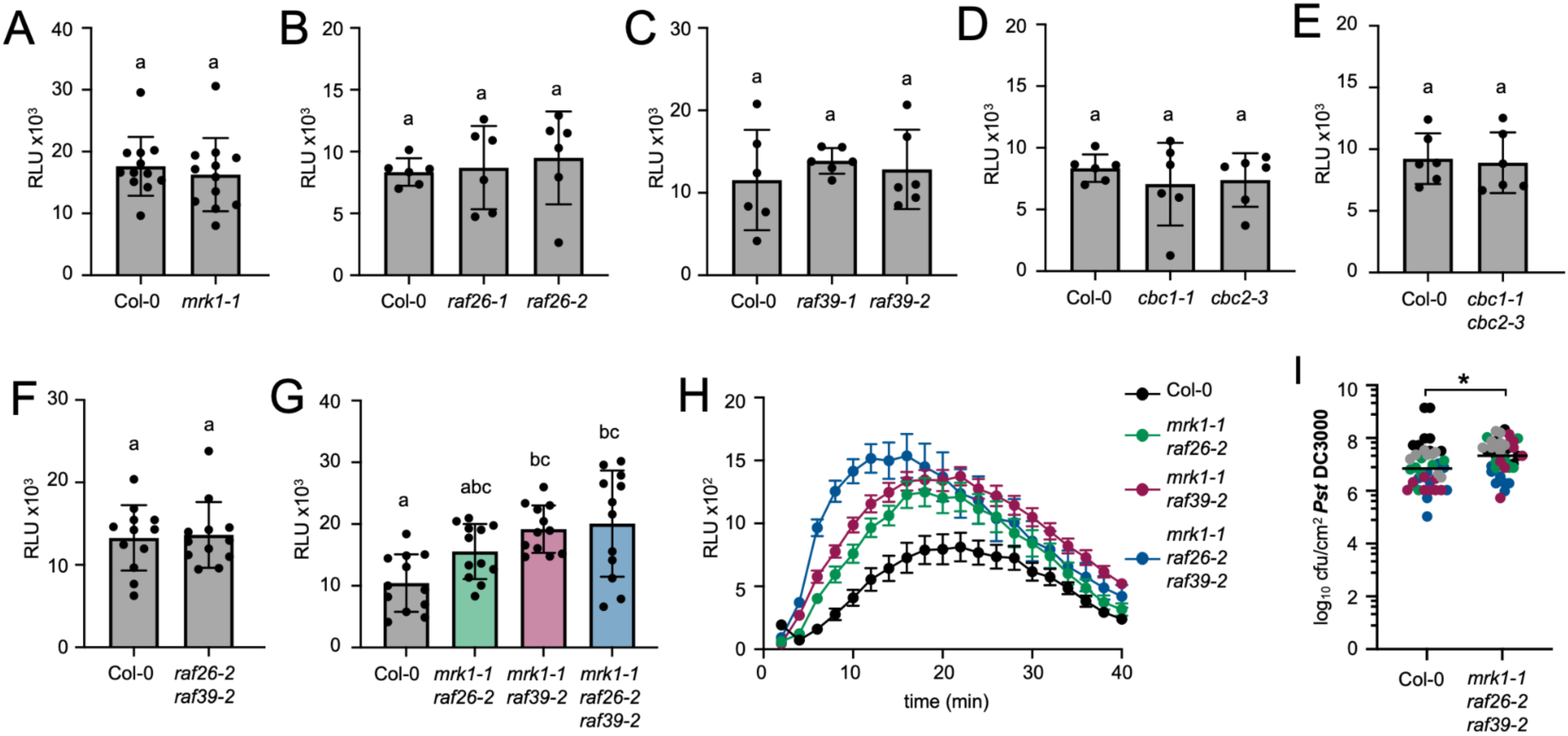
Flg22-triggered ROS in single and double C7-Raf mutants. **(A-H)** ROS production measured in relative light units (RLUs) after treatment with 100 nM flg22. Values represent means +/-standard deviation (n=6-12). Data presented in A, B, and D was collected by KRS; data in C was collected by MGD; data in E was collected by JM; data in F, G, and H was collected by BD. Data in G and H are from the same experiment, presented in G as total RLU and in H as a burst over 40 minutes (values in H are means +/-standard error (n=12). Lower-case letters indicate statistically significant groups determined by a one-way ANOVA followed by Tukey’s post-hoc test (*p*<0.005). These assays were repeated several times over a 5 year period by KRS, MGD, BD and JM; representative experiments are shown. **(I)** Growth of *Pseudomonas syringae* pv. *tomato* (*Pst*) isolate DC3000 3 days after syringe-inoculation. Data from 5 independent biological replicates are plotted together, denoted by black, gray, blue, green, and magenta dots. Values are colony forming units (cfu) per leaf area (cm^2^) from 8 samples per genotype (each sample contains 3 leaf discs from 3 different infected plants). The line represents the mean (n=40). The asterisk indicates slightly significantly different groups, determined by a Student’s unpaired t-test (*p*=0.0281). Data was collected by MGD over a 12 month period. Credits for genetic crosses and genotyping are provided in **Table S1**.

**Figure S4.**
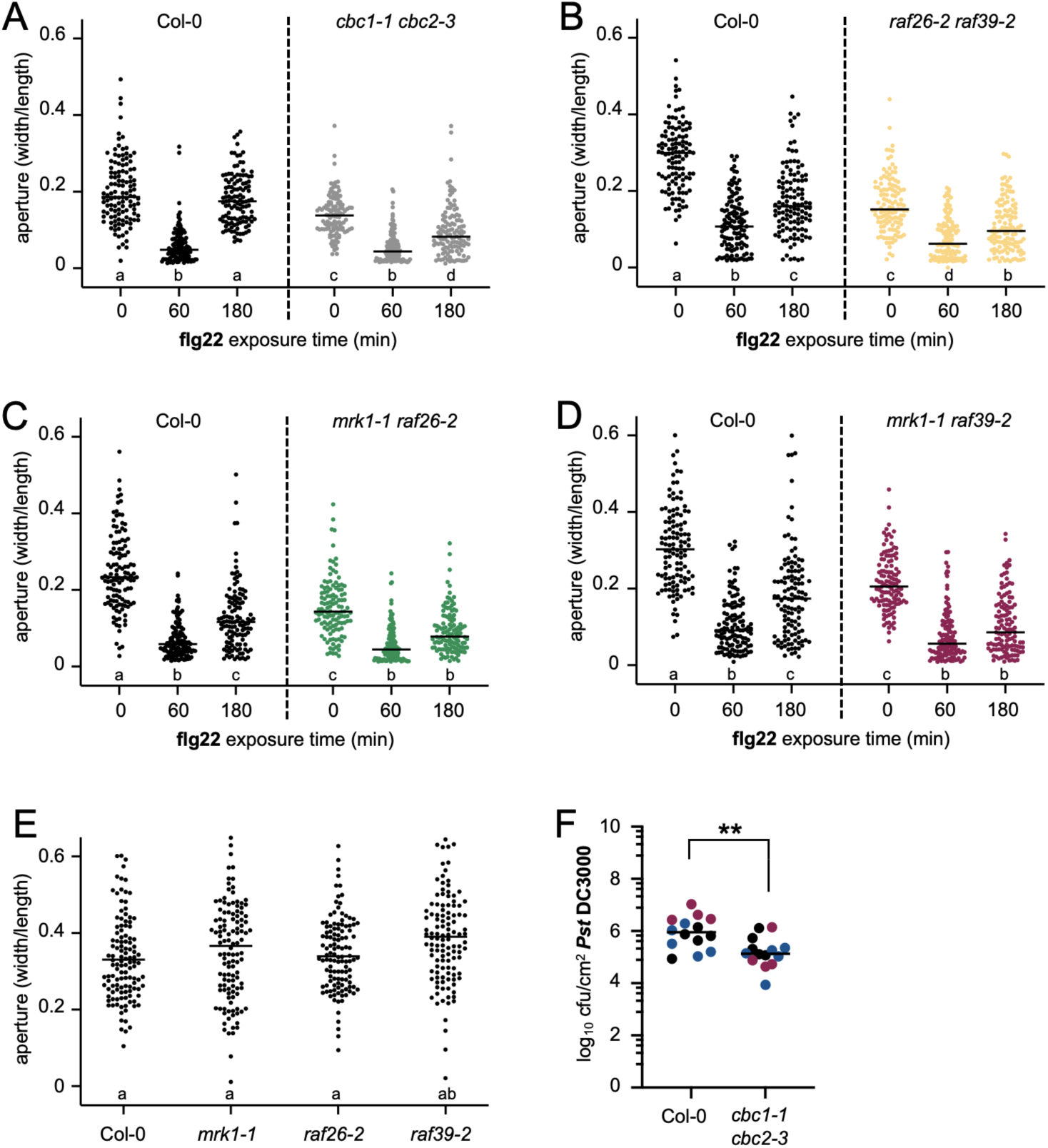
Stomatal aperture in single and double C7-Raf mutants and resistance to *Pst* DC3000 in *cbc1-1 cbc2-3*. **(A-E)** Stomatal apertures prior to (0 min) and following exposure to 1 μM flg22 (60, 180 min). Individual values are plotted and represent ratios of stomatal width:length. The straight line represents the mean (n=120). Lower case letters indicate statistically significant groups, determined by a one-way ANOVA followed by Tukey’s post-hoc test (*p*<0.025). These experiments were completed at least 3 times each over a 12 month period with similar results by BD; representative data is shown. **(F)** Growth of *Pseudomonas syringae* pv. *tomato* (*Pst*) isolate DC3000 3 days after spray-inoculation. Data from 3 independent biological replicates are plotted together, denoted by gray, blue, and magenta dots. Values are colony forming units (cfu) per leaf area (cm^2^) from 4-5 samples per genotype (each sample contains 3 leaf discs from 3 different infected plants). The line represents the mean (n=14). Asterisks indicate significantly different groups, determined by a Student’s unpaired t-test (*p*=0.0026). Data was collected by AR over a 6 month period. Credits for genetic crosses and genotyping are provided in **Table S1**.

**Figure S5.**
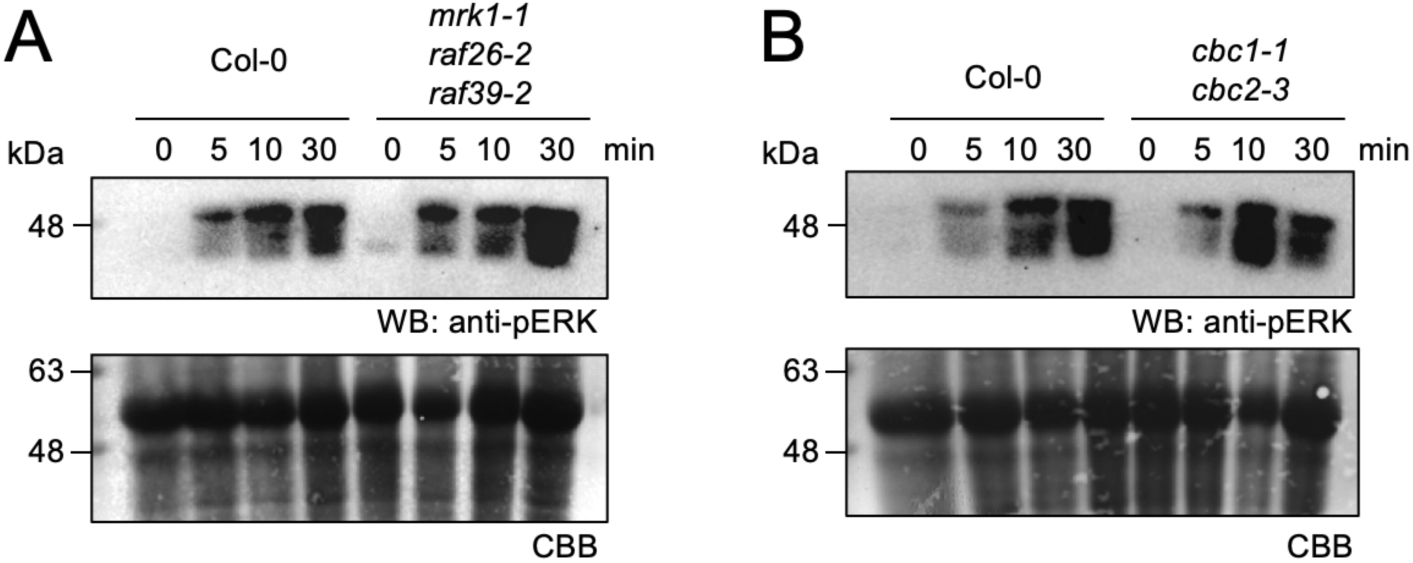
Flg22-triggered activation of MAPK in C7 Raf-like mutants. **(A-B)** Western blots indicating the activation of MAPKs before (0 min) and after exposure to 1 μM flg22 (5, 10, 30 min) in the indicated genotypes. The anti-pERK antibody recognizes the phosphorylated/activated forms of MPK6, MPK3, and MPK4/11. Coomassie Brilliant Blue (CBB) staining of the same membranes indicates loading. Experiments were completed at least 3 times over a 6 month period with similar results by MGD; representative data is shown. Credits for genetic crosses and genotyping are provided in **Table S1**.

**Figure S6.**
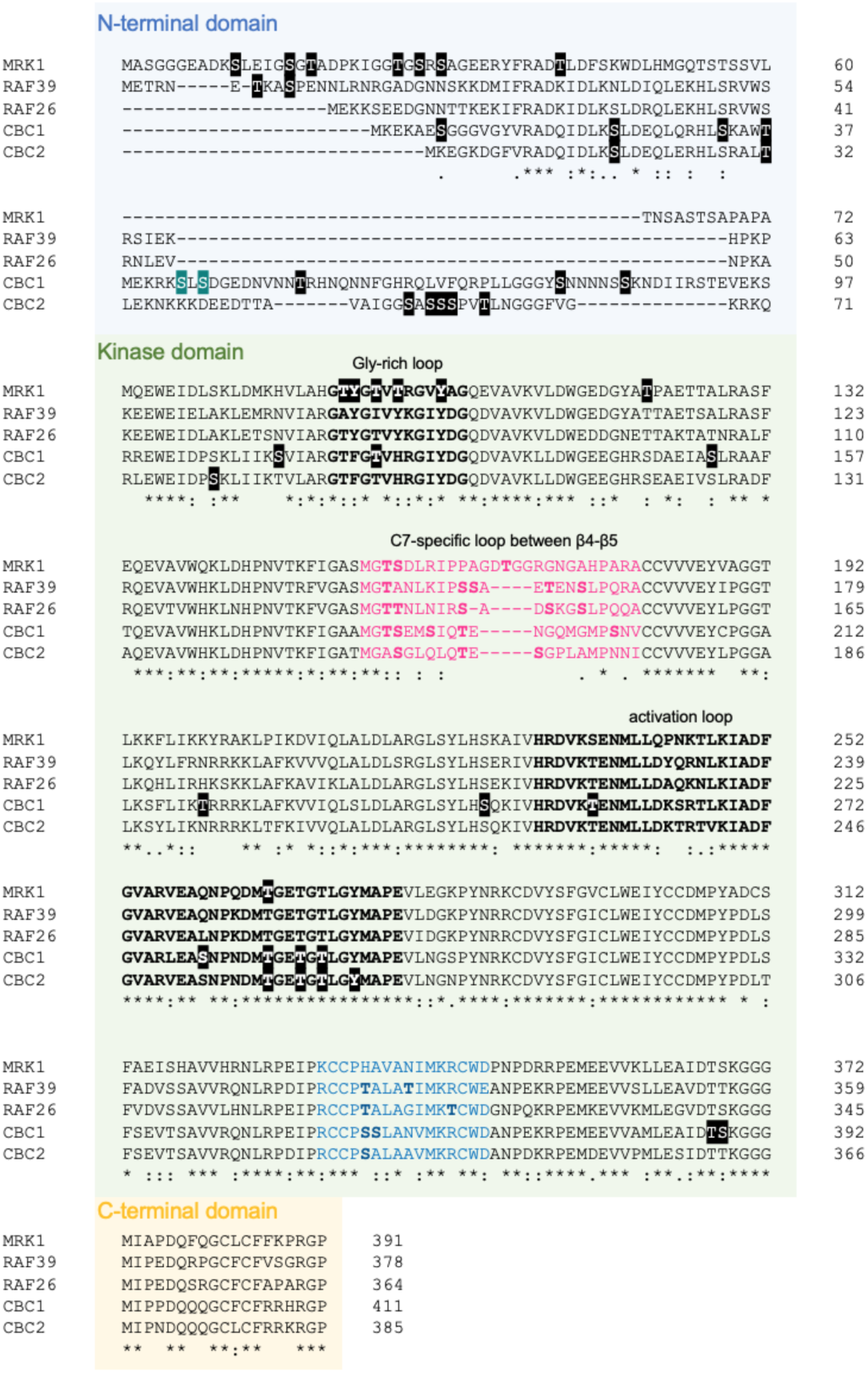
Phosphorylation sites on C7 Raf-like kinases. Multiple sequence alignment of the C7 Raf-like kinases indicating the general locations of the N-terminal (blue), kinase (green), and C-terminal (yellow) domains. Residues outlined in black have been identified as phosphosites curated in online databases based on the following studies (Wang *et al*., 2013a,b; Hoehenwarter *et al*., 2013; Wu *et al*., 2013; Roitinger *et al*., 2015; Marondedze *et al*., 2016; Nukarinen *et al*., 2016; Bhaskara *et al*., 2017; Hiyama *et al*., 2017; Al-Momani *et al*., 2018; Song *et al*., 2018; Takahashi *et al*., 2022). Ser43 and Ser45 of CBC1 are outlined in turquoise as they have been functionally assessed (Hiyama *et al*., 2017). Residues in bold are phosphorylatable residues in regions of sequence divergence compared to MRK1. Alignment was generated using Clustal Omega; asterisks indicate identical amino acids and colons indicate similar amino acids. Analysis by BD and JM.

## Notes

### Competing Interest Statement

The authors have declared no competing interest.

