## Supplementary material for "Subfamily C7 Raf-like kinases MRK1, RAF26, and RAF39 regulate immune homeostasis and stomatal opening in *Arabidopsis thaliana*": File S1

### Analysis of spectral counts of enriched proteins

#### Immunoafinity enrichment (co-IP) assay to study protein-protein interaction

##### For: Jacqueline Monaghan

##### By: Jan Sklenar

##### 24 May, 2023, 18:23

### Protein-protein interaction assay

We study protein-protein interactions using immunoaffinity enrichment experiments, a.k.a. coimmunoprecipitation (coIP). The tagged proteins specifically enrich on an affinity media and the proteins co-purified with them form a list candidates that may form a complex with the tagged protein of interest. To increase specificity of the enrichment we use a negative controls, i.e. wild type, mutant or the tag only that in theory should only capture random or non specificly bound proteins.

### Main script (data analysis)

#### Data description

Proteins identified in immunoafinity enrichments (pull-down) were measured with data dependent method on high resolution LC-MS systems, Orbitrap Fusion. The acquired spectra were peak-picked and searched by Mascot search engine (Matrix Science Ltd.) to identify the peptide sequences from the search space defined by the background proteome. The peptides were combined into proteins based on the principle of parsimony by the search engine. Resulting proteins can be further described by quantitative values based on the number of spectra that identify them. The individual runs were combined in the Scaffold program (Proteome Software Inc.), where the data were evaluated and filtered to contain less than 1% false positives (FDR) and resulted matrix exported as a spreadsheet.

The matrix of proteins detected in different samples serves as the input for an R script for further processing and visualization.

#### Read in the data

Read the ‘csv’ file.

#### Remove contaminants and decoys

Here we remove hits of the typical proteomics artefacts such as keratin, trypsin, and decoy search hits.

#### Define groups of replicates

The raw file names need to be replaced with a comprehensible names, then the samples need to be grouped.

### Calculate total spectral counts in sample groups

Total spectral counts of replicates within the sample groups should not change profoundly and serve as a basic quality control.

### Pairwise comparisons

We reduce the data analysis to a series of binary comparisons by selecting “treatments” and “controls”. This simplifies complex experimental designs and helps to determine how the individual samples compare to controls. A disadvantage is that we may get many plots and tables that has to be compared and evaluated.

```
## The following groups of samples will be compared in
selected_pairs
```

```
## [[1]]
## [[1]]$treat
## [1] "m28_22"
##
## [[1]]$ctrl
## [1] "c_g_22" "c_n_22"
```

```
## Meanningful group names for replicates
names(ename[ename %in% unlist(lapply(selected_pairs, function(x) unique(x)))])
```

```
## [1] "mock_cpk28_LD22" "mock_cpk28_LD22" "mock_cpk28_LD22" "ctrl_nls_L22"
## [5] "ctrl_nls_L22"    "ctrl_nls_L22"    "ctrl_gfp_L22"    "ctrl_gfp_L22"
```

```
## Numbers of replicates
lengths <- lapply(selected_pairs_cols, function(x) sapply(x, function(y) length(y)))
lengths
```

```
## [[1]]
## treat  ctrl
##     3     5
```

### Main calculations

1. Removal of typical contaminants, such as keratin, trypsin, etc.
2. Imputation of missing values to allow the fold change calculation
3. Calculate mean spectral count for the treatment and the control
4. Calculate ratio log2(treatment/control)
5. Set threshold for the ratios that signify the specific protein-protein interaction
6. Set reproducibility Filter - a percentage of replicates the proteins must be found therein
7. Visualize the results in a bar-plot and a table

### Calculations in details

##### Imputation

Data imputation in proteomics is an active research area. While several aproaches were suggested, there has not been a clear consensus in the community of how to deal with the missing data in various experiments and data formats. Here our strategy attempts for the imputation to be as optimistic as possible to provide a list of possible protein-protein interaction candidates. We cannot make any assumptions on the number of proteins that will be enriched and the abundance of the specific proteins. For this reason normalization is not used, and we simply assume the same amount of input tissue was used in every experiment. We can expect a substantial number of proteins will bind non specifically to the immunoafinity beads. After the enrichment, we are dealing with a specific sub-proteome. These facts are contributing factors to many missing values are often observed. We impute them in the following stages.

- Zeros in Scaffold means a lower probability hit, we convert is to 1.
- Missing values are then imputed within sample groups in pairs being compared.
- A group unique hits (with the missing values in the other group) are converted to 1.
- All the other missing values within sample groups are imputed as a mean. This has no effect on the mean magnitude.

##### Fold changes

After the missing value imputation we can calculates mean log2 fold changes in the selected sample groups, treatment and control. Then we take ratio treatment / control. We mark proteins that were identified uniquely in the groups.

### Filters used

Filters helps us to bring forward the specific and reproducible results.

```
## Fold change filter (log2) - we keep 'larger than' AND 'smaller than'
## a specified value, with exception of the hits
## identified uniquely (with a missing value in the other sample type)
thres <- 1

## Reproducibility filter (%) - we keep only hits found in at least
## certain percentage of replicates (of the same sample type)
perc  <- 50
```

### Plots

### Result evaluation

The plots below show the calculated log2 fold changes between the treatment and the control.

Three different situations are color coded:

- Red: Unique in Treatment [t]
- Blue: Unique in Control [c]
- Green: Ratio [t/c]

In the immunoafinity enrichment experiments, the proteins of interest can be identified exclusively in the samples with the tagged protein [t] (red) or in controls [c] (blue). While the former is the sought result, the latter proteins are irrelevant to studied tagged protein, identified most likely due to their non specific binding to the affinity media. The proteins that occur both in the control and in the treatment [t/c] (green) could be binding the tagged protein specifically if the ratio shows an extreme (positive) value.

***It is our evaluation that makes identified proteins a possible protein-protein interaction candidate with given tagged protein.***

The fold change ratio could only be calculated by imputing 1 in the control. That often makes proteins appearing among the small ratios. In this situation there was nothing found in the control and we have to ask: Is the fact we do not see anything in the control due to the complex of tagged protein that interacts with the co-purified protein or was the protein found by a chance? Answer is not simple, however more spectra we found increases the that a protein is part of the complex with our tagged protein. Moreover, we require reproducibility of protein detection in biological sample replicates as another criterion for labeling protein as an interaction candidate.

```
## [1] "mock_cpk28_LD22 [t] / ctrl_nls_L22 [c]; ctrl_gfp_L22 [c]"
```

```
##
##
## =====================
## [1] "Pair No.:  1"
## =====================
```

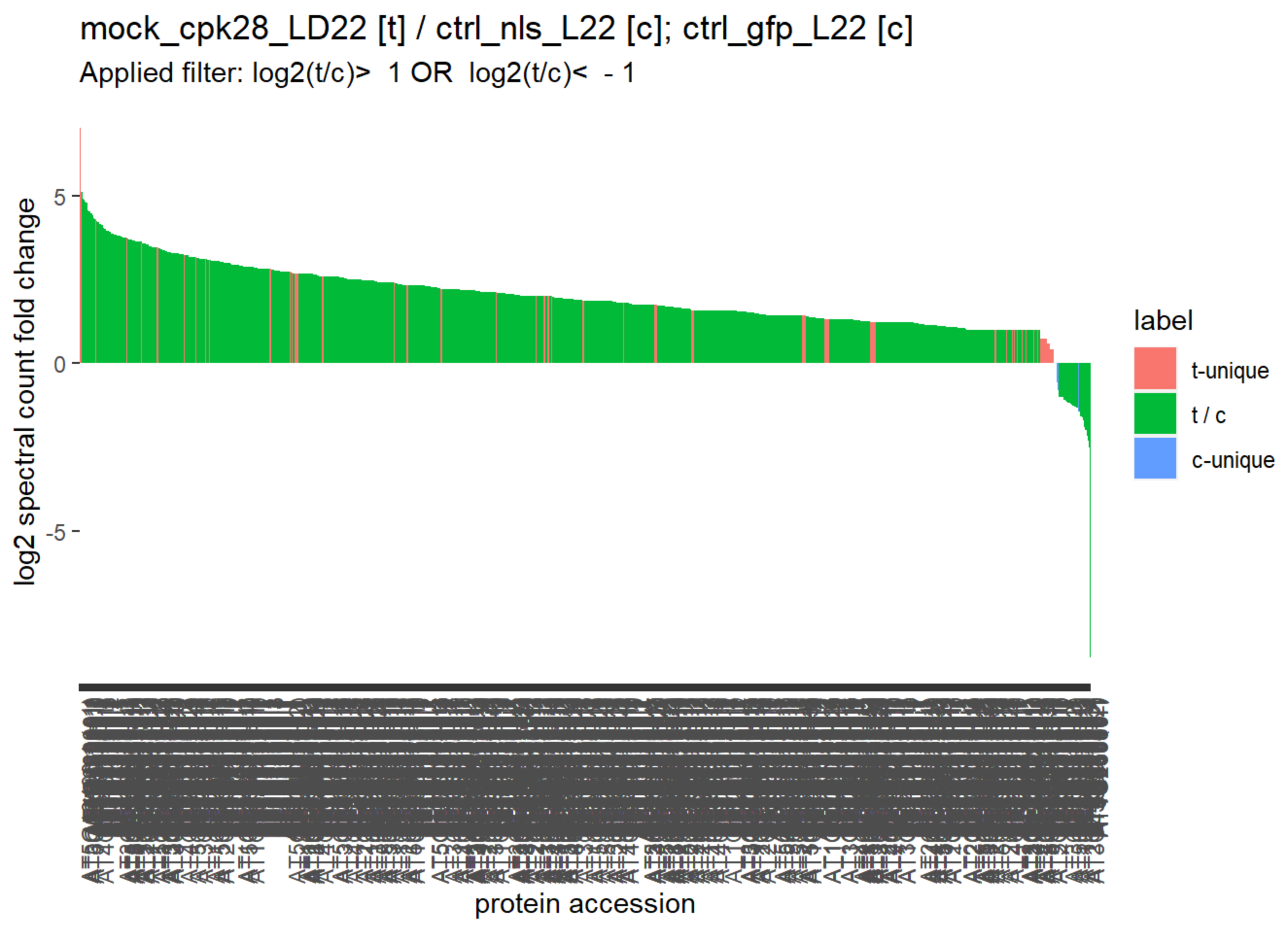

```
##
##
## |     |Accession.Number | tc_lg|label    |description
|
## |:----|:----------------|-----:|:--------|:---------------------------------------
--|
## |1220 |AT5G13490.1 (+1) |  7.04|t-unique |AAC2 &#124; ADP/ATP carrier 2 &#124; chr5:4
3 |
## |125  |AT4G33090.1      |  5.44|t / c    |APM1, ATAPM1 &#124; aminopeptidase M1 &#12
4; |
## |1811 |AT5G43010.1      |  5.10|t-unique |RPT4A &#124; regulatory particle triple
|
## |227  |AT3G08510.1 (+1) |  5.10|t / c    |ATPLC2, PLC2 &#124; phospholipase C 2 &#12
4; |
## |763  |AT4G28520.1      |  4.92|t / c    |CRU3, CRC &#124; cruciferin 3 &#124; chr4:1
4 |
## |626  |AT2G17890.1      |  4.88|t / c    |CPK16 &#124; calcium-dependent protein
|
## |1    |AT5G66210.1 (+1) |  4.82|t / c    |CPK28 &#124; calcium-dependent protein
|
## |1184 |AT1G75780.1      |  4.79|t / c    |TUB1 &#124; tubulin beta-1 chain &#124; chr
1 |
## |236  |AT4G22690.1      |  4.78|t / c    |CYP706A1 &#124; cytochrome P450, family
|
## |147  |AT3G46970.1      |  4.56|t / c    |ATPHS2, PHS2 &#124; alpha-glucan phosph
|
## |43   |AT2G18960.1      |  4.53|t / c    |AHA1, PMA, OST2, HA1 &#124; H(+)-ATPase
|
## |390  |AT4G35470.1      |  4.52|t / c    |PIRL4 &#124; plant intracellular ras gr
|
## |490  |AT1G17840.1      |  4.48|t / c    |WBC11, ABCG11, DSO, COF1, ATWBC11
|
## |488  |AT1G54520.1      |  4.46|t / c    |&#124; unknown protein; FUNCTIONS IN:
|
## |162  |AT3G09840.1      |  4.42|t / c    |CDC48, ATCDC48, CDC48A &#124; cell divi
|
## |212  |AT1G02150.1      |  4.36|t / c    |&#124; Tetratricopeptide repeat (TPR)-
|
## |377  |AT1G06410.1      |  4.32|t / c    |ATTPS7, TPS7, ATTPSA &#124; trehalose-p
|
## |169  |AT4G31480.1 (+1) |  4.30|t / c    |&#124; Coatomer, beta subunit &#124; chr4:1
|
## |499  |AT5G46580.1      |  4.30|t-unique |&#124; pentatricopeptide (PPR) repeat-
|
## |109  |AT1G59870.1      |  4.23|t / c    |PEN3, PDR8, ATPDR8, ABCG36, ATABCG
|
## |450  |AT2G45960.3      |  4.22|t / c    |PIP1B, TMP-A, ATHH2, PIP1;2 &#124; plas
|
## |460  |AT5G16715.1      |  4.22|t / c    |EMB2247 &#124; ATP binding;valine-tRNA
|
## |187  |AT5G65720.1      |  4.16|t / c    |ATNIFS1, NIFS1, NFS1, ATNFS1 &#124; nit
|
```

```
## |459  |AT3G14840.2      |  4.14|t / c    |&#124; Leucine-rich repeat transmembra
|
## |385  |AT4G30190.1      |  4.12|t / c    |AHA2, PMA2, HA2 &#124; H(+)-ATPase 2 &#124;
|
## |588  |AT4G10790.1      |  4.12|t / c    |&#124; UBX domain-containing protein &#124;
|
## |1210 |AT3G21370.1      |  4.04|t / c    |BGLU19 &#124; beta glucosidase 19 &#124; ch
r |
## |481  |AT2G44490.1      |  4.00|t / c    |PEN2, BGLU26 &#124; Glycosyl hydrolase
|
## |500  |AT1G79050.1      |  4.00|t / c    |&#124; recA DNA recombination family p
|
## |1733 |AT2G47650.1      |  4.00|t / c    |UXS4 &#124; UDP-xylose synthase 4 &#124; ch
r |
## |76   |AT5G46800.1      |  3.94|t / c    |BOU &#124; Mitochondrial substrate carr
|
## |416  |AT2G41560.1      |  3.94|t / c    |ACA4 &#124; autoinhibited Ca(2+)-ATPase
|
## |526  |AT2G22500.1      |  3.94|t / c    |UCP5, ATPUMP5, DIC1 &#124; uncoupling p
|
## |472  |AT2G41790.1      |  3.93|t / c    |&#124; Insulinase (Peptidase family M1
|
## |307  |AT1G01790.1      |  3.91|t / c    |KEA1, ATKEA1 &#124; K+ efflux antiporte
|
## |405  |AT5G20490.1      |  3.87|t / c    |XIK, ATXIK, XI-17 &#124; Myosin family
|
## |554  |AT5G01920.1      |  3.87|t / c    |STN8 &#124; Protein kinase superfamily
|
## |229  |AT1G68830.1      |  3.86|t / c    |STN7 &#124; STT7 homolog STN7 &#124; chr1:2
5 |
## |509  |AT1G56500.1      |  3.84|t / c    |&#124; haloacid dehalogenase-like hydr
|
## |1163 |AT1G05150.1      |  3.84|t / c    |&#124; Calcium-binding tetratricopepti
|
## |419  |AT5G38660.2      |  3.83|t / c    |APE1 &#124; acclimation of photosynthes
|
## |258  |AT5G62670.1      |  3.82|t / c    |AHA11, HA11 &#124; H(+)-ATPase 11 &#124; ch
r |
## |516  |AT3G59350.1 (+1) |  3.81|t / c    |&#124; Protein kinase superfamily prot
|
## |562  |AT5G05170.1      |  3.81|t / c    |CESA3, IXR1, ATCESA3, ATH-B, CEV1
|
## |610  |AT1G30360.1      |  3.81|t / c    |ERD4 &#124; Early-responsive to dehydra
|
## |1076 |AT3G25800.1      |  3.81|t / c    |PDF1, PR 65, PP2AA2 &#124; protein phos
|
## |143  |AT1G80480.1      |  3.77|t / c    |PTAC17 &#124; plastid transcriptionally
|
## |226  |AT5G05010.1 (+1) |  3.77|t / c    |&#124; clathrin adaptor complexes medi
|
## |411  |AT2G46520.1      |  3.74|t / c    |&#124; cellular apoptosis susceptibili
|
```

```
## |531  |AT3G50950.1 (+1) |  3.74|t / c    |ZAR1 &#124; HOPZ-ACTIVATED RESISTANCE 1
|
## |575  |AT5G57110.1 (+1) |  3.74|t / c    |ACA8, AT-ACA8 &#124; autoinhibited Ca2+
|
## |713  |AT2G39010.1      |  3.74|t / c    |PIP2E, PIP2;6 &#124; plasma membrane in
|
## |1028 |AT5G43470.1 (+1) |  3.74|t-unique |RPP8, HRT, RCY1 &#124; Disease resistan
|
## |281  |AT1G69830.1      |  3.72|t / c    |ATAMY3, AMY3 &#124; alpha-amylase-like
|
## |83   |AT5G14040.1      |  3.70|t / c    |PHT3;1 &#124; phosphate transporter 3;1
|
## |603  |AT4G17090.1      |  3.70|t / c    |CT-BMY, BAM3, BMY8 &#124; chloroplast b
|
## |1245 |AT2G47450.1      |  3.70|t / c    |CAO, CPSRP43 &#124; chloroplast signal
|
## |128  |AT1G78570.1      |  3.69|t / c    |RHM1, ROL1, ATRHM1 &#124; rhamnose bios
|
## |403  |AT1G10510.1      |  3.68|t / c    |emb2004 &#124; RNI-like superfamily pro
|
## |517  |AT1G13320.1 (+1) |  3.66|t / c    |PP2AA3 &#124; protein phosphatase 2A  s
|
## |656  |AT5G48880.2 (+1) |  3.66|t / c    |PKT2, KAT5 &#124; peroxisomal 3-keto-ac
|
## |408  |AT1G80410.2      |  3.65|t / c    |EMB2753 &#124; tetratricopeptide repeat
|
## |96   |AT2G36250.1 (+1) |  3.64|t / c    |FTSZ2-1, ATFTSZ2-1 &#124; Tubulin/FtsZ
|
## |256  |AT2G15620.1      |  3.64|t / c    |NIR1, NIR, ATHNIR &#124; nitrite reduct
|
## |150  |AT2G20580.1      |  3.63|t / c    |RPN1A, ATRPN1A &#124; 26S proteasome re
|
## |371  |AT4G23650.1      |  3.62|t / c    |CDPK6, CPK3 &#124; calcium-dependent pr
|
## |618  |AT5G22800.1      |  3.62|t / c    |EMB86, EMB1030, EMB263 &#124; Alanyl-tR
|
## |629  |AT3G63260.1      |  3.62|t / c    |ATMRK1 &#124; Protein kinase superfamil
|
## |733  |AT1G06700.1 (+1) |  3.62|t / c    |&#124; Protein kinase superfamily prot
|
## |986  |AT1G52290.1      |  3.58|t-unique |&#124; Protein kinase superfamily prot
|
## |527  |AT3G27240.1      |  3.58|t / c    |&#124; Cytochrome C1 family &#124; chr3:100
|
## |776  |AT5G55610.1      |  3.58|t / c    |&#124; unknown protein; LOCATED IN: mi
|
## |273  |AT1G73110.1      |  3.57|t / c    |&#124; P-loop containing nucleoside tr
|
## |367  |AT1G22410.1      |  3.57|t / c    |&#124; Class-II DAHP synthetase family
|
## |183  |AT5G60790.1      |  3.55|t / c    |ATGCN1, GCN1 &#124; ABC transporter fam
|
```

```
## |572  |AT1G03160.1      |  3.54|t / c    |FZL &#124; FZO-like &#124; chr1:761321-7660
5 |
## |463  |AT5G66680.1      |  3.53|t / c    |DGL1 &#124; dolichyl-diphosphooligosacc
|
## |686  |AT4G19710.2      |  3.50|t / c    |AK-HSDH II, AK-HSDH &#124; aspartate ki
|
## |852  |AT2G26250.1      |  3.50|t / c    |FDH, KCS10 &#124; 3-ketoacyl-CoA syntha
|
## |897  |AT5G57350.1 (+1) |  3.48|t / c    |AHA3, ATAHA3, HA3 &#124; H(+)-ATPase 3
|
## |515  |AT1G74960.1 (+2) |  3.47|t / c    |FAB1, KAS2, ATKAS2 &#124; fatty acid bi
|
## |196  |AT4G29130.1      |  3.46|t / c    |ATHXK1, GIN2, HXK1 &#124; hexokinase 1
|
## |537  |AT1G01960.1      |  3.46|t / c    |EDA10 &#124; SEC7-like guanine nucleoti
|
## |545  |AT4G16390.1      |  3.46|t / c    |SVR7 &#124; pentatricopeptide (PPR) rep
|
## |595  |AT2G32450.1      |  3.46|t / c    |&#124; Calcium-binding tetratricopepti
|
## |708  |AT3G09740.1      |  3.46|t / c    |SYP71, ATSYP71 &#124; syntaxin of plant
|
## |712  |AT1G30470.1      |  3.46|t-unique |&#124; SIT4 phosphatase-associated fam
|
## |926  |AT2G31880.1      |  3.46|t-unique |SOBIR1, EVR &#124; Leucine-rich repeat
|
## |504  |AT3G62830.1 (+1) |  3.42|t / c    |UXS2, ATUXS2, AUD1 &#124; NAD(P)-bindin
|
## |1231 |AT2G22360.1      |  3.42|t / c    |&#124; DNAJ heat shock family protein
|
## |140  |AT3G19820.1 (+2) |  3.41|t / c    |DWF1, DIM, EVE1, DIM1, CBB1 &#124; cell
|
## |559  |AT5G55280.1      |  3.39|t / c    |FTSZ1-1, ATFTSZ1-1, CPFTSZ &#124; homol
|
## |721  |AT2G32080.1 (+1) |  3.37|t / c    |PUR ALPHA-1 &#124; purin-rich alpha 1 &#12
4; |
## |725  |AT3G01060.1      |  3.37|t / c    |&#124; unknown protein; Has 640 Blast
|
## |1698 |AT2G19860.1      |  3.37|t / c    |ATHXK2, HXK2 &#124; hexokinase 2 &#124; chr
2 |
## |20   |AT3G08580.1 (+1) |  3.35|t / c    |AAC1 &#124; ADP/ATP carrier 1 &#124; chr3:2
6 |
## |164  |AT3G29320.1      |  3.35|t / c    |&#124; Glycosyl transferase, family 35
|
## |370  |AT5G27380.1      |  3.32|t / c    |GSH2, GSHB &#124; glutathione synthetas
|
## |748  |AT2G27600.1      |  3.32|t / c    |SKD1, VPS4, ATSKD1 &#124; AAA-type ATPa
|
## |751  |AT1G60780.1      |  3.32|t / c    |HAP13 &#124; Clathrin adaptor complexes
|
## |274  |AT3G59780.1      |  3.31|t / c    |&#124; Rhodanese/Cell cycle control ph
|
```

```
## |1356 |AT5G38660.1      |  3.30|t / c    |APE1 &#124; acclimation of photosynthes
|
## |551  |AT5G17380.1      |  3.29|t / c    |&#124; Thiamine pyrophosphate dependen
|
## |573  |AT5G58260.1      |  3.29|t / c    |&#124; oxidoreductases, acting on NADH
|
## |464  |AT5G23540.1      |  3.28|t / c    |&#124; Mov34/MPN/PAD-1 family protein
|
## |292  |AT1G53750.1      |  3.27|t / c    |RPT1A &#124; regulatory particle triple
|
## |532  |AT3G55360.1      |  3.27|t / c    |CER10, ECR, ATTSC13, TSC13 &#124; 3-oxo
|
## |769  |AT1G04810.1      |  3.27|t / c    |&#124; 26S proteasome regulatory compl
|
## |797  |AT3G44340.1      |  3.27|t / c    |CEF &#124; clone eighty-four &#124; chr3:16
0 |
## |901  |AT3G53520.4      |  3.27|t / c    |UXS1 &#124; UDP-glucuronic acid decarbo
|
## |280  |AT4G22890.1 (+2) |  3.26|t / c    |PGR5-LIKE A &#124; PGR5-LIKE A &#124; chr4:
1 |
## |295  |AT4G39980.1      |  3.26|t / c    |DHS1 &#124; 3-deoxy-D-arabino-heptuloso
|
## |773  |AT4G00630.1      |  3.26|t / c    |KEA2, ATKEA2 &#124; K+ efflux antiporte
|
## |507  |AT1G18270.3      |  3.25|t / c    |&#124; ketose-bisphosphate aldolase cl
|
## |541  |AT5G08540.1      |  3.25|t / c    |&#124; unknown protein; FUNCTIONS IN:
|
## |612  |AT5G17990.1      |  3.25|t / c    |TRP1, pat1 &#124; tryptophan biosynthes
|
## |949  |AT4G37200.1      |  3.22|t-unique |HCF164 &#124; Thioredoxin superfamily p
|
## |266  |AT4G34450.1      |  3.22|t / c    |&#124; coatomer gamma-2 subunit, putat
|
## |323  |AT1G50480.1      |  3.22|t / c    |THFS &#124; 10-formyltetrahydrofolate s
|
## |363  |AT3G54110.1      |  3.22|t / c    |ATPUMP1, UCP, PUMP1, ATUCP1, UCP1
|
## |677  |AT5G06530.1 (+1) |  3.22|t / c    |&#124; ABC-2 type transporter family p
|
## |624  |AT1G64740.1      |  3.18|t / c    |TUA1 &#124; alpha-1 tubulin &#124; chr1:240
5 |
## |485  |AT3G24430.1      |  3.17|t / c    |HCF101 &#124; ATP binding &#124; chr3:88687
3 |
## |694  |AT5G54160.1      |  3.17|t / c    |ATOMT1, OMT1 &#124; O-methyltransferase
|
## |756  |AT4G25960.1      |  3.17|t / c    |PGP2 &#124; P-glycoprotein 2 &#124; chr4:13
1 |
## |920  |AT1G70940.1      |  3.17|t / c    |PIN3, ATPIN3 &#124; Auxin efflux carrie
|
## |930  |AT4G30340.1      |  3.17|t / c    |ATDGK7, DGK7 &#124; diacylglycerol kina
|
```

```
## |983  |AT3G61430.1 (+1) |  3.17|t / c    |PIP1A, ATPIP1, PIP1, PIP1;1 &#124; plas
|
## |1009 |AT4G36070.2      |  3.17|t / c    |CPK18 &#124; calcium-dependent protein
|
## |913  |AT5G66200.1      |  3.17|t-unique |ARO2 &#124; armadillo repeat only 2 &#124;
c |
## |246  |AT1G70410.2      |  3.14|t / c    |ATBCA4, BCA4 &#124; beta carbonic anhyd
|
## |648  |AT3G26520.1      |  3.14|t / c    |TIP2, SITIP, GAMMA-TIP2, TIP1;2 &#124;
|
## |120  |AT5G19760.1      |  3.13|t / c    |&#124; Mitochondrial substrate carrier
|
## |560  |AT5G51820.1      |  3.12|t / c    |PGM, ATPGMP, PGM1, STF1 &#124; phosphog
|
## |568  |AT2G04842.1      |  3.12|t / c    |EMB2761 &#124; threonyl-tRNA synthetase
|
## |577  |AT5G61020.1      |  3.12|t / c    |ECT3 &#124; evolutionarily conserved C-
|
## |742  |AT4G29900.1      |  3.12|t / c    |ACA10, CIF1, ATACA10 &#124; autoinhibit
|
## |749  |AT5G58410.1      |  3.12|t / c    |&#124; HEAT/U-box domain-containing pr
|
## |831  |AT5G52320.1      |  3.12|t / c    |CYP96A4 &#124; cytochrome P450, family
|
## |846  |AT3G10380.1      |  3.12|t / c    |SEC8, ATSEC8 &#124; subunit of exocyst
|
## |701  |AT1G45474.1 (+1) |  3.12|t-unique |LHCA5 &#124; photosystem I light harves
|
## |423  |AT3G52750.1      |  3.11|t / c    |FTSZ2-2 &#124; Tubulin/FtsZ family prot
|
## |875  |AT3G07770.1      |  3.09|t / c    |Hsp89.1, AtHsp90.6, AtHsp90-6 &#124; HE
|
## |1200 |AT4G31390.1      |  3.09|t / c    |&#124; Protein kinase superfamily prot
|
## |1479 |AT5G11380.1      |  3.09|t-unique |DXPS3 &#124; 1-deoxy-D-xylulose 5-phosp
|
## |542  |AT4G30010.1      |  3.08|t / c    |&#124; unknown protein; FUNCTIONS IN:
|
## |579  |AT3G07160.1      |  3.06|t / c    |ATGSL10, gsl10, CALS9 &#124; glucan syn
|
## |674  |AT5G64940.1 (+1) |  3.06|t / c    |ATATH13, ATH13, ATOSA1, OSA1 &#124; ABC
|
## |802  |AT5G19690.1      |  3.06|t / c    |STT3A &#124; staurosporin and temperatu
|
## |841  |AT3G45190.1      |  3.06|t / c    |&#124; SIT4 phosphatase-associated fam
|
## |1029 |AT1G31230.1      |  3.06|t / c    |AK-HSDH I, AK-HSDH &#124; aspartate kin
|
## |85   |AT2G38040.1 (+1) |  3.05|t / c    |CAC3 &#124; acetyl Co-enzyme a carboxyl
|
## |38   |AT1G06950.1      |  3.04|t / c    |ATTIC110, TIC110 &#124; translocon at t
|
```

```
## |134  |AT5G54770.1      |  3.04|t / c    |THI1, TZ, THI4 &#124; thiazole biosynth
|
## |530  |AT1G79560.1      |  3.04|t / c    |EMB156, EMB36, EMB1047, FTSH12 &#124; F
|
## |602  |AT3G26710.1      |  3.04|t / c    |CCB1 &#124; cofactor assembly of comple
|
## |248  |AT2G32730.1      |  3.03|t / c    |&#124; 26S proteasome regulatory compl
|
## |253  |AT5G12470.1      |  3.03|t / c    |&#124; Protein of unknown function (DU
|
## |277  |AT5G19990.1      |  3.02|t / c    |RPT6A, ATSUG1 &#124; regulatory particl
|
## |194  |AT4G29040.1      |  3.01|t / c    |RPT2a &#124; regulatory particle AAA-AT
|
## |98   |AT5G44340.1      |  3.00|t / c    |TUB4 &#124; tubulin beta chain 4 &#124; chr
5 |
## |609  |AT2G45710.1      |  3.00|t / c    |&#124; Zinc-binding ribosomal protein
|
## |642  |AT2G37710.1      |  3.00|t / c    |RLK &#124; receptor lectin kinase &#124; ch
r |
## |784  |AT2G18730.1      |  3.00|t / c    |ATDGK3, DGK3 &#124; diacylglycerol kina
|
## |832  |AT1G51500.1      |  3.00|t / c    |CER5, D3, ABCG12, WBC12, ATWBC12 &#124;
|
## |887  |AT3G51140.1      |  3.00|t / c    |&#124; Protein of unknown function (DU
|
## |290  |AT5G22060.1      |  2.98|t / c    |ATJ2, J2 &#124; DNAJ homologue 2 &#124; chr
5 |
## |1197 |AT4G28470.1      |  2.98|t / c    |RPN1B, ATRPN1B &#124; 26S proteasome re
|
## |60   |AT1G09340.1      |  2.97|t / c    |CRB, CSP41B, HIP1.3 &#124; chloroplast
|
## |617  |AT5G13630.1      |  2.94|t / c    |GUN5, CCH, CHLH, CCH1, ABAR &#124; magn
|
## |811  |AT5G05780.1      |  2.94|t / c    |RPN8A, AE3, ATHMOV34 &#124; RP non-ATPa
|
## |820  |AT3G20000.1      |  2.94|t / c    |TOM40 &#124; translocase of the outer m
|
## |894  |AT3G10920.2      |  2.94|t / c    |MSD1 &#124; manganese superoxide dismut
|
## |1027 |AT3G23750.1      |  2.94|t / c    |&#124; Leucine-rich repeat protein kin
|
## |1113 |AT1G64430.1 (+1) |  2.94|t / c    |&#124; Pentatricopeptide repeat (PPR)
|
## |289  |AT5G14780.1      |  2.93|t / c    |FDH &#124; formate dehydrogenase &#124; chr
5 |
## |510  |AT4G39960.1      |  2.93|t / c    |&#124; Molecular chaperone Hsp40/DnaJ
|
## |13   |AT5G62690.1 (+1) |  2.92|t / c    |TUB2 &#124; tubulin beta chain 2 &#124; chr
5 |
## |278  |AT5G17170.1      |  2.92|t / c    |ENH1 &#124; rubredoxin family protein &#12
4; |
```

```
## |184  |AT4G24190.1      |  2.91|t / c    |SHD, HSP90.7, AtHsp90.7, AtHsp90-7
|
## |358  |AT1G03475.1      |  2.91|t / c    |LIN2, HEMF1, ATCPO-I &#124; Coproporphy
|
## |581  |AT3G43300.1      |  2.90|t / c    |ATMIN7, BEN1 &#124; HOPM interactor 7 &#12
4; |
## |497  |AT4G35830.1      |  2.88|t / c    |ACO1 &#124; aconitase 1 &#124; chr4:1697300
7 |
## |866  |AT3G02450.1      |  2.88|t / c    |&#124; cell division protein ftsH, put
|
## |589  |AT1G70770.1 (+1) |  2.87|t / c    |&#124; Protein of unknown function DUF
|
## |659  |AT4G21710.1      |  2.87|t / c    |NRPB2, EMB1989, RPB2 &#124; DNA-directe
|
## |778  |AT1G11330.1      |  2.87|t / c    |&#124; S-locus lectin protein kinase f
|
## |809  |AT5G04130.1      |  2.87|t / c    |GYRB2 &#124; DNA GYRASE B2 &#124; chr5:1122
0 |
## |960  |AT1G25490.1      |  2.87|t / c    |RCN1, REGA, ATB BETA BETA, EER1 &#124;
|
## |997  |AT4G04770.1      |  2.87|t / c    |ATABC1, LAF6, ATNAP1, ABC1 &#124; ATP b
|
## |1039 |AT1G77590.1      |  2.87|t / c    |LACS9 &#124; long chain acyl-CoA synthe
|
## |1088 |AT4G04040.1      |  2.87|t / c    |MEE51 &#124; Phosphofructokinase family
|
## |1138 |AT2G23670.1      |  2.87|t / c    |YCF37 &#124; homolog of Synechocystis Y
|
## |1422 |AT5G47930.1      |  2.87|t / c    |&#124; Zinc-binding ribosomal protein
|
## |739  |AT3G61470.1      |  2.86|t / c    |LHCA2 &#124; photosystem I light harves
|
## |421  |AT5G20280.1      |  2.85|t / c    |ATSPS1F, SPS1F &#124; sucrose phosphate
|
## |446  |AT3G07100.1      |  2.84|t / c    |ERMO2, SEC24A &#124; Sec23/Sec24 protei
|
## |520  |AT3G48750.1      |  2.84|t / c    |CDKA;1, CDC2AAT, CDK2, CDC2, CDC2A
|
## |633  |AT2G31810.1      |  2.83|t / c    |&#124; ACT domain-containing small sub
|
## |170  |AT4G35250.1      |  2.81|t / c    |&#124; NAD(P)-binding Rossmann-fold su
|
## |179  |AT5G50850.1      |  2.81|t / c    |MAB1 &#124; Transketolase family protei
|
## |384  |AT5G19550.1      |  2.81|t / c    |ASP2, AAT2 &#124; aspartate aminotransf
|
## |548  |AT5G24690.1      |  2.81|t / c    |&#124; Protein of unknown function (DU
|
## |628  |AT1G79600.1      |  2.81|t / c    |&#124; Protein kinase superfamily prot
|
## |698  |AT5G43900.3      |  2.81|t / c    |MYA2 &#124; myosin 2 &#124; chr5:17657241-1
7 |
```

```
## |827  |AT5G22330.1      |  2.81|t / c    |ATTIP49A, RIN1 &#124; P-loop containing
|
## |905  |AT2G20990.3      |  2.81|t / c    |SYTA &#124; synaptotagmin A &#124; chr2:901
4 |
## |921  |AT4G34830.1      |  2.81|t / c    |MRL1 &#124; Pentatricopeptide repeat (P
|
## |940  |AT5G23630.1      |  2.81|t / c    |PDR2, MIA &#124; phosphate deficiency r
|
## |1000 |AT1G21250.1      |  2.81|t / c    |WAK1, PRO25 &#124; cell wall-associated
|
## |1570 |AT1G16670.1      |  2.81|t / c    |&#124; Protein kinase superfamily prot
|
## |1060 |AT3G28710.1      |  2.81|t-unique |&#124; ATPase, V0/A0 complex, subunit
|
## |1077 |AT5G02890.1      |  2.81|t-unique |&#124; HXXXD-type acyl-transferase fam
|
## |284  |AT1G54780.1      |  2.80|t / c    |TLP18.3 &#124; thylakoid lumen 18.3 kDa
|
## |64   |AT2G30950.1      |  2.79|t / c    |VAR2, FTSH2 &#124; FtsH extracellular p
|
## |693  |AT4G28080.1      |  2.79|t / c    |&#124; Tetratricopeptide repeat (TPR)-
|
## |327  |AT5G03940.1      |  2.77|t / c    |FFC, 54CP, CPSRP54, SRP54CP &#124; chlo
|
## |495  |AT3G62530.1      |  2.77|t / c    |&#124; ARM repeat superfamily protein
|
## |351  |AT5G03880.1      |  2.76|t / c    |&#124; Thioredoxin family protein &#124; ch
|
## |105  |AT1G20010.1      |  2.75|t / c    |TUB5 &#124; tubulin beta-5 chain &#124; chr
1 |
## |149  |ATCG00500.1      |  2.75|t / c    |ACCD &#124; acetyl-CoA carboxylase carb
|
## |492  |AT3G01290.1      |  2.75|t / c    |&#124; SPFH/Band 7/PHB domain-containi
|
## |806  |AT4G13770.1      |  2.75|t / c    |CYP83A1, REF2 &#124; cytochrome P450, f
|
## |680  |AT5G20890.1      |  2.74|t / c    |&#124; TCP-1/cpn60 chaperonin family p
|
## |699  |AT3G23820.1      |  2.74|t / c    |GAE6 &#124; UDP-D-glucuronate 4-epimera
|
## |761  |ATCG00430.1      |  2.74|t / c    |PSBG &#124; photosystem II reaction cen
|
## |804  |AT4G23250.1      |  2.74|t / c    |EMB1290, DUF26-21, RKC1, CRK17 &#124; k
|
## |842  |AT5G02160.1      |  2.74|t / c    |&#124; unknown protein; FUNCTIONS IN:
|
## |975  |AT5G64740.1      |  2.74|t / c    |CESA6, IXR2, E112, PRC1 &#124; cellulos
|
## |1044 |AT3G23660.1      |  2.74|t / c    |&#124; Sec23/Sec24 protein transport f
|
## |1053 |AT5G03040.1 (+2) |  2.74|t / c    |iqd2 &#124; IQ-domain 2 &#124; chr5:710380-
7 |
```

```
## |1126 |AT3G63520.1      |  2.74|t / c    |CCD1, ATCCD1, ATNCED1, NCED1 &#124; car
|
## |1160 |AT1G63000.1      |  2.74|t / c    |NRS/ER, UER1 &#124; nucleotide-rhamnose
|
## |915  |AT5G58670.1      |  2.74|t-unique |ATPLC1, ATPLC, PLC1 &#124; phospholipas
|
## |1302 |AT3G47620.1      |  2.74|t-unique |AtTCP14, TCP14 &#124; TEOSINTE BRANCHED
|
## |123  |AT2G29550.1      |  2.71|t / c    |TUB7 &#124; tubulin beta-7 chain &#124; chr
2 |
## |2231 |AT4G39350.1      |  2.70|t-unique |CESA2, ATH-A, ATCESA2 &#124; cellulose
|
## |451  |AT1G18500.1      |  2.67|t / c    |MAML-4, IPMS1 &#124; methylthioalkylmal
|
## |805  |AT1G53500.1      |  2.67|t / c    |MUM4, RHM2, ATRHM2, ATMUM4 &#124; NAD-d
|
## |822  |AT1G11260.1      |  2.66|t-unique |STP1, ATSTP1 &#124; sugar transporter 1
|
## |1051 |AT4G22310.1      |  2.66|t-unique |&#124; Uncharacterised protein family
|
## |1207 |AT1G62640.1 (+1) |  2.66|t-unique |KAS III &#124; 3-ketoacyl-acyl carrier
|
## |1522 |AT1G75220.1      |  2.66|t-unique |&#124; Major facilitator superfamily p
|
## |118  |AT5G46290.1      |  2.66|t / c    |KASI, KAS1 &#124; 3-ketoacyl-acyl carri
|
## |570  |AT4G34090.1      |  2.66|t / c    |&#124; unknown protein; FUNCTIONS IN:
|
## |632  |AT3G62010.2      |  2.66|t / c    |&#124; unknown protein; LOCATED IN: ce
|
## |729  |AT2G29190.1 (+1) |  2.66|t / c    |APUM2, PUM2 &#124; pumilio 2 &#124; chr2:12
5 |
## |760  |AT4G38630.1      |  2.66|t / c    |RPN10, MCB1, ATMCB1, MBP1 &#124; regula
|
## |770  |AT1G71220.1 (+1) |  2.66|t / c    |EBS1, UGGT, PSL2 &#124; UDP-glucose:gly
|
## |882  |AT4G31500.1      |  2.66|t / c    |CYP83B1, SUR2, RNT1, RED1, ATR4 &#124;
|
## |908  |AT2G47240.1 (+1) |  2.66|t / c    |CER8, LACS1 &#124; AMP-dependent synthe
|
## |989  |AT5G22770.1 (+2) |  2.66|t / c    |alpha-ADR &#124; alpha-adaptin &#124; chr5:
7 |
## |1016 |AT1G19450.1      |  2.66|t / c    |&#124; Major facilitator superfamily p
|
## |1046 |AT1G74730.1      |  2.66|t / c    |&#124; Protein of unknown function (DU
|
## |1049 |AT4G08850.1      |  2.66|t / c    |&#124; Leucine-rich repeat receptor-li
|
## |1068 |AT1G01220.1      |  2.66|t / c    |FKGP, AtFKGP &#124; L-fucokinase/GDP-L-
|
## |1128 |AT1G08380.1      |  2.66|t / c    |PSAO &#124; photosystem I subunit O &#124;
c |
```

```
## |1266 |AT5G05200.1      |  2.66|t / c    |&#124; Protein kinase superfamily prot
|
## |1608 |AT4G12320.1      |  2.66|t / c    |CYP706A6 &#124; cytochrome P450, family
|
## |182  |AT5G58290.1      |  2.65|t / c    |RPT3 &#124; regulatory particle triple-
|
## |235  |AT1G70730.1      |  2.64|t / c    |PGM2 &#124; Phosphoglucomutase/phosphom
|
## |461  |AT4G35100.1 (+1) |  2.64|t / c    |PIP3, PIP3A, PIP2;7, SIMIP &#124; plasm
|
## |473  |AT2G33040.1      |  2.64|t / c    |ATP3 &#124; gamma subunit of Mt ATP syn
|
## |601  |AT1G73990.1      |  2.64|t / c    |SPPA, SPPA1 &#124; signal peptide pepti
|
## |135  |AT5G12250.1      |  2.63|t / c    |TUB6 &#124; beta-6 tubulin &#124; chr5:3961
3 |
## |131  |AT3G05530.1      |  2.62|t / c    |RPT5A, ATS6A.2 &#124; regulatory partic
|
## |106  |AT1G71500.1      |  2.60|t / c    |&#124; Rieske (2Fe-2S) domain-containi
|
## |121  |AT2G04030.1      |  2.60|t / c    |CR88, EMB1956, HSP90.5, Hsp88.1, A
|
## |296  |AT5G46110.3      |  2.60|t / c    |APE2, TPT &#124; Glucose-6-phosphate/ph
|
## |941  |AT1G01610.1      |  2.58|t-unique |ATGPAT4, GPAT4 &#124; glycerol-3-phosph
|
## |1204 |AT1G22710.1      |  2.58|t-unique |SUC2, SUT1, ATSUC2 &#124; sucrose-proto
|
## |353  |AT4G25080.4      |  2.58|t / c    |CHLM &#124; magnesium-protoporphyrin IX
|
## |420  |AT5G16070.1      |  2.58|t / c    |&#124; TCP-1/cpn60 chaperonin family p
|
## |689  |AT5G65620.1      |  2.58|t / c    |&#124; Zincin-like metalloproteases fa
|
## |813  |AT3G19960.2      |  2.58|t / c    |ATM1 &#124; myosin 1 &#124; chr3:6949787-69
5 |
## |826  |AT1G78830.1      |  2.58|t / c    |&#124; Curculin-like (mannose-binding)
|
## |851  |AT5G66470.1      |  2.58|t / c    |&#124; RNA binding;GTP binding &#124; chr5:
|
## |884  |AT1G29310.1      |  2.58|t / c    |&#124; SecY protein transport family p
|
## |917  |AT5G17020.1 (+1) |  2.58|t / c    |XPO1A, ATCRM1, ATXPO1, XPO1, HIT2
|
## |1045 |AT4G32410.1      |  2.58|t / c    |CESA1, RSW1, AtCESA1 &#124; cellulose s
|
## |1112 |AT5G59730.2      |  2.58|t / c    |ATEXO70H7, EXO70H7 &#124; exocyst subun
|
## |1165 |AT3G51550.1      |  2.58|t / c    |FER &#124; Malectin/receptor-like prote
|
## |1190 |AT3G20810.1 (+1) |  2.58|t / c    |JMJD5 &#124; 2-oxoglutarate (2OG) and F
|
```

```
## |1192 |AT3G25680.1      |  2.58|t / c    |&#124; FUNCTIONS IN: molecular_functio
|
## |1236 |AT1G07650.2      |  2.58|t / c    |&#124; Leucine-rich repeat transmembra
|
## |1267 |AT1G73650.2      |  2.58|t / c    |&#124; Protein of unknown function (DU
|
## |1340 |AT2G29140.1      |  2.58|t / c    |APUM3, PUM3 &#124; pumilio 3 &#124; chr2:12
5 |
## |1382 |AT1G22700.2      |  2.58|t / c    |&#124; Tetratricopeptide repeat (TPR)-
|
## |240  |AT1G45201.1      |  2.57|t / c    |ATTLL1, TLL1 &#124; triacylglycerol lip
|
## |252  |AT3G56940.1      |  2.57|t / c    |CRD1, CHL27, ACSF &#124; dicarboxylate
|
## |469  |AT3G63460.1      |  2.57|t / c    |&#124; transducin family protein / WD-
|
## |482  |AT3G18490.1      |  2.57|t / c    |&#124; Eukaryotic aspartyl protease fa
|
## |606  |AT2G42210.2      |  2.56|t / c    |ATOEP16-3, OEP16-3 &#124; Mitochondrial
|
## |110  |AT4G27440.1 (+1) |  2.55|t / c    |PORB &#124; protochlorophyllide oxidore
|
## |735  |AT1G06530.1      |  2.53|t / c    |&#124; Tropomyosin-related &#124; chr1:2001
|
## |313  |AT3G22890.1      |  2.52|t / c    |APS1 &#124; ATP sulfurylase 1 &#124; chr3:8
1 |
## |843  |AT3G51820.1      |  2.52|t / c    |ATG4, G4, CHLG &#124; UbiA prenyltransf
|
## |458  |AT3G58730.1      |  2.51|t / c    |&#124; vacuolar ATP synthase subunit D
|
## |732  |AT2G40840.1      |  2.50|t / c    |DPE2 &#124; disproportionating enzyme 2
|
## |780  |AT4G30950.1      |  2.50|t / c    |FAD6, FADC, SFD4 &#124; fatty acid desa
|
## |803  |AT4G16990.2      |  2.50|t / c    |RLM3 &#124; disease resistance protein
|
## |821  |AT1G55160.3      |  2.50|t / c    |&#124; unknown protein; FUNCTIONS IN:
|
## |954  |AT3G28860.1      |  2.50|t / c    |ATMDR1, ATMDR11, PGP19, MDR11, MDR
|
## |982  |AT5G60540.1      |  2.50|t / c    |EMB2407, ATPDX2, PDX2 &#124; pyridoxine
|
## |1004 |AT2G16950.1 (+1) |  2.50|t / c    |TRN1, ATTRN1 &#124; transportin 1 &#124; ch
r |
## |1024 |AT1G78915.1 (+2) |  2.50|t / c    |&#124; Tetratricopeptide repeat (TPR)-
|
## |1035 |AT2G23200.1      |  2.50|t / c    |&#124; Protein kinase superfamily prot
|
## |1110 |AT4G27700.1      |  2.50|t / c    |&#124; Rhodanese/Cell cycle control ph
|
## |1139 |AT4G04850.2      |  2.50|t / c    |KEA3 &#124; K+ efflux antiporter 3 &#124; c
h |
```

```
## |1291 |AT1G06000.1      |  2.50|t / c    |&#124; UDP-Glycosyltransferase superfa
|
## |1336 |AT5G21430.1      |  2.50|t / c    |&#124; Chaperone DnaJ-domain superfami
|
## |643  |AT5G11670.1      |  2.49|t / c    |ATNADP-ME2, NADP-ME2 &#124; NADP-malic
|
## |895  |AT4G33510.1      |  2.49|t / c    |DHS2 &#124; 3-deoxy-d-arabino-heptuloso
|
## |80   |AT1G74470.1      |  2.48|t / c    |&#124; Pyridine nucleotide-disulphide
|
## |145  |AT1G09130.1 (+1) |  2.48|t / c    |&#124; ATP-dependent caseinolytic (Clp
|
## |254  |AT2G33150.1      |  2.48|t / c    |PKT3, PED1, KAT2 &#124; peroxisomal 3-k
|
## |357  |AT5G52520.1      |  2.48|t / c    |OVA6, PRORS1 &#124; Class II aaRS and b
|
## |51   |AT5G42270.1      |  2.47|t / c    |VAR1, FTSH5 &#124; FtsH extracellular p
|
## |230  |AT1G45000.1      |  2.47|t / c    |&#124; AAA-type ATPase family protein
|
## |276  |AT3G58610.1 (+2) |  2.47|t / c    |&#124; ketol-acid reductoisomerase &#124; c
|
## |337  |AT1G52510.1      |  2.47|t / c    |&#124; alpha/beta-Hydrolases superfami
|
## |82   |AT3G54890.4      |  2.46|t / c    |LHCA1 &#124; photosystem I light harves
|
## |508  |AT4G20890.1      |  2.46|t / c    |TUB9 &#124; tubulin beta-9 chain &#124; chr
4 |
## |1153 |AT3G05910.1      |  2.46|t / c    |&#124; Pectinacetylesterase family pro
|
## |1452 |AT4G30610.1      |  2.46|t / c    |BRS1, SCPL24 &#124; alpha/beta-Hydrolas
|
## |1487 |AT3G03100.1      |  2.46|t / c    |&#124; NADH:ubiquinone oxidoreductase,
|
## |1973 |AT3G05000.1      |  2.46|t / c    |&#124; Transport protein particle (TRA
|
## |178  |AT5G58330.1 (+1) |  2.45|t / c    |&#124; lactate/malate dehydrogenase fa
|
## |344  |AT1G50250.1      |  2.44|t / c    |FTSH1 &#124; FTSH protease 1 &#124; chr1:18
6 |
## |448  |AT1G14810.1      |  2.44|t / c    |&#124; semialdehyde dehydrogenase fami
|
## |707  |AT5G62790.1      |  2.43|t / c    |DXR, PDE129 &#124; 1-deoxy-D-xylulose 5
|
## |382  |AT3G63410.1      |  2.42|t / c    |APG1, VTE3, IEP37, E37 &#124; S-adenosy
|
## |679  |AT5G45510.1 (+1) |  2.42|t / c    |&#124; Leucine-rich repeat (LRR) famil
|
## |746  |AT5G67630.1      |  2.42|t / c    |&#124; P-loop containing nucleoside tr
|
## |793  |AT3G62700.1      |  2.42|t / c    |ATMRP10, MRP10, ABCC14 &#124; multidrug
|
```

```
## |863  |AT4G26300.1      |  2.42|t / c    |emb1027 &#124; Arginyl-tRNA synthetase,
|
## |864  |AT3G25690.1 (+1) |  2.42|t / c    |CHUP1 &#124; Hydroxyproline-rich glycop
|
## |900  |AT4G24330.1      |  2.42|t / c    |&#124; Protein of unknown function (DU
|
## |914  |AT5G38990.1      |  2.42|t / c    |&#124; Malectin/receptor-like protein
|
## |932  |AT1G17580.1      |  2.42|t / c    |MYA1, ATMYA1, XI-1 &#124; myosin 1 &#124; c
h |
## |1023 |AT2G22125.1      |  2.42|t / c    |CSI1 &#124; binding &#124; chr2:9406793-941
4 |
## |1079 |AT2G38750.1      |  2.42|t / c    |ANNAT4 &#124; annexin 4 &#124; chr2:1619658
2 |
## |1086 |AT2G47390.1      |  2.42|t / c    |&#124; Prolyl oligopeptidase family pr
|
## |1137 |AT4G37000.1      |  2.42|t / c    |ACD2, ATRCCR &#124; accelerated cell de
|
## |1144 |AT1G15980.1      |  2.42|t / c    |NDF1, NDH48 &#124; NDH-dependent cyclic
|
## |1209 |AT1G53430.1 (+1) |  2.42|t / c    |&#124; Leucine-rich repeat transmembra
|
## |1227 |AT4G33220.1      |  2.42|t / c    |PME44, ATPME44 &#124; pectin methyleste
|
## |1304 |AT3G26210.1      |  2.42|t / c    |CYP71B23 &#124; cytochrome P450, family
|
## |1305 |AT2G34560.1      |  2.42|t / c    |&#124; P-loop containing nucleoside tr
|
## |2107 |AT1G21270.1      |  2.42|t-unique |WAK2 &#124; wall-associated kinase 2 &#124;
|
## |392  |AT1G13440.1      |  2.39|t / c    |GAPC-2, GAPC2 &#124; glyceraldehyde-3-p
|
## |462  |AT1G01320.2      |  2.37|t / c    |&#124; Tetratricopeptide repeat (TPR)-
|
## |1047 |AT1G05140.1      |  2.37|t / c    |&#124; Peptidase M50 family protein &#124;
|
## |22   |AT3G04120.1      |  2.36|t / c    |GAPC, GAPC-1, GAPC1 &#124; glyceraldehy
|
## |293  |AT2G19940.1 (+1) |  2.36|t / c    |&#124; oxidoreductases, acting on the
|
## |400  |AT4G34200.1      |  2.36|t / c    |EDA9 &#124; D-3-phosphoglycerate dehydr
|
## |691  |AT1G15730.1      |  2.35|t / c    |&#124; Cobalamin biosynthesis CobW-lik
|
## |868  |AT3G51160.1      |  2.35|t / c    |MUR1, MUR_1, GMD2 &#124; NAD(P)-binding
|
## |1175 |AT2G05710.1      |  2.35|t / c    |ACO3 &#124; aconitase 3 &#124; chr2:2141591
- |
## |268  |AT5G41670.1 (+1) |  2.34|t / c    |&#124; 6-phosphogluconate dehydrogenas
|
## |591  |AT2G32480.1      |  2.34|t / c    |ARASP &#124; ARABIDOPSIS SERIN PROTEASE
|
```

```
## |879  |AT3G48110.1      |  2.34|t / c    |EDD1, EDD &#124; glycine-tRNA ligases &#12
4; |
## |42   |AT1G04820.1 (+1) |  2.33|t / c    |TUA4, TOR2 &#124; tubulin alpha-4 chain
|
## |1202 |AT3G57650.1      |  2.32|t-unique |LPAT2 &#124; lysophosphatidyl acyltrans
|
## |1440 |AT4G15550.1      |  2.32|t-unique |IAGLU &#124; indole-3-acetate beta-D-gl
|
## |53   |AT5G09810.1      |  2.32|t / c    |ACT7 &#124; actin 7 &#124; chr5:3052809-305
4 |
## |521  |AT2G44160.1      |  2.32|t / c    |MTHFR2 &#124; methylenetetrahydrofolate
|
## |717  |AT4G03550.1      |  2.32|t / c    |ATGSL05, GSL05, ATGSL5, PMR4, GSL5
|
## |720  |AT1G29150.1      |  2.32|t / c    |ATS9, RPN6 &#124; non-ATPase subunit 9
|
## |755  |AT1G70320.1      |  2.32|t / c    |UPL2 &#124; ubiquitin-protein ligase 2
|
## |783  |AT4G23940.1      |  2.32|t / c    |&#124; FtsH extracellular protease fam
|
## |936  |AT2G45060.1      |  2.32|t / c    |&#124; Uncharacterised conserved prote
|
## |970  |AT5G47910.1      |  2.32|t / c    |RBOHD, ATRBOHD &#124; respiratory burst
|
## |1026 |AT1G79870.1      |  2.32|t / c    |&#124; D-isomer specific 2-hydroxyacid
|
## |1080 |AT3G42170.1      |  2.32|t / c    |&#124; BED zinc finger ;hAT family dim
|
## |1095 |AT4G24810.2      |  2.32|t / c    |&#124; Protein kinase superfamily prot
|
## |1219 |AT1G64710.1      |  2.32|t / c    |&#124; GroES-like zinc-binding dehydro
|
## |1229 |AT5G08530.1      |  2.32|t / c    |CI51 &#124; 51 kDa subunit of complex I
|
## |1380 |AT3G48140.1      |  2.32|t / c    |&#124; B12D protein &#124; chr3:17778471-17
|
## |1392 |AT2G35780.1      |  2.32|t / c    |scpl26 &#124; serine carboxypeptidase-l
|
## |1648 |AT5G58490.1      |  2.32|t / c    |&#124; NAD(P)-binding Rossmann-fold su
|
## |1766 |AT1G79990.1      |  2.32|t / c    |&#124; structural molecules &#124; chr1:300
|
## |1810 |AT3G22750.1      |  2.32|t / c    |&#124; Protein kinase superfamily prot
|
## |91   |AT5G23060.1      |  2.31|t / c    |CaS &#124; calcium sensing receptor &#124;
c |
## |661  |AT3G52730.1      |  2.30|t / c    |&#124; ubiquinol-cytochrome C reductas
|
## |762  |AT3G19480.1      |  2.30|t / c    |&#124; D-3-phosphoglycerate dehydrogen
|
## |19   |AT1G49240.1      |  2.29|t / c    |ACT8 &#124; actin 8 &#124; chr1:18216539-18
2 |
```

```
## |50   |AT3G08940.2      |  2.29|t / c    |LHCB4.2 &#124; light harvesting complex
|
## |188  |AT5G19770.1 (+1) |  2.29|t / c    |TUA3 &#124; tubulin alpha-3 &#124; chr5:668
2 |
## |869  |AT5G12860.1 (+1) |  2.29|t / c    |DiT1 &#124; dicarboxylate transporter 1
|
## |4    |AT2G39730.1      |  2.28|t / c    |RCA &#124; rubisco activase &#124; chr2:165
7 |
## |356  |AT2G44640.1      |  2.28|t / c    |&#124; FUNCTIONS IN: molecular_functio
|
## |185  |AT3G19170.1      |  2.27|t / c    |ATPREP1, ATZNMP, PREP1 &#124; presequen
|
## |502  |AT5G40770.1      |  2.27|t / c    |ATPHB3, PHB3 &#124; prohibitin 3 &#124; chr
5 |
## |261  |AT2G39730.2      |  2.26|t / c    |RCA &#124; rubisco activase &#124; chr2:165
7 |
## |478  |AT4G01800.1      |  2.26|t / c    |AGY1, AtcpSecA, SECA1 &#124; Albino or
|
## |583  |AT1G06430.1      |  2.26|t / c    |FTSH8 &#124; FTSH protease 8 &#124; chr1:19
6 |
## |263  |AT1G51980.1      |  2.25|t / c    |&#124; Insulinase (Peptidase family M1
|
## |724  |AT1G01080.2      |  2.25|t / c    |&#124; RNA-binding (RRM/RBD/RNP motifs
|
## |834  |AT1G19920.1      |  2.25|t / c    |APS2, ASA1 &#124; Pseudouridine synthas
|
## |959  |AT1G14150.1      |  2.25|t / c    |PQL1, PQL2 &#124; PsbQ-like 2 &#124; chr1:4
8 |
## |1277 |AT1G61790.1      |  2.22|t-unique |&#124; Oligosaccharyltransferase compl
|
## |1300 |AT4G17770.1      |  2.22|t-unique |ATTPS5, TPS5 &#124; trehalose phosphata
|
## |711  |AT3G61050.1 (+1) |  2.22|t / c    |NTMC2TYPE4, NTMC2T4 &#124; Calcium-depe
|
## |816  |AT2G20760.1      |  2.22|t / c    |&#124; Clathrin light chain protein &#124;
|
## |819  |AT4G24620.1      |  2.22|t / c    |PGI1, PGI &#124; phosphoglucose isomera
|
## |865  |AT3G04340.1      |  2.22|t / c    |emb2458 &#124; FtsH extracellular prote
|
## |995  |AT4G34350.1      |  2.22|t / c    |CLB6, ISPH, HDR &#124; 4-hydroxy-3-meth
|
## |1014 |AT2G27860.1      |  2.22|t / c    |AXS1 &#124; UDP-D-apiose/UDP-D-xylose s
|
## |1021 |AT2G25800.1      |  2.22|t / c    |&#124; Protein of unknown function (DU
|
## |1022 |AT4G36220.1      |  2.22|t / c    |FAH1, CYP84A1 &#124; ferulic acid 5-hyd
|
## |1025 |AT1G47550.1 (+1) |  2.22|t / c    |SEC3A &#124; exocyst complex component
|
## |1073 |AT1G12770.1      |  2.22|t / c    |ISE1, EMB1586 &#124; P-loop containing
|
```

```
## |1083 |AT2G20920.1      |  2.22|t / c    |&#124; Protein of unknown function (DU
|
## |1230 |AT3G53180.1      |  2.22|t / c    |&#124; glutamate-ammonia ligases;catal
|
## |1349 |AT5G35590.1      |  2.22|t / c    |PAA1 &#124; proteasome alpha subunit A1
|
## |1519 |AT4G36480.1 (+1) |  2.22|t / c    |ATLCB1, LCB1, EMB2779, FBR11 &#124; lon
|
## |1577 |AT3G51890.1      |  2.22|t / c    |&#124; Clathrin light chain protein &#124;
|
## |1675 |AT1G14930.1      |  2.22|t / c    |&#124; Polyketide cyclase/dehydrase an
|
## |662  |AT1G16720.1      |  2.21|t / c    |HCF173 &#124; high chlorophyll fluoresc
|
## |213  |ATCG01110.1      |  2.20|t / c    |NDHH &#124; NAD(P)H dehydrogenase subun
|
## |667  |AT3G22960.1      |  2.20|t / c    |PKP1, PKP-ALPHA &#124; Pyruvate kinase
|
## |214  |AT1G79040.1      |  2.19|t / c    |PSBR &#124; photosystem II subunit R &#124;
|
## |320  |AT1G20200.1      |  2.19|t / c    |EMB2719, HAP15 &#124; PAM domain (PCI/P
|
## |432  |AT1G11750.1      |  2.19|t / c    |CLPP6, NCLPP1, NCLPP6 &#124; CLP protea
|
## |592  |AT3G61440.1      |  2.19|t / c    |ATCYSC1, ARATH;BSAS3;1, CYSC1 &#124; cy
|
## |1168 |AT5G23860.1 (+1) |  2.19|t / c    |TUB8 &#124; tubulin beta 8 &#124; chr5:8042
9 |
## |404  |AT3G56150.1 (+1) |  2.18|t / c    |EIF3C, ATEIF3C-1, EIF3C-1, ATTIF3C
|
## |523  |AT3G06510.2      |  2.18|t / c    |SFR2 &#124; Glycosyl hydrolase superfam
|
## |1159 |AT2G07707.1 (+1) |  2.18|t / c    |&#124; Plant mitochondrial ATPase, F0
|
## |21   |AT3G14420.1 (+1) |  2.17|t / c    |&#124; Aldolase-type TIM barrel family
|
## |168  |AT1G62750.1      |  2.17|t / c    |ATSCO1, ATSCO1/CPEF-G, SCO1 &#124; Tran
|
## |586  |AT5G13650.2      |  2.17|t / c    |&#124; elongation factor family protei
|
## |828  |AT4G21150.1 (+1) |  2.17|t / c    |HAP6 &#124; ribophorin II (RPN2) family
|
## |873  |AT4G39080.1      |  2.17|t / c    |VHA-A3 &#124; vacuolar proton ATPase A3
|
## |911  |AT1G76180.1 (+1) |  2.17|t / c    |ERD14 &#124; Dehydrin family protein &#124;
|
## |1244 |AT1G13060.1      |  2.17|t / c    |PBE1 &#124; 20S proteasome beta subunit
|
## |1286 |AT4G28740.1      |  2.17|t / c    |&#124; FUNCTIONS IN: molecular_functio
|
## |2066 |AT3G16050.1      |  2.17|t / c    |A37, ATPDX1.2, PDX1.2 &#124; pyridoxine
|
```

```
## |333  |AT1G03630.2      |  2.16|t / c    |POR C, PORC &#124; protochlorophyllide
|
## |1102 |AT3G22845.1      |  2.15|t / c    |&#124; emp24/gp25L/p24 family/GOLD fam
|
## |32   |AT4G04640.1      |  2.14|t / c    |ATPC1 &#124; ATPase, F1 complex, gamma
|
## |364  |AT2G37620.1 (+2) |  2.14|t / c    |ACT1, AAc1 &#124; actin 1 &#124; chr2:15779
7 |
## |524  |AT4G08870.1      |  2.14|t / c    |&#124; Arginase/deacetylase superfamil
|
## |543  |AT1G52360.1      |  2.14|t / c    |&#124; Coatomer, beta' subunit &#124; chr1:
|
## |474  |AT3G44110.1      |  2.13|t / c    |ATJ3, ATJ &#124; DNAJ homologue 3 &#124; ch
r |
## |893  |AT5G49030.3      |  2.12|t / c    |OVA2 &#124; tRNA synthetase class I (I,
|
## |968  |AT5G64580.1      |  2.12|t / c    |&#124; AAA-type ATPase family protein
|
## |969  |AT2G43950.1      |  2.12|t / c    |OEP37, ATOEP37 &#124; chloroplast outer
|
## |1069 |AT1G08930.1 (+1) |  2.12|t / c    |ERD6 &#124; Major facilitator superfami
|
## |1081 |AT3G49720.1 (+1) |  2.12|t / c    |&#124; unknown protein; FUNCTIONS IN:
|
## |1087 |AT2G05840.1      |  2.12|t / c    |PAA2 &#124; 20S proteasome subunit PAA2
|
## |1178 |AT3G23300.1      |  2.12|t / c    |&#124; S-adenosyl-L-methionine-depende
|
## |1238 |AT1G09795.1      |  2.12|t / c    |ATATP-PRT2, HISN1B, ATP-PRT2 &#124; ATP
|
## |1243 |AT5G58100.1      |  2.12|t / c    |&#124; unknown protein; INVOLVED IN: p
|
## |1254 |AT2G44530.2      |  2.12|t / c    |&#124; Phosphoribosyltransferase famil
|
## |1255 |AT1G29790.1 (+1) |  2.12|t / c    |&#124; S-adenosyl-L-methionine-depende
|
## |1329 |AT3G51420.1      |  2.12|t / c    |SSL4, ATSSL4 &#124; strictosidine synth
|
## |1365 |AT5G11450.1      |  2.12|t / c    |&#124; Mog1/PsbP/DUF1795-like photosys
|
## |1399 |AT5G47200.1      |  2.12|t / c    |ATRABD2B, ATRAB1A, RAB1A &#124; RAB GTP
|
## |1416 |AT3G02350.1      |  2.12|t / c    |GAUT9 &#124; galacturonosyltransferase
|
## |1710 |AT3G54470.1      |  2.12|t / c    |&#124; uridine 5'-monophosphate syntha
|
## |2035 |AT4G00360.1      |  2.12|t-unique |CYP86A2, ATT1 &#124; cytochrome P450, f
|
## |47   |AT2G07698.1      |  2.10|t / c    |&#124; ATPase, F1 complex, alpha subun
|
## |747  |AT3G01310.2      |  2.09|t / c    |&#124; Phosphoglycerate mutase-like fa
|
```

```
## |829  |AT1G04530.1      |  2.09|t / c    |TPR4 &#124; Tetratricopeptide repeat (T
|
## |1010 |AT5G08650.1      |  2.09|t / c    |&#124; Small GTP-binding protein &#124; chr
|
## |71   |AT3G47470.1      |  2.08|t / c    |LHCA4, CAB4 &#124; light-harvesting chl
|
## |222  |AT3G63160.1      |  2.08|t / c    |&#124; FUNCTIONS IN: molecular_functio
|
## |380  |AT4G11420.1      |  2.08|t / c    |EIF3A, ATEIF3A-1, EIF3A-1, ATTIF3A
|
## |948  |AT2G21960.1      |  2.08|t / c    |&#124; unknown protein; LOCATED IN: ch
|
## |951  |AT2G27730.1      |  2.08|t / c    |&#124; copper ion binding &#124; chr2:11820
|
## |1061 |AT5G64290.1      |  2.08|t / c    |DCT, DIT2.1 &#124; dicarboxylate transp
|
## |155  |AT3G53420.1 (+1) |  2.07|t / c    |PIP2A, PIP2, PIP2;1 &#124; plasma membr
|
## |361  |AT3G04790.1      |  2.07|t / c    |&#124; Ribose 5-phosphate isomerase, t
|
## |646  |AT1G09620.1      |  2.07|t / c    |&#124; ATP binding;leucine-tRNA ligase
|
## |727  |AT3G11710.1      |  2.07|t / c    |ATKRS-1 &#124; lysyl-tRNA synthetase 1
|
## |90   |AT1G23310.1      |  2.06|t / c    |GGT1, AOAT1, GGAT1 &#124; glutamate:gly
|
## |647  |AT3G46060.1 (+2) |  2.06|t / c    |ARA3, ARA-3, ATRABE1C, ATRAB8A, RA
|
## |703  |AT1G80030.1 (+2) |  2.06|t / c    |&#124; Molecular chaperone Hsp40/DnaJ
|
## |794  |AT4G24820.1 (+1) |  2.06|t / c    |&#124; 26S proteasome, regulatory subu
|
## |1201 |AT1G72730.1      |  2.06|t / c    |&#124; DEA(D/H)-box RNA helicase famil
|
## |1309 |AT2G47110.1 (+1) |  2.06|t / c    |UBQ6 &#124; ubiquitin 6 &#124; chr2:1934470
1 |
## |696  |AT5G51070.1      |  2.05|t / c    |ERD1, CLPD, SAG15 &#124; Clp ATPase &#124;
c |
## |1827 |AT3G15980.1 (+3) |  2.05|t / c    |&#124; Coatomer, beta' subunit &#124; chr3:
|
## |393  |AT5G63570.1      |  2.04|t / c    |GSA1 &#124; glutamate-1-semialdehyde-2,
|
## |620  |ATCG00420.1      |  2.04|t / c    |NDHJ &#124; NADH dehydrogenase subunit
|
## |1091 |AT3G48730.1      |  2.04|t / c    |GSA2 &#124; glutamate-1-semialdehyde 2,
|
## |379  |AT3G15730.1      |  2.03|t / c    |PLDALPHA1, PLD &#124; phospholipase D a
|
## |259  |AT2G38230.1      |  2.02|t / c    |ATPDX1.1, PDX1.1 &#124; pyridoxine bios
|
## |635  |AT3G12110.1      |  2.02|t / c    |ACT11 &#124; actin-11 &#124; chr3:3858116-3
8 |
```

```
## |243  |AT4G03280.1      |  2.01|t / c    |PETC, PGR1 &#124; photosynthetic electr
|
## |480  |AT1G02560.1      |  2.00|t / c    |CLPP5, NCLPP5, NCLPP1 &#124; nuclear en
|
## |518  |AT2G10940.1 (+1) |  2.00|t / c    |&#124; Bifunctional inhibitor/lipid-tr
|
## |546  |AT2G22250.2 (+1) |  2.00|t / c    |ATAAT, AAT, MEE17 &#124; aspartate amin
|
## |637  |AT2G39770.1 (+1) |  2.00|t / c    |CYT1, VTC1, SOZ1, EMB101, GMP1 &#124; G
|
## |810  |AT3G20050.1      |  2.00|t / c    |ATTCP-1, TCP-1 &#124; T-complex protein
|
## |848  |AT5G49810.1      |  2.00|t / c    |MMT &#124; methionine S-methyltransfera
|
## |855  |AT4G25130.1      |  2.00|t / c    |PMSR4 &#124; peptide met sulfoxide redu
|
## |909  |AT3G18190.1      |  2.00|t / c    |&#124; TCP-1/cpn60 chaperonin family p
|
## |939  |AT3G01440.1      |  2.00|t / c    |PQL1, PQL2 &#124; PsbQ-like 1 &#124; chr3:1
6 |
## |958  |AT2G36810.1      |  2.00|t / c    |&#124; ARM repeat superfamily protein
|
## |996  |AT3G11830.1      |  2.00|t / c    |&#124; TCP-1/cpn60 chaperonin family p
|
## |1103 |AT1G26850.1 (+1) |  2.00|t / c    |&#124; S-adenosyl-L-methionine-depende
|
## |1135 |AT5G63510.2      |  2.00|t / c    |GAMMA CAL1 &#124; gamma carbonic anhydr
|
## |1191 |AT4G38580.1      |  2.00|t / c    |ATFP6, HIPP26, FP6 &#124; farnesylated
|
## |1260 |AT2G01350.1      |  2.00|t / c    |QPT &#124; quinolinate phoshoribosyltra
|
## |1265 |AT4G33050.3      |  2.00|t-unique |EDA39 &#124; calmodulin-binding family
|
## |1303 |AT3G09090.1 (+1) |  2.00|t / c    |DEX1 &#124; defective in exine formatio
|
## |1308 |AT1G51660.1      |  2.00|t / c    |ATMKK4, MKK4, ATMEK4 &#124; mitogen-act
|
## |1334 |AT3G57280.1      |  2.00|t / c    |&#124; Transmembrane proteins 14C &#124; ch
|
## |1385 |AT4G30920.1      |  2.00|t / c    |&#124; Cytosol aminopeptidase family p
|
## |1396 |AT1G74910.1 (+1) |  2.00|t / c    |&#124; ADP-glucose pyrophosphorylase f
|
## |1428 |AT5G61910.4      |  2.00|t / c    |&#124; DCD (Development and Cell Death
|
## |1429 |AT1G26550.1      |  2.00|t / c    |&#124; FKBP-like peptidyl-prolyl cis-t
|
## |1439 |AT3G25070.1      |  2.00|t-unique |RIN4 &#124; RPM1 interacting protein 4
|
## |1467 |AT4G24220.1 (+1) |  2.00|t-unique |VEP1, AWI31 &#124; NAD(P)-binding Rossm
|
```

```
## |1472 |AT4G32590.1      |  2.00|t / c    |&#124; 2Fe-2S ferredoxin-like superfam
|
## |1506 |AT1G50430.1 (+1) |  2.00|t / c    |DWF5, PA, LE, ST7R, 7RED &#124; Ergoste
|
## |1518 |AT3G44620.1 (+1) |  2.00|t-unique |&#124; protein tyrosine phosphatases;p
|
## |1529 |AT5G22640.1      |  2.00|t / c    |emb1211 &#124; MORN (Membrane Occupatio
|
## |1654 |AT5G17520.1      |  2.00|t-unique |RCP1, MEX1 &#124; root cap 1 (RCP1) &#124;
c |
## |1773 |AT1G30440.1      |  2.00|t / c    |&#124; Phototropic-responsive NPH3 fam
|
## |1814 |AT2G30490.1      |  2.00|t / c    |ATC4H, C4H, CYP73A5, REF3 &#124; cinnam
|
## |2043 |AT1G32220.1      |  2.00|t-unique |&#124; NAD(P)-binding Rossmann-fold su
|
## |33   |AT1G61520.1 (+1) |  1.98|t / c    |LHCA3 &#124; photosystem I light harves
|
## |395  |AT1G32500.1      |  1.97|t / c    |ATNAP6, NAP6 &#124; non-intrinsic ABC p
|
## |181  |AT2G47730.1      |  1.96|t / c    |ATGSTF8, ATGSTF5, GST6, GSTF8 &#124; gl
|
## |305  |AT1G29900.1      |  1.96|t / c    |CARB &#124; carbamoyl phosphate synthet
|
## |535  |AT3G46740.1      |  1.96|t / c    |TOC75-III, MAR1 &#124; translocon at th
|
## |600  |AT5G28840.1 (+1) |  1.96|t / c    |GME &#124; GDP-D-mannose 3',5'-epimeras
|
## |136  |ATCG00800.1      |  1.95|t / c    |&#124; structural constituent of ribos
|
## |173  |AT3G14415.1 (+1) |  1.95|t / c    |&#124; Aldolase-type TIM barrel family
|
## |374  |AT4G01100.2      |  1.95|t / c    |ADNT1 &#124; adenine nucleotide transpo
|
## |640  |AT5G42240.1      |  1.95|t / c    |scpl42 &#124; serine carboxypeptidase-l
|
## |48   |AT3G62250.1      |  1.94|t / c    |UBQ5 &#124; ubiquitin 5 &#124; chr3:2303713
8 |
## |102  |ATCG00130.1      |  1.94|t / c    |ATPF &#124; ATPase, F0 complex, subunit
|
## |340  |AT4G31990.1 (+2) |  1.93|t / c    |ASP5, AAT3, ATAAT1 &#124; aspartate ami
|
## |453  |AT3G02360.1 (+1) |  1.93|t / c    |&#124; 6-phosphogluconate dehydrogenas
|
## |501  |AT4G11150.1      |  1.92|t / c    |TUF, emb2448, TUFF, VHA-E1 &#124; vacuo
|
## |514  |AT5G55190.1      |  1.92|t / c    |RAN3, ATRAN3 &#124; RAN GTPase 3 &#124; chr
5 |
## |561  |AT4G39710.2      |  1.92|t / c    |FKBP16-2 &#124; FK506-binding protein 1
|
## |953  |AT4G29840.1      |  1.92|t / c    |MTO2, TS &#124; Pyridoxal-5'-phosphate-
|
```

```
## |1063 |AT5G45280.2      |  1.92|t / c    |&#124; Pectinacetylesterase family pro
|
## |1890 |AT5G43780.1      |  1.92|t / c    |APS4 &#124; Pseudouridine synthase/arch
|
## |398  |AT4G02510.1      |  1.91|t / c    |TOC159, TOC86, PPI2, TOC160, ATTOC
|
## |484  |AT3G06650.1      |  1.91|t / c    |ACLB-1 &#124; ATP-citrate lyase B-1 &#124;
c |
## |678  |AT5G13430.1      |  1.91|t / c    |&#124; Ubiquinol-cytochrome C reductas
|
## |1020 |AT1G48520.1      |  1.91|t / c    |GATB &#124; GLU-ADT subunit B &#124; chr1:1
7 |
## |257  |AT3G54050.1 (+1) |  1.90|t / c    |HCEF1 &#124; high cyclic electron flow
|
## |605  |AT1G51805.1      |  1.90|t / c    |&#124; Leucine-rich repeat protein kin
|
## |881  |AT1G23190.1      |  1.90|t / c    |PGM3 &#124; Phosphoglucomutase/phosphom
|
## |44   |AT5G50920.1      |  1.89|t / c    |CLPC, ATHSP93-V, HSP93-V, DCA1, CL
|
## |300  |AT1G30380.1      |  1.89|t / c    |PSAK &#124; photosystem I subunit K &#124;
c |
## |439  |AT3G48870.1      |  1.89|t / c    |ATCLPC, ATHSP93-III, HSP93-III &#124; C
|
## |955  |AT4G14960.2      |  1.89|t / c    |TUA6 &#124; Tubulin/FtsZ family protein
|
## |138  |AT3G02090.1      |  1.88|t / c    |MPPBETA &#124; Insulinase (Peptidase fa
|
## |486  |AT1G65960.2      |  1.88|t / c    |GAD2 &#124; glutamate decarboxylase 2 &#12
4; |
## |1864 |AT2G31040.1      |  1.87|t-unique |&#124; ATP synthase protein I -related
|
## |2024 |AT1G32050.1      |  1.87|t-unique |&#124; SCAMP family protein &#124; chr1:115
|
## |771  |AT1G30400.1 (+1) |  1.87|t / c    |ATMRP1, EST1, ABCC1, ATABCC1, MRP1
|
## |775  |AT1G14610.1      |  1.87|t / c    |TWN2, VALRS &#124; valyl-tRNA synthetas
|
## |856  |AT3G20790.1      |  1.87|t / c    |&#124; NAD(P)-binding Rossmann-fold su
|
## |867  |AT1G72750.1      |  1.87|t / c    |ATTIM23-2, TIM23-2 &#124; translocase i
|
## |874  |AT1G02080.1      |  1.87|t / c    |&#124; transcription regulators &#124; chr1
|
## |916  |AT5G58140.1 (+2) |  1.87|t / c    |PHOT2, NPL1 &#124; phototropin 2 &#124; chr
5 |
## |950  |AT2G38670.1      |  1.87|t / c    |PECT1 &#124; phosphorylethanolamine cyt
|
## |1001 |AT2G24820.1      |  1.87|t / c    |TIC55-II &#124; translocon at the inner
|
## |1038 |AT1G63770.3      |  1.87|t / c    |&#124; Peptidase M1 family protein &#124; c
|
```

```
## |1101 |AT4G36250.1      |  1.87|t / c    |ALDH3F1 &#124; aldehyde dehydrogenase 3
|
## |1120 |AT2G24180.1      |  1.87|t / c    |CYP71B6 &#124; cytochrome p450 71b6 &#124;
c |
## |1212 |AT1G16300.1      |  1.87|t / c    |GAPCP-2 &#124; glyceraldehyde-3-phospha
|
## |1253 |AT2G18330.1      |  1.87|t / c    |&#124; AAA-type ATPase family protein
|
## |1256 |AT4G01690.1      |  1.87|t / c    |PPOX, HEMG1, PPO1 &#124; Flavin contain
|
## |1285 |AT5G16660.2 (+1) |  1.87|t / c    |&#124; unknown protein; FUNCTIONS IN:
|
## |1296 |AT2G36850.1      |  1.87|t / c    |ATGSL08, GSL8, GSL08, ATGSL8, CHOR
|
## |1374 |AT5G11040.1      |  1.87|t / c    |TRS120, AtTRS120 &#124; TRS120 &#124; chr5:
3 |
## |1391 |AT5G26830.1      |  1.87|t / c    |&#124; Threonyl-tRNA synthetase &#124; chr5
|
## |1404 |AT2G12190.1      |  1.87|t / c    |&#124; Cytochrome P450 superfamily pro
|
## |1406 |AT1G31800.1      |  1.87|t / c    |CYP97A3, LUT5 &#124; cytochrome P450, f
|
## |1415 |AT4G36390.1      |  1.87|t / c    |&#124; Methylthiotransferase &#124; chr4:17
|
## |1426 |AT4G28710.1      |  1.87|t / c    |XIH, ATXIH &#124; Myosin family protein
|
## |1482 |AT3G15660.1 (+1) |  1.87|t / c    |ATGRX4, GRX4 &#124; glutaredoxin 4 &#124; c
h |
## |1600 |AT1G14670.1      |  1.87|t / c    |&#124; Endomembrane protein 70 protein
|
## |1645 |AT1G56200.1      |  1.87|t / c    |emb1303 &#124; embryo defective 1303 &#124;
|
## |1647 |AT2G28430.1      |  1.87|t / c    |&#124; unknown protein; Has 28 Blast h
|
## |1972 |AT2G25870.1      |  1.87|t / c    |&#124; haloacid dehalogenase-like hydr
|
## |593  |AT2G35840.1 (+2) |  1.86|t / c    |&#124; Sucrose-6F-phosphate phosphohyd
|
## |792  |AT1G12000.1      |  1.86|t / c    |&#124; Phosphofructokinase family prot
|
## |115  |AT1G76030.1      |  1.85|t / c    |&#124; ATPase, V1 complex, subunit B p
|
## |999  |AT5G42390.1      |  1.85|t / c    |&#124; Insulinase (Peptidase family M1
|
## |1872 |AT1G75990.1      |  1.85|t / c    |&#124; PAM domain (PCI/PINT associated
|
## |208  |AT5G35360.3      |  1.83|t / c    |CAC2 &#124; acetyl Co-enzyme a carboxyl
|
## |1124 |AT1G71810.1      |  1.83|t / c    |&#124; Protein kinase superfamily prot
|
## |27   |AT4G10340.1      |  1.82|t / c    |LHCB5 &#124; light harvesting complex o
|
```

```
## |174  |AT1G48030.1 (+1) |  1.82|t / c    |mtLPD1 &#124; mitochondrial lipoamide d
|
## |611  |AT5G63420.1      |  1.81|t / c    |emb2746 &#124; RNA-metabolising metallo
|
## |665  |AT3G42050.1      |  1.81|t / c    |&#124; vacuolar ATP synthase subunit H
|
## |1262 |AT3G44330.1      |  1.81|t / c    |&#124; INVOLVED IN: protein processing
|
## |1328 |AT1G04170.1      |  1.81|t / c    |EIF2 GAMMA &#124; eukaryotic translatio
|
## |1633 |AT4G19490.1 (+1) |  1.81|t / c    |ATVPS54, VPS54 &#124; VPS54 &#124; chr4:106
1 |
## |1738 |AT1G72170.1      |  1.81|t / c    |&#124; Domain of unknown function (DUF
|
## |1940 |AT5G18230.1      |  1.81|t / c    |&#124; transcription regulator NOT2/NO
|
## |1976 |AT5G04530.1      |  1.81|t / c    |KCS19 &#124; 3-ketoacyl-CoA synthase 19
|
## |2120 |AT5G56760.1      |  1.81|t-unique |ATSERAT1;1, SAT5, SAT-52, SERAT1;1
|
## |239  |AT5G42650.1      |  1.80|t / c    |AOS, CYP74A, DDE2 &#124; allene oxide s
|
## |815  |AT5G64050.1      |  1.80|t / c    |ATERS, OVA3, ERS &#124; glutamate tRNA
|
## |1222 |AT5G15490.1      |  1.80|t / c    |&#124; UDP-glucose 6-dehydrogenase fam
|
## |197  |AT1G12900.1      |  1.79|t / c    |GAPA-2 &#124; glyceraldehyde 3-phosphat
|
## |406  |AT1G49970.1      |  1.79|t / c    |CLPR1, NCLPP5, SVR2 &#124; CLP protease
|
## |88   |AT5G25980.2      |  1.78|t / c    |TGG2, BGLU37 &#124; glucoside glucohydr
|
## |10   |ATCG00120.1      |  1.77|t / c    |ATPA &#124; ATP synthase subunit alpha
|
## |355  |AT5G65010.2      |  1.77|t / c    |ASN2 &#124; asparagine synthetase 2 &#124;
c |
## |58   |AT3G13920.1      |  1.76|t / c    |EIF4A1, RH4, TIF4A1 &#124; eukaryotic t
|
## |129  |ATCG00470.1      |  1.76|t / c    |ATPE &#124; ATP synthase epsilon chain
|
## |56   |AT1G56070.1      |  1.75|t / c    |LOS1 &#124; Ribosomal protein S5/Elonga
|
## |345  |AT1G62020.1      |  1.74|t / c    |&#124; Coatomer, alpha subunit &#124; chr1:
|
## |706  |AT5G45390.1      |  1.74|t / c    |CLPP4, NCLPP4 &#124; CLP protease P4 &#124;
|
## |880  |AT5G15450.1      |  1.74|t / c    |APG6, CLPB3, CLPB-P &#124; casein lytic
|
## |1074 |AT3G10690.1      |  1.74|t / c    |GYRA &#124; DNA GYRASE A &#124; chr3:333961
2 |
## |1085 |AT3G62360.1      |  1.74|t / c    |&#124; Carbohydrate-binding-like fold
|
```

```
## |1092 |AT5G14950.1      |  1.74|t / c    |GMII, ATGMII &#124; golgi alpha-mannosi
|
## |1093 |ATCG00170.1      |  1.74|t / c    |RPOC2 &#124; DNA-directed RNA polymeras
|
## |1111 |AT3G13330.1      |  1.74|t / c    |PA200 &#124; proteasome activating prot
|
## |1131 |AT2G28800.1 (+1) |  1.74|t / c    |ALB3 &#124; 63 kDa inner membrane famil
|
## |1164 |AT5G14060.1 (+1) |  1.74|t / c    |CARAB-AK-LYS &#124; Aspartate kinase fa
|
## |1228 |AT3G51260.1      |  1.74|t / c    |PAD1 &#124; 20S proteasome  alpha subun
|
## |1237 |AT4G00026.1      |  1.74|t / c    |&#124; FUNCTIONS IN: molecular_functio
|
## |1239 |AT2G35800.1      |  1.74|t / c    |&#124; mitochondrial substrate carrier
|
## |1339 |AT3G07700.1 (+2) |  1.74|t / c    |&#124; Protein kinase superfamily prot
|
## |1363 |AT4G32250.1 (+2) |  1.74|t / c    |&#124; Protein kinase superfamily prot
|
## |1376 |AT3G62150.1      |  1.74|t / c    |PGP21 &#124; P-glycoprotein 21 &#124; chr3:
2 |
## |1383 |AT1G50370.1      |  1.74|t / c    |&#124; Calcineurin-like metallo-phosph
|
## |1476 |AT5G67560.1      |  1.74|t / c    |ATARLA1D, ARLA1D &#124; ADP-ribosylatio
|
## |1520 |AT5G23300.1      |  1.74|t / c    |PYRD &#124; pyrimidine d &#124; chr5:784779
2 |
## |1544 |AT5G59250.1      |  1.74|t / c    |&#124; Major facilitator superfamily p
|
## |1547 |AT3G24530.1      |  1.74|t / c    |&#124; AAA-type ATPase family protein
|
## |1784 |AT1G64900.1      |  1.74|t / c    |CYP89A2, CYP89 &#124; cytochrome P450,
|
## |1839 |AT5G11480.1      |  1.74|t-unique |&#124; P-loop containing nucleoside tr
|
## |1929 |AT5G55710.1      |  1.74|t-unique |&#124; FUNCTIONS IN: molecular_functio
|
## |1966 |AT5G10780.1 (+1) |  1.74|t-unique |&#124; CONTAINS InterPro DOMAIN/s: Unc
|
## |28   |AT3G26650.1      |  1.72|t / c    |GAPA, GAPA-1 &#124; glyceraldehyde 3-ph
|
## |211  |AT5G09660.2      |  1.72|t / c    |PMDH2 &#124; peroxisomal NAD-malate deh
|
## |424  |AT1G08520.1      |  1.72|t / c    |ALB1, ALB-1V, V157, PDE166, CHLD &#124;
|
## |608  |AT3G03960.1      |  1.72|t / c    |&#124; TCP-1/cpn60 chaperonin family p
|
## |1050 |AT2G32060.1 (+2) |  1.72|t / c    |&#124; Ribosomal protein L7Ae/L30e/S12
|
## |306  |AT1G34430.1      |  1.71|t / c    |EMB3003 &#124; 2-oxoacid dehydrogenases
|
```

```
## |528  |AT1G02500.1 (+1) |  1.71|t / c    |SAM1, SAM-1, MAT1, AtSAM1 &#124; S-aden
|
## |684  |AT3G59970.3      |  1.71|t / c    |MTHFR1 &#124; methylenetetrahydrofolate
|
## |790  |AT1G54270.1      |  1.71|t / c    |EIF4A-2 &#124; eif4a-2 &#124; chr1:20260495
- |
## |1108 |AT5G15090.1 (+1) |  1.70|t / c    |VDAC3, ATVDAC3 &#124; voltage dependent
|
## |1752 |AT1G15930.1 (+1) |  1.70|t / c    |&#124; Ribosomal protein L7Ae/L30e/S12
|
## |75   |AT1G78900.1 (+1) |  1.69|t / c    |VHA-A &#124; vacuolar ATP synthase subu
|
## |765  |AT5G27390.1      |  1.69|t / c    |&#124; Mog1/PsbP/DUF1795-like photosys
|
## |15   |AT1G42970.1      |  1.68|t / c    |GAPB &#124; glyceraldehyde-3-phosphate
|
## |574  |AT2G28900.1      |  1.68|t / c    |OEP16, ATOEP16-L, ATOEP16-1, OEP16
|
## |594  |AT2G05990.1 (+1) |  1.68|t / c    |MOD1, ENR1 &#124; NAD(P)-binding Rossma
|
## |695  |AT2G21870.1      |  1.68|t / c    |MGP1 &#124; copper ion binding;cobalt i
|
## |919  |AT2G33530.1      |  1.68|t / c    |scpl46 &#124; serine carboxypeptidase-l
|
## |972  |AT1G09430.1      |  1.68|t / c    |ACLA-3 &#124; ATP-citrate lyase A-3 &#124;
c |
## |30   |AT5G14740.1      |  1.67|t / c    |CA2, CA18, BETA CA2 &#124; carbonic anh
|
## |79   |AT3G46780.1      |  1.67|t / c    |PTAC16 &#124; plastid transcriptionally
|
## |186  |AT1G72150.1      |  1.67|t / c    |PATL1 &#124; PATELLIN 1 &#124; chr1:2714855
8 |
## |45   |AT5G17920.1 (+1) |  1.66|t / c    |ATCIMS, ATMETS, ATMS1 &#124; Cobalamin-
|
## |165  |AT1G55670.1      |  1.66|t / c    |PSAG &#124; photosystem I subunit G &#124;
c |
## |286  |AT1G09750.1      |  1.66|t / c    |&#124; Eukaryotic aspartyl protease fa
|
## |1065 |AT2G48070.1 (+1) |  1.66|t / c    |RPH1 &#124; resistance to phytophthora
|
## |116  |AT3G52930.1      |  1.65|t / c    |&#124; Aldolase superfamily protein &#124;
|
## |70   |AT2G21390.1      |  1.64|t / c    |&#124; Coatomer, alpha subunit &#124; chr2:
|
## |442  |AT1G35720.1      |  1.64|t / c    |ANNAT1, OXY5, ATOXY5 &#124; annexin 1 &#12
4; |
## |673  |AT1G65260.1      |  1.64|t / c    |PTAC4, VIPP1 &#124; plastid transcripti
|
## |744  |ATCG00670.1      |  1.64|t / c    |CLPP1, PCLPP &#124; plastid-encoded CLP
|
## |942  |AT2G46280.1 (+1) |  1.64|t / c    |TRIP-1, TIF3I1 &#124; TGF-beta receptor
|
```

```
## |1031 |AT1G73600.2      |  1.64|t / c    |&#124; S-adenosyl-L-methionine-depende
|
## |1678 |AT1G74040.1      |  1.64|t / c    |IMS1, MAML-3, IPMS2 &#124; 2-isopropylm
|
## |16   |AT4G20360.1      |  1.63|t / c    |ATRAB8D, ATRABE1B, RABE1b &#124; RAB GT
|
## |299  |AT3G47520.1      |  1.62|t / c    |MDH &#124; malate dehydrogenase &#124; chr
3: |
## |736  |ATCG01090.1      |  1.62|t / c    |NDHI &#124; NADPH dehydrogenases &#124; chr
C |
## |597  |AT1G69740.1 (+1) |  1.60|t / c    |HEMB1 &#124; Aldolase superfamily prote
|
## |1269 |AT2G34470.1 (+1) |  1.58|t-unique |UREG, PSKF109 &#124; urease accessory p
|
## |1697 |AT5G25940.1      |  1.58|t-unique |&#124; early nodulin-related &#124; chr5:90
|
## |1965 |AT3G62550.1      |  1.58|t-unique |&#124; Adenine nucleotide alpha hydrol
|
## |750  |AT2G18710.1      |  1.58|t / c    |SCY1 &#124; SECY homolog 1 &#124; chr2:8112
2 |
## |799  |AT1G64190.1      |  1.58|t / c    |&#124; 6-phosphogluconate dehydrogenas
|
## |808  |AT3G13750.1      |  1.58|t / c    |BGAL1 &#124; beta galactosidase 1 &#124; ch
r |
## |840  |AT5G11770.1      |  1.58|t / c    |&#124; NADH-ubiquinone oxidoreductase
|
## |937  |AT5G13030.1      |  1.58|t / c    |&#124; unknown protein; FUNCTIONS IN:
|
## |957  |AT1G63970.1 (+1) |  1.58|t / c    |ISPF, MECPS &#124; isoprenoid F &#124; chr
1: |
## |1002 |AT1G27450.3      |  1.58|t / c    |APT1 &#124; adenine phosphoribosyl tran
|
## |1119 |AT4G16180.2      |  1.58|t / c    |&#124; unknown protein; FUNCTIONS IN:
|
## |1171 |AT5G45620.1      |  1.58|t / c    |&#124; Proteasome component (PCI) doma
|
## |1275 |AT4G27800.1      |  1.58|t / c    |TAP38, PPH1 &#124; thylakoid-associated
|
## |1280 |AT5G59420.1      |  1.58|t / c    |ORP3C &#124; OSBP(oxysterol binding pro
|
## |1287 |AT1G63710.1      |  1.58|t / c    |CYP86A7 &#124; cytochrome P450, family
|
## |1299 |AT4G10120.1 (+1) |  1.58|t / c    |ATSPS4F &#124; Sucrose-phosphate syntha
|
## |1353 |AT1G70290.1      |  1.58|t / c    |ATTPS8, TPS8, ATTPSC &#124; trehalose-6
|
## |1360 |AT5G13120.1      |  1.58|t / c    |ATCYP20-2, CYP20-2 &#124; cyclophilin 2
|
## |1362 |AT3G61650.1      |  1.58|t / c    |TUBG1 &#124; gamma-tubulin &#124; chr3:2281
2 |
## |1378 |AT1G56050.1      |  1.58|t / c    |&#124; GTP-binding protein-related &#124; c
|
```

```
## |1400 |AT3G56630.1      |  1.58|t / c    |CYP94D2 &#124; cytochrome P450, family
|
## |1436 |AT4G19120.1 (+1) |  1.58|t / c    |ERD3 &#124; S-adenosyl-L-methionine-dep
|
## |1447 |AT2G26910.1      |  1.58|t / c    |PDR4, ATPDR4 &#124; pleiotropic drug re
|
## |1509 |AT1G18730.1      |  1.58|t / c    |NDF6 &#124; NDH dependent flow 6 &#124; chr
1 |
## |1527 |AT5G14120.1      |  1.58|t / c    |&#124; Major facilitator superfamily p
|
## |1532 |AT1G67730.1      |  1.58|t / c    |YBR159, KCR1, ATKCR1 &#124; beta-ketoac
|
## |1551 |AT1G53210.1      |  1.58|t / c    |&#124; sodium/calcium exchanger family
|
## |1576 |AT5G08740.1      |  1.58|t / c    |NDC1 &#124; NAD(P)H dehydrogenase C1 &#124;
|
## |1578 |AT3G07810.2      |  1.58|t / c    |&#124; RNA-binding (RRM/RBD/RNP motifs
|
## |1594 |AT2G41040.1      |  1.58|t / c    |&#124; S-adenosyl-L-methionine-depende
|
## |1614 |AT5G67385.1      |  1.58|t / c    |&#124; Phototropic-responsive NPH3 fam
|
## |1635 |AT5G26710.1      |  1.58|t / c    |&#124; Glutamyl/glutaminyl-tRNA synthe
|
## |1638 |AT1G64770.1      |  1.58|t / c    |NDF2, NDH45 &#124; NDH-dependent cyclic
|
## |1680 |AT2G35790.1      |  1.58|t / c    |&#124; unknown protein; CONTAINS Inter
|
## |1728 |AT1G64520.1      |  1.58|t / c    |RPN12a &#124; regulatory particle non-A
|
## |1729 |AT1G18210.1 (+1) |  1.58|t / c    |&#124; Calcium-binding EF-hand family
|
## |1742 |AT3G48720.1      |  1.58|t / c    |&#124; HXXXD-type acyl-transferase fam
|
## |1852 |AT2G32680.1      |  1.58|t / c    |AtRLP23, RLP23 &#124; receptor like pro
|
## |1855 |AT3G17970.1      |  1.58|t / c    |atToc64-III, TOC64-III &#124; transloco
|
## |1901 |AT1G27970.2      |  1.58|t / c    |NTF2B &#124; nuclear transport factor 2
|
## |1915 |AT4G12310.1      |  1.58|t / c    |CYP706A5 &#124; cytochrome P450, family
|
## |1931 |AT1G65220.1      |  1.58|t / c    |&#124; ARM repeat superfamily protein
|
## |1952 |AT4G11820.1      |  1.58|t / c    |MVA1, HMGS, EMB2778, FKP1 &#124; hydrox
|
## |1975 |AT1G79390.1      |  1.58|t / c    |&#124; unknown protein; Has 30201 Blas
|
## |1987 |AT3G15680.1      |  1.58|t / c    |&#124; Ran BP2/NZF zinc finger-like su
|
## |59   |AT1G31330.1      |  1.57|t / c    |PSAF &#124; photosystem I subunit F &#124;
c |
```

```
## |251  |AT3G23990.1      |  1.57|t / c    |HSP60, HSP60-3B &#124; heat shock prote
|
## |279  |AT5G23120.1      |  1.57|t / c    |HCF136 &#124; photosystem II stability/
|
## |241  |AT1G68010.1      |  1.56|t / c    |HPR, ATHPR1 &#124; hydroxypyruvate redu
|
## |338  |AT4G37910.1      |  1.56|t / c    |mtHsc70-1 &#124; mitochondrial heat sho
|
## |445  |AT1G15690.1      |  1.56|t / c    |AVP1, ATAVP3, AVP-3, AtVHP1;1 &#124; In
|
## |329  |AT1G10760.1      |  1.55|t / c    |SEX1, SOP1, SOP, GWD1, GWD &#124; Pyruv
|
## |352  |AT3G29360.1 (+1) |  1.55|t / c    |&#124; UDP-glucose 6-dehydrogenase fam
|
## |931  |AT5G16010.1      |  1.55|t / c    |&#124; 3-oxo-5-alpha-steroid 4-dehydro
|
## |1606 |AT5G39740.1 (+1) |  1.55|t / c    |OLI7, RPL5B &#124; ribosomal protein L5
|
## |81   |AT2G33800.1      |  1.54|t / c    |&#124; Ribosomal protein S5 family pro
|
## |190  |AT3G44310.1 (+1) |  1.54|t / c    |NIT1, ATNIT1, NITI &#124; nitrilase 1 &#12
4; |
## |103  |AT3G01500.2      |  1.53|t / c    |CA1 &#124; carbonic anhydrase 1 &#124; chr
3: |
## |132  |AT2G31610.1      |  1.53|t / c    |&#124; Ribosomal protein S3 family pro
|
## |341  |AT3G61820.1      |  1.53|t / c    |&#124; Eukaryotic aspartyl protease fa
|
## |1036 |AT5G09900.2      |  1.53|t / c    |EMB2107, RPN5A, MSA &#124; 26S proteaso
|
## |1223 |AT4G24830.1      |  1.53|t / c    |&#124; arginosuccinate synthase family
|
## |84   |AT5G01530.1      |  1.52|t / c    |LHCB4.1 &#124; light harvesting complex
|
## |69   |AT4G13940.1      |  1.51|t / c    |HOG1, EMB1395, SAHH1, MEE58, ATSAH
|
## |93   |AT1G33120.1 (+1) |  1.51|t / c    |&#124; Ribosomal protein L6 family &#124; c
|
## |127  |AT3G04840.1      |  1.51|t / c    |&#124; Ribosomal protein S3Ae &#124; chr3:1
|
## |160  |AT2G13360.1 (+1) |  1.51|t / c    |AGT, AGT1, SGAT &#124; alanine:glyoxyla
|
## |427  |AT1G79920.2      |  1.51|t / c    |&#124; Heat shock protein 70 (Hsp 70)
|
## |764  |AT1G09780.1      |  1.51|t / c    |&#124; Phosphoglycerate mutase, 2,3-bi
|
## |1183 |AT2G40100.1      |  1.50|t / c    |LHCB4.3 &#124; light harvesting complex
|
## |41   |AT1G15820.1      |  1.49|t / c    |LHCB6, CP24 &#124; light harvesting com
|
## |158  |AT5G54270.1      |  1.49|t / c    |LHCB3, LHCB3*1 &#124; light-harvesting
|
```

```
## |205  |AT2G36530.1      |  1.49|t / c    |LOS2, ENO2 &#124; Enolase &#124; chr2:15321
0 |
## |468  |AT2G46820.1 (+1) |  1.49|t / c    |PTAC8, TMP14, PSAP, PSI-P &#124; photos
|
## |1173 |AT5G44070.1      |  1.49|t / c    |CAD1, ARA8, ATPCS1, PCS1 &#124; phytoch
|
## |569  |AT3G23400.1      |  1.47|t / c    |FIB4 &#124; Plastid-lipid associated pr
|
## |585  |AT1G01300.1      |  1.47|t / c    |&#124; Eukaryotic aspartyl protease fa
|
## |757  |ATCG00660.1      |  1.47|t / c    |RPL20 &#124; ribosomal protein L20 &#124; c
h |
## |11   |ATCG00480.1      |  1.46|t / c    |ATPB, PB &#124; ATP synthase subunit be
|
## |860  |AT4G30810.1      |  1.46|t / c    |scpl29 &#124; serine carboxypeptidase-l
|
## |114  |AT3G17390.1      |  1.45|t / c    |MTO3, SAMS3, MAT4 &#124; S-adenosylmeth
|
## |308  |AT5G47210.1      |  1.45|t / c    |&#124; Hyaluronan / mRNA binding famil
|
## |850  |AT4G38510.5      |  1.45|t / c    |&#124; ATPase, V1 complex, subunit B p
|
## |311  |AT4G09000.1      |  1.43|t / c    |GRF1, GF14 CHI &#124; general regulator
|
## |55   |AT3G12780.1      |  1.42|t / c    |PGK1 &#124; phosphoglycerate kinase 1 &#12
4; |
## |302  |AT1G04410.1      |  1.42|t / c    |&#124; Lactate/malate dehydrogenase fa
|
## |683  |AT4G23850.1      |  1.42|t / c    |LACS4 &#124; AMP-dependent synthetase a
|
## |685  |AT3G58140.1      |  1.42|t / c    |&#124; phenylalanyl-tRNA synthetase cl
|
## |728  |AT4G38740.1      |  1.42|t / c    |ROC1 &#124; rotamase CYP 1 &#124; chr4:1808
3 |
## |876  |AT3G06050.1      |  1.42|t / c    |PRXIIF, ATPRXIIF &#124; peroxiredoxin I
|
## |886  |AT5G25757.1 (+1) |  1.42|t / c    |&#124; RNA polymerase I-associated fac
|
## |888  |AT3G20390.1      |  1.42|t / c    |&#124; endoribonuclease L-PSP family p
|
## |924  |ATCG01130.1      |  1.42|t / c    |YCF1.2 &#124; Ycf1 protein &#124; chrC:1238
8 |
## |998  |ATCG00190.1      |  1.42|t / c    |RPOB &#124; RNA polymerase subunit beta
|
## |1040 |AT5G20830.1 (+1) |  1.42|t / c    |SUS1, ASUS1, atsus1 &#124; sucrose synt
|
## |1054 |AT3G27925.1      |  1.42|t / c    |DEGP1, Deg1 &#124; DegP protease 1 &#124; c
h |
## |1059 |AT5G24020.1      |  1.42|t / c    |MIND, ARC11, ATMIND1 &#124; septum site
|
## |1109 |AT1G21630.1      |  1.42|t / c    |&#124; Calcium-binding EF hand family
|
```

```
## |1132 |AT3G52180.1      |  1.42|t / c    |ATPTPKIS1, DSP4, SEX4, ATSEX4 &#124; du
|
## |1134 |AT2G01140.1      |  1.42|t / c    |&#124; Aldolase superfamily protein &#124;
|
## |1145 |AT1G30580.1      |  1.42|t / c    |&#124; GTP binding &#124; chr1:10831953-108
|
## |1147 |AT5G20090.1 (+1) |  1.42|t / c    |&#124; Uncharacterised protein family
|
## |1259 |AT4G03430.1      |  1.42|t / c    |STA1, EMB2770 &#124; pre-mRNA splicing
|
## |1264 |AT2G36390.1      |  1.42|t / c    |SBE2.1, BE3 &#124; starch branching enz
|
## |1273 |AT2G14610.1      |  1.42|t / c    |PR1, PR 1, ATPR1 &#124; pathogenesis-re
|
## |1279 |AT5G47040.1      |  1.42|t / c    |LON2 &#124; lon protease 2 &#124; chr5:1909
3 |
## |1297 |AT3G50930.1      |  1.42|t / c    |BCS1 &#124; cytochrome BC1 synthesis &#124;
|
## |1330 |AT1G62180.1      |  1.42|t / c    |APR2, APSR, PRH43, PRH, ATAPR2 &#124; 5
|
## |1373 |AT5G50530.1 (+1) |  1.42|t / c    |&#124; CBS / octicosapeptide/Phox/Bemp
|
## |1405 |AT1G07990.1      |  1.42|t / c    |&#124; SIT4 phosphatase-associated fam
|
## |1443 |AT1G21440.1      |  1.42|t / c    |&#124; Phosphoenolpyruvate carboxylase
|
## |1465 |AT5G10690.1      |  1.42|t / c    |&#124; pentatricopeptide (PPR) repeat-
|
## |1470 |AT1G19150.1      |  1.42|t / c    |LHCA6, LHCA2*1 &#124; photosystem I lig
|
## |1513 |AT2G47840.1      |  1.42|t / c    |&#124; Uncharacterised conserved prote
|
## |1533 |AT3G46530.1      |  1.42|t / c    |RPP13 &#124; NB-ARC domain-containing d
|
## |1649 |AT2G40660.1      |  1.42|t / c    |&#124; Nucleic acid-binding, OB-fold-l
|
## |1791 |AT1G32550.1      |  1.42|t / c    |FdC1 &#124; 2Fe-2S ferredoxin-like supe
|
## |1835 |AT4G25100.1 (+4) |  1.42|t / c    |FSD1, ATFSD1 &#124; Fe superoxide dismu
|
## |1896 |AT5G48970.1      |  1.42|t / c    |&#124; Mitochondrial substrate carrier
|
## |1897 |AT3G54090.1      |  1.42|t / c    |FLN1 &#124; fructokinase-like 1 &#124; chr
3: |
## |1946 |AT5G14420.1 (+3) |  1.42|t / c    |RGLG2 &#124; RING domain ligase2 &#124; chr
5 |
## |1981 |AT5G40890.1      |  1.42|t / c    |ATCLC-A, CLC-A, CLCA, ATCLCA &#124; chl
|
## |2081 |AT3G52870.1      |  1.42|t / c    |&#124; IQ calmodulin-binding motif fam
|
## |1454 |AT4G24550.2      |  1.42|t-unique |&#124; Clathrin adaptor complexes medi
|
```

```
## |1545 |AT1G48210.1 (+1) |  1.42|t-unique |&#124; Protein kinase superfamily prot
|
## |1657 |AT3G15000.1      |  1.42|t-unique |&#124; cobalt ion binding &#124; chr3:50503
|
## |1686 |AT2G33120.1      |  1.42|t-unique |SAR1, VAMP722, ATVAMP722 &#124; synapto
|
## |346  |AT1G47128.1      |  1.41|t / c    |RD21, RD21A &#124; Granulin repeat cyst
|
## |582  |AT1G56190.1      |  1.41|t / c    |&#124; Phosphoglycerate kinase family
|
## |77   |AT3G11130.1      |  1.40|t / c    |&#124; Clathrin, heavy chain &#124; chr3:34
|
## |491  |AT5G01410.1      |  1.39|t / c    |PDX1, ATPDX1.3, RSR4, PDX1.3, ATPD
|
## |175  |ATCG00340.1      |  1.38|t / c    |PSAB &#124; Photosystem I, PsaA/PsaB pr
|
## |489  |AT1G09640.1      |  1.38|t / c    |&#124; Translation elongation factor E
|
## |807  |AT1G30630.1      |  1.38|t / c    |&#124; Coatomer epsilon subunit &#124; chr1
|
## |833  |AT3G10670.1      |  1.38|t / c    |ATNAP7, NAP7 &#124; non-intrinsic ABC p
|
## |282  |AT4G01050.1      |  1.37|t / c    |TROL &#124; thylakoid rhodanese-like &#124;
|
## |40   |AT3G60750.1      |  1.36|t / c    |&#124; Transketolase &#124; chr3:22454004-2
|
## |260  |AT4G19410.2      |  1.36|t / c    |&#124; Pectinacetylesterase family pro
|
## |540  |AT3G09820.2      |  1.35|t / c    |ADK1 &#124; adenosine kinase 1 &#124; chr3:
3 |
## |576  |AT1G63940.4      |  1.35|t / c    |MDAR6 &#124; monodehydroascorbate reduc
|
## |714  |AT5G60600.2      |  1.35|t / c    |GCPE, ISPG, CSB3, CLB4, HDS &#124; 4-hy
|
## |818  |AT2G41680.1      |  1.35|t / c    |NTRC &#124; NADPH-dependent thioredoxin
|
## |922  |AT1G34000.1      |  1.35|t / c    |OHP2 &#124; one-helix protein 2 &#124; chr
1: |
## |413  |AT5G08280.1      |  1.34|t / c    |HEMC &#124; hydroxymethylbilane synthas
|
## |553  |AT1G53310.1 (+2) |  1.34|t / c    |ATPPC1, PEPC1, ATPEPC1, PPC1 &#124; pho
|
## |754  |AT1G02280.1 (+1) |  1.34|t / c    |TOC33, ATTOC33, PPI1 &#124; translocon
|
## |204  |AT2G39800.3      |  1.33|t / c    |P5CS1 &#124; delta1-pyrroline-5-carboxy
|
## |331  |AT2G21330.1      |  1.33|t / c    |FBA1 &#124; fructose-bisphosphate aldol
|
## |1198 |AT4G36500.1      |  1.32|t-unique |&#124; unknown protein; FUNCTIONS IN:
|
## |1957 |AT3G24570.1      |  1.32|t-unique |&#124; Peroxisomal membrane 22 kDa (Mp
|
```

```
## |2090 |AT1G76405.2      |  1.32|t-unique |&#124; unknown protein; FUNCTIONS IN:
|
## |2147 |AT4G08520.1      |  1.32|t-unique |&#124; SNARE-like superfamily protein
|
## |2159 |AT3G19490.1      |  1.32|t-unique |ATNHD1, NHD1 &#124; sodium:hydrogen ant
|
## |68   |AT5G28540.1      |  1.32|t / c    |BIP1 &#124; heat shock protein 70 (Hsp
|
## |449  |AT1G18080.1      |  1.32|t / c    |ATARCA, RACK1A_AT, RACK1A &#124; Transd
|
## |550  |AT3G52500.1      |  1.32|t / c    |&#124; Eukaryotic aspartyl protease fa
|
## |710  |AT5G27640.2      |  1.32|t / c    |TIF3B1, EIF3B, ATEIF3B-1, EIF3B-1,
|
## |743  |AT5G66420.2      |  1.32|t / c    |&#124; LOCATED IN: cellular_component
|
## |898  |AT5G65020.1      |  1.32|t / c    |ANNAT2 &#124; annexin 2 &#124; chr5:2597391
5 |
## |946  |AT1G70820.1      |  1.32|t / c    |&#124; phosphoglucomutase, putative /
|
## |956  |AT4G31300.2      |  1.32|t / c    |PBA1 &#124; N-terminal nucleophile amin
|
## |961  |AT5G36230.1      |  1.32|t / c    |&#124; ARM repeat superfamily protein
|
## |1064 |AT1G56450.1      |  1.32|t / c    |PBG1 &#124; 20S proteasome beta subunit
|
## |1122 |AT1G28290.1 (+1) |  1.32|t / c    |AGP31 &#124; arabinogalactan protein 31
|
## |1182 |AT3G17240.1 (+1) |  1.32|t / c    |mtLPD2 &#124; lipoamide dehydrogenase 2
|
## |1195 |AT5G27120.1      |  1.32|t / c    |&#124; NOP56-like pre RNA processing r
|
## |1345 |AT1G14345.1      |  1.32|t / c    |&#124; NAD(P)-linked oxidoreductase su
|
## |1348 |AT4G35790.1      |  1.32|t / c    |ATPLDDELTA, PLDDELTA &#124; phospholipa
|
## |1351 |AT3G61260.1      |  1.32|t / c    |&#124; Remorin family protein &#124; chr3:2
|
## |1395 |AT5G16300.1      |  1.32|t / c    |&#124; Vps51/Vps67 family (components
|
## |1549 |AT5G08060.1      |  1.32|t / c    |&#124; unknown protein; FUNCTIONS IN:
|
## |1560 |AT4G33650.2      |  1.32|t / c    |DRP3A &#124; dynamin-related protein 3A
|
## |1725 |AT4G03080.1      |  1.32|t / c    |BSL1 &#124; BRI1 suppressor 1 (BSU1)-li
|
## |1747 |AT1G02120.1      |  1.32|t / c    |VAD1 &#124; GRAM domain family protein
|
## |1754 |AT3G04210.1      |  1.32|t / c    |&#124; Disease resistance protein (TIR
|
## |1755 |AT4G39990.1      |  1.32|t / c    |ATRABA4B, ATRAB11G, ATGB3, RABA4B
|
```

```
## |1767 |AT4G27585.1      |  1.32|t / c    |&#124; SPFH/Band 7/PHB domain-containi
|
## |124  |AT1G32060.1      |  1.31|t / c    |PRK &#124; phosphoribulokinase &#124; chr1:
1 |
## |388  |AT4G02520.1      |  1.31|t / c    |ATGSTF2, ATPM24.1, ATPM24, GST2, G
|
## |1090 |AT4G01850.1 (+1) |  1.30|t / c    |SAM-2, MAT2, SAM2, AtSAM2 &#124; S-aden
|
## |57   |AT4G37930.1      |  1.29|t / c    |SHM1, STM, SHMT1 &#124; serine transhyd
|
## |796  |AT4G12830.1      |  1.29|t / c    |&#124; alpha/beta-Hydrolases superfami
|
## |1169 |AT1G49750.1      |  1.29|t / c    |&#124; Leucine-rich repeat (LRR) famil
|
## |107  |AT2G42600.1 (+1) |  1.28|t / c    |ATPPC2, PPC2 &#124; phosphoenolpyruvate
|
## |598  |AT2G20420.1      |  1.28|t / c    |&#124; ATP citrate lyase (ACL) family
|
## |734  |AT3G23810.1      |  1.28|t / c    |SAHH2, ATSAHH2 &#124; S-adenosyl-l-homo
|
## |34   |AT1G07920.1 (+5) |  1.27|t / c    |&#124; GTP binding Elongation factor T
|
## |334  |AT5G18380.1 (+1) |  1.27|t / c    |&#124; Ribosomal protein S5 domain 2-l
|
## |202  |AT1G57720.1 (+1) |  1.26|t / c    |&#124; Translation elongation factor E
|
## |817  |AT5G15650.1      |  1.26|t / c    |RGP2, ATRGP2 &#124; reversibly glycosyl
|
## |966  |AT1G59900.1      |  1.26|t / c    |AT-E1 ALPHA, E1 ALPHA &#124; pyruvate d
|
## |1005 |AT1G33810.1      |  1.26|t / c    |&#124; unknown protein; FUNCTIONS IN:
|
## |1011 |AT3G51800.1 (+2) |  1.26|t / c    |ATG2, EBP1, ATEBP1 &#124; metallopeptid
|
## |456  |AT4G02930.1      |  1.25|t / c    |&#124; GTP binding Elongation factor T
|
## |465  |AT5G07030.1      |  1.25|t / c    |&#124; Eukaryotic aspartyl protease fa
|
## |519  |AT1G29930.1      |  1.25|t / c    |CAB1, AB140, CAB140, LHCB1.3 &#124; chl
|
## |607  |AT2G24270.1 (+2) |  1.25|t / c    |ALDH11A3 &#124; aldehyde dehydrogenase
|
## |666  |AT3G56910.1      |  1.25|t / c    |PSRP5 &#124; plastid-specific 50S ribos
|
## |994  |AT3G03780.1 (+2) |  1.24|t / c    |ATMS2, MS2 &#124; methionine synthase 2
|
## |1320 |AT4G23460.1      |  1.22|t-unique |&#124; Adaptin family protein &#124; chr4:1
|
## |1488 |AT4G32400.1      |  1.22|t-unique |EMB104, SHS1, EMB42, ATBT1 &#124; Mitoc
|
## |1552 |AT4G30580.1      |  1.22|t-unique |ATS2, EMB1995, LPAT1 &#124; Phospholipi
|
```

```
## |1739 |AT1G74690.1      |  1.22|t-unique |IQD31 &#124; IQ-domain 31 &#124; chr1:28061
4 |
## |1781 |ATCG00740.1      |  1.22|t-unique |RPOA &#124; RNA polymerase subunit alph
|
## |1938 |AT3G47960.1      |  1.22|t-unique |&#124; Major facilitator superfamily p
|
## |386  |AT2G37220.1      |  1.22|t / c    |&#124; RNA-binding (RRM/RBD/RNP motifs
|
## |396  |AT4G13930.1      |  1.22|t / c    |SHM4 &#124; serine hydroxymethyltransfe
|
## |556  |AT3G01280.1      |  1.22|t / c    |VDAC1, ATVDAC1 &#124; voltage dependent
|
## |614  |AT2G24200.1 (+1) |  1.22|t / c    |&#124; Cytosol aminopeptidase family p
|
## |772  |AT5G24650.1      |  1.22|t / c    |&#124; Mitochondrial import inner memb
|
## |910  |AT4G23100.1 (+1) |  1.22|t / c    |RML1, PAD2, GSH1, CAD2, ATECS1, GS
|
## |992  |AT1G47260.1      |  1.22|t / c    |APFI, GAMMA CA2 &#124; gamma carbonic a
|
## |1003 |AT5G19620.1      |  1.22|t / c    |EMB213, OEP80, ATOEP80, TOC75 &#124; ou
|
## |1149 |AT2G20530.1 (+1) |  1.22|t / c    |ATPHB6, PHB6 &#124; prohibitin 6 &#124; chr
2 |
## |1154 |AT2G19520.1      |  1.22|t / c    |FVE, ACG1, MSI4, NFC4, NFC04, ATMS
|
## |1158 |AT2G42790.1      |  1.22|t / c    |CSY3 &#124; citrate synthase 3 &#124; chr2:
1 |
## |1213 |AT5G42740.1      |  1.22|t / c    |&#124; Sugar isomerase (SIS) family pr
|
## |1327 |AT1G64400.1      |  1.22|t / c    |LACS3 &#124; AMP-dependent synthetase a
|
## |1344 |AT5G47500.1      |  1.22|t / c    |&#124; Pectin lyase-like superfamily p
|
## |1347 |AT5G03280.1      |  1.22|t / c    |EIN2, PIR2, CKR1, ERA3, ORE3, ORE2
|
## |1367 |AT1G31410.1      |  1.22|t / c    |&#124; putrescine-binding periplasmic
|
## |1384 |AT5G43060.1      |  1.22|t / c    |&#124; Granulin repeat cysteine protea
|
## |1388 |AT3G10350.1      |  1.22|t / c    |&#124; P-loop containing nucleoside tr
|
## |1430 |AT4G12590.1      |  1.22|t / c    |&#124; Protein of unknown function DUF
|
## |1437 |AT3G02780.1      |  1.22|t / c    |IPP2, IPIAT1, IDI2 &#124; isopentenyl p
|
## |1448 |AT5G51740.1      |  1.22|t / c    |&#124; Peptidase family M48 family pro
|
## |1457 |AT5G45950.1      |  1.22|t / c    |&#124; GDSL-like Lipase/Acylhydrolase
|
## |1462 |AT3G62120.1 (+1) |  1.22|t / c    |&#124; Class II aaRS and biotin synthe
|
```

```
## |1504 |AT1G55450.1 (+1) |  1.22|t / c    |&#124; S-adenosyl-L-methionine-depende
|
## |1512 |AT5G46630.1      |  1.22|t / c    |&#124; Clathrin adaptor complexes medi
|
## |1514 |AT1G02205.3      |  1.22|t / c    |CER1 &#124; Fatty acid hydroxylase supe
|
## |1573 |AT5G23890.1      |  1.22|t / c    |&#124; LOCATED IN: mitochondrion, chlo
|
## |1596 |AT1G52540.1      |  1.22|t / c    |&#124; Protein kinase superfamily prot
|
## |1603 |AT4G29810.1      |  1.22|t / c    |ATMKK2, MKK2, MK1 &#124; MAP kinase kin
|
## |1613 |AT1G71410.1      |  1.22|t / c    |&#124; ARM repeat superfamily protein
|
## |1617 |AT1G22280.1      |  1.22|t / c    |PAPP2C &#124; phytochrome-associated pr
|
## |1624 |AT5G64860.1      |  1.22|t / c    |DPE1 &#124; disproportionating enzyme &#12
4; |
## |1625 |AT3G13180.1      |  1.22|t / c    |&#124; NOL1/NOP2/sun family protein /
|
## |1631 |AT5G38830.1      |  1.22|t / c    |&#124; Cysteinyl-tRNA synthetase, clas
|
## |1692 |AT5G03455.1      |  1.22|t / c    |CDC25, ARATH;CDC25, ACR2 &#124; Rhodane
|
## |1709 |AT5G04900.1      |  1.22|t / c    |NOL &#124; NYC1-like &#124; chr5:1434826-14
3 |
## |1759 |AT5G42765.1      |  1.22|t / c    |&#124; INVOLVED IN: biological_process
|
## |1867 |AT3G49240.1      |  1.22|t / c    |emb1796 &#124; Pentatricopeptide repeat
|
## |1879 |AT5G27350.1      |  1.22|t / c    |SFP1 &#124; Major facilitator superfami
|
## |1893 |AT3G27390.1      |  1.22|t / c    |&#124; unknown protein; FUNCTIONS IN:
|
## |2089 |AT4G28510.1      |  1.22|t / c    |ATPHB1, PHB1 &#124; prohibitin 1 &#124; chr
4 |
## |152  |AT4G32260.1      |  1.21|t / c    |&#124; ATPase, F0 complex, subunit B/B
|
## |29   |AT2G34420.1      |  1.20|t / c    |LHB1B2, LHCB1.5 &#124; photosystem II l
|
## |991  |AT3G10060.1      |  1.20|t / c    |&#124; FKBP-like peptidyl-prolyl cis-t
|
## |201  |AT2G34430.1      |  1.19|t / c    |LHB1B1, LHCB1.4 &#124; light-harvesting
|
## |321  |ATCG00160.1      |  1.19|t / c    |RPS2 &#124; ribosomal protein S2 &#124; chr
C |
## |291  |AT3G63140.1      |  1.18|t / c    |CSP41A &#124; chloroplast stem-loop bin
|
## |430  |AT2G27530.1 (+1) |  1.18|t / c    |PGY1 &#124; Ribosomal protein L1p/L10e
|
## |269  |AT5G19220.1      |  1.17|t / c    |ADG2, APL1 &#124; ADP glucose pyrophosp
|
```

```
## |287  |AT3G11630.1      |  1.17|t / c    |&#124; Thioredoxin superfamily protein
|
## |1445 |AT5G20500.1      |  1.17|t / c    |&#124; Glutaredoxin family protein &#124; c
|
## |1540 |AT4G39460.1 (+1) |  1.17|t / c    |SAMC1, SAMT1 &#124; S-adenosylmethionin
|
## |1902 |AT1G72930.1      |  1.17|t / c    |TIR &#124; toll/interleukin-1 receptor-
|
## |176  |AT3G63490.1      |  1.16|t / c    |&#124; Ribosomal protein L1p/L10e fami
|
## |89   |AT4G38970.1      |  1.15|t / c    |FBA2 &#124; fructose-bisphosphate aldol
|
## |154  |ATCG00540.1      |  1.15|t / c    |PETA &#124; photosynthetic electron tra
|
## |494  |AT4G37925.1      |  1.15|t / c    |NDH-M &#124; subunit NDH-M of NAD(P)H:p
|
## |552  |AT4G14880.1 (+3) |  1.15|t / c    |OASA1, OLD3, CYTACS1 &#124; O-acetylser
|
## |1240 |AT5G42130.1      |  1.15|t / c    |&#124; Mitochondrial substrate carrier
|
## |86   |AT3G20820.1      |  1.14|t / c    |&#124; Leucine-rich repeat (LRR) famil
|
## |324  |AT1G18540.1      |  1.14|t / c    |&#124; Ribosomal protein L6 family pro
|
## |1402 |AT5G51550.1      |  1.14|t / c    |EXL3 &#124; EXORDIUM like 3 &#124; chr5:209
3 |
## |94   |AT5G26742.1 (+1) |  1.13|t / c    |emb1138 &#124; DEAD box RNA helicase (R
|
## |122  |AT1G11860.1 (+2) |  1.13|t / c    |&#124; Glycine cleavage T-protein fami
|
## |534  |AT5G37510.1 (+1) |  1.13|t / c    |EMB1467, CI76 &#124; NADH-ubiquinone de
|
## |619  |AT1G12410.1      |  1.13|t / c    |CLPR2, NCLPP2, CLP2 &#124; CLP protease
|
## |690  |AT4G03520.1      |  1.13|t / c    |ATHM2 &#124; Thioredoxin superfamily pr
|
## |1032 |AT1G78300.1      |  1.13|t / c    |GRF2, 14-3-3OMEGA, GF14 OMEGA &#124; ge
|
## |342  |AT5G11420.1      |  1.12|t / c    |&#124; Protein of unknown function, DU
|
## |571  |AT2G36460.1      |  1.12|t / c    |&#124; Aldolase superfamily protein &#124;
|
## |740  |AT5G04590.1      |  1.12|t / c    |SIR &#124; sulfite reductase &#124; chr5:13
1 |
## |872  |AT3G08530.1      |  1.12|t / c    |&#124; Clathrin, heavy chain &#124; chr3:25
|
## |963  |AT2G34590.1      |  1.12|t / c    |&#124; Transketolase family protein &#124;
|
## |1019 |AT2G44060.1 (+1) |  1.12|t / c    |&#124; Late embryogenesis abundant pro
|
## |61   |AT1G55490.1 (+1) |  1.11|t / c    |CPN60B, LEN1 &#124; chaperonin 60 beta
|
```

```
## |166  |AT3G27690.1      |  1.10|t / c    |LHCB2.4, LHCB2.3, LHCB2 &#124; photosys
|
## |362  |AT4G12800.1      |  1.10|t / c    |PSAL &#124; photosystem I subunit l &#124;
c |
## |1078 |AT5G19370.1      |  1.10|t / c    |&#124; rhodanese-like domain-containin
|
## |37   |AT3G45140.1      |  1.09|t / c    |LOX2, ATLOX2 &#124; lipoxygenase 2 &#124; c
h |
## |512  |AT3G52880.1      |  1.09|t / c    |ATMDAR1, MDAR1 &#124; monodehydroascorb
|
## |839  |AT5G35790.1      |  1.09|t / c    |G6PD1 &#124; glucose-6-phosphate dehydr
|
## |1117 |AT3G06580.1      |  1.09|t / c    |GAL1, GALK &#124; Mevalonate/galactokin
|
## |1166 |AT4G25450.1      |  1.09|t / c    |ATNAP8, NAP8 &#124; non-intrinsic ABC p
|
## |1276 |AT1G07250.1      |  1.09|t / c    |UGT71C4 &#124; UDP-glucosyl transferase
|
## |429  |AT5G35970.1      |  1.08|t / c    |&#124; P-loop containing nucleoside tr
|
## |723  |AT5G67030.1      |  1.08|t / c    |ABA1, LOS6, NPQ2, ATABA1, ZEP, IBS
|
## |812  |AT1G08360.1      |  1.08|t / c    |&#124; Ribosomal protein L1p/L10e fami
|
## |112  |AT1G44575.1      |  1.07|t / c    |NPQ4, PSBS &#124; Chlorophyll A-B bindi
|
## |547  |AT4G17040.1      |  1.07|t / c    |CLPR4 &#124; CLP protease R subunit 4 &#12
4; |
## |1055 |AT4G02620.1      |  1.07|t / c    |&#124; vacuolar ATPase subunit F famil
|
## |1225 |AT1G74880.1      |  1.07|t / c    |NDH-O &#124; NAD(P)H:plastoquinone dehy
|
## |54   |AT5G04140.2      |  1.06|t / c    |GLU1, GLS1, GLUS, FD-GOGAT &#124; gluta
|
## |533  |AT1G76160.1      |  1.06|t / c    |sks5 &#124; SKU5 similar 5 &#124; chr1:2857
8 |
## |697  |AT5G10450.4      |  1.06|t / c    |GRF6 &#124; G-box regulating factor 6 &#12
4; |
## |890  |AT2G33210.1 (+1) |  1.05|t / c    |HSP60-2 &#124; heat shock protein 60-2
|
## |1089 |AT5G59880.1      |  1.05|t / c    |ADF3 &#124; actin depolymerizing factor
|
## |1143 |AT5G23740.1      |  1.05|t / c    |RPS11-BETA &#124; ribosomal protein S11
|
## |529  |AT2G36880.1 (+1) |  1.04|t / c    |MAT3 &#124; methionine adenosyltransfer
|
## |978  |AT5G16050.1      |  1.04|t / c    |GRF5, GF14 UPSILON &#124; general regul
|
## |264  |AT4G24280.1      |  1.02|t / c    |cpHsc70-1 &#124; chloroplast heat shock
|
## |444  |AT1G01090.1      |  1.00|t / c    |PDH-E1 ALPHA &#124; pyruvate dehydrogen
|
```

```
## |511  |AT3G53460.4      |  1.00|t / c    |CP29 &#124; chloroplast RNA-binding pro
|
## |513  |AT5G44020.1      |  1.00|t / c    |&#124; HAD superfamily, subfamily IIIB
|
## |525  |AT1G16880.1      |  1.00|t / c    |&#124; uridylyltransferase-related &#124; c
|
## |730  |AT5G61790.1      |  1.00|t / c    |CNX1, ATCNX1 &#124; calnexin 1 &#124; chr5:
2 |
## |1056 |AT5G06870.1      |  1.00|t / c    |PGIP2, ATPGIP2 &#124; polygalacturonase
|
## |1094 |AT5G56730.1      |  1.00|t / c    |&#124; Insulinase (Peptidase family M1
|
## |1115 |AT5G57290.1 (+2) |  1.00|t / c    |&#124; 60S acidic ribosomal protein fa
|
## |1123 |AT4G12880.1      |  1.00|t / c    |ENODL19, AtENODL19 &#124; early nodulin
|
## |1133 |AT5G63400.1      |  1.00|t / c    |ADK1 &#124; adenylate kinase 1 &#124; chr5:
2 |
## |1170 |AT4G39520.1      |  1.00|t / c    |&#124; GTP-binding protein-related &#124; c
|
## |1196 |AT5G05000.1 (+2) |  1.00|t / c    |TOC34, ATTOC34, OEP34 &#124; translocon
|
## |1246 |AT3G51460.1      |  1.00|t / c    |RHD4 &#124; Phosphoinositide phosphatas
|
## |1312 |AT3G25140.1      |  1.00|t / c    |GAUT8, QUA1 &#124; Nucleotide-diphospho
|
## |1315 |AT5G24490.1      |  1.00|t / c    |&#124; 30S ribosomal protein, putative
|
## |1318 |AT3G45780.1 (+1) |  1.00|t / c    |PHOT1, NPH1, JK224, RPT1 &#124; phototr
|
## |1321 |AT1G58848.1 (+3) |  1.00|t / c    |&#124; Disease resistance protein (CC-
|
## |1357 |AT4G31850.1      |  1.00|t / c    |PGR3 &#124; proton gradient regulation
|
## |1394 |AT3G53700.1      |  1.00|t / c    |MEE40 &#124; Pentatricopeptide repeat (
|
## |1411 |AT5G23140.1      |  1.00|t / c    |CLPP2, NCLPP7 &#124; nuclear-encoded CL
|
## |1412 |AT4G37980.1      |  1.00|t / c    |ELI3-1, ELI3, ATCAD7, CAD7 &#124; elici
|
## |1413 |AT2G22300.1 (+1) |  1.00|t / c    |CAMTA3, SR1 &#124; signal responsive 1
|
## |1421 |AT5G39570.1      |  1.00|t / c    |&#124; FUNCTIONS IN: molecular_functio
|
## |1423 |AT4G36210.3      |  1.00|t / c    |&#124; Protein of unknown function (DU
|
## |1456 |AT1G12230.2      |  1.00|t / c    |&#124; Aldolase superfamily protein &#124;
|
## |1458 |AT3G25660.1      |  1.00|t / c    |&#124; Amidase family protein &#124; chr3:9
|
## |1461 |AT5G19940.1      |  1.00|t / c    |&#124; Plastid-lipid associated protei
|
```

```
## |1469 |AT2G45180.1      |  1.00|t / c    |&#124; Bifunctional inhibitor/lipid-tr
|
## |1507 |AT5G16390.1 (+1) |  1.00|t / c    |CAC1, CAC1A, BCCP, BCCP1 &#124; chlorop
|
## |1525 |AT4G19170.1      |  1.00|t / c    |NCED4, CCD4 &#124; nine-cis-epoxycarote
|
## |1541 |AT3G22630.1      |  1.00|t / c    |PBD1, PRCGB &#124; 20S proteasome beta
|
## |1555 |AT5G03910.1      |  1.00|t / c    |ATATH12, ATH12 &#124; ABC2 homolog 12 &#12
4; |
## |1566 |AT1G74100.1      |  1.00|t-unique |SOT16, ATSOT16, CORI-7, ATST5A &#124; s
|
## |1567 |AT5G48230.2      |  1.00|t-unique |EMB1276, ACAT2 &#124; acetoacetyl-CoA t
|
## |1585 |AT4G11840.1      |  1.00|t / c    |PLDGAMMA3 &#124; phospholipase D gamma
|
## |1591 |AT1G74260.1      |  1.00|t / c    |PUR4 &#124; purine biosynthesis 4 &#124; ch
r |
## |1601 |AT2G19480.1 (+2) |  1.00|t / c    |NFA02, NFA2, NAP1;2 &#124; nucleosome a
|
## |1602 |AT5G04930.1      |  1.00|t / c    |ALA1 &#124; aminophospholipid ATPase 1
|
## |1609 |ATCG00520.1      |  1.00|t / c    |YCF4 &#124; unfolded protein binding &#124;
|
## |1643 |AT4G27060.1      |  1.00|t / c    |TOR1, SPR2, CN &#124; ARM repeat superf
|
## |1658 |AT3G47833.1      |  1.00|t / c    |&#124; unknown protein; BEST Arabidops
|
## |1662 |AT3G53110.1      |  1.00|t / c    |LOS4 &#124; P-loop containing nucleosid
|
## |1672 |AT2G03510.1      |  1.00|t / c    |&#124; SPFH/Band 7/PHB domain-containi
|
## |1700 |AT4G16450.1 (+1) |  1.00|t / c    |&#124; unknown protein; FUNCTIONS IN:
|
## |1706 |AT1G12640.1      |  1.00|t / c    |&#124; MBOAT (membrane bound O-acyl tr
|
## |1707 |AT4G27070.1      |  1.00|t-unique |TSB2 &#124; tryptophan synthase beta-su
|
## |1722 |AT3G49560.1      |  1.00|t / c    |&#124; Mitochondrial import inner memb
|
## |1764 |AT3G04490.1      |  1.00|t / c    |&#124; unknown protein; BEST Arabidops
|
## |1769 |AT2G42770.1      |  1.00|t / c    |&#124; Peroxisomal membrane 22 kDa (Mp
|
## |1778 |AT3G24550.1      |  1.00|t / c    |ATPERK1, PERK1 &#124; proline extensin-
|
## |1813 |AT3G16950.1      |  1.00|t / c    |LPD1, ptlpd1 &#124; lipoamide dehydroge
|
## |1821 |AT1G26120.1      |  1.00|t-unique |ICME-LIKE1 &#124; alpha/beta-Hydrolases
|
## |1822 |AT3G01780.1      |  1.00|t / c    |TPLATE &#124; ARM repeat superfamily pr
|
```

```
## |1823 |AT5G54180.1      |  1.00|t-unique |PTAC15 &#124; plastid transcriptionally
|
## |1825 |AT3G06350.1      |  1.00|t / c    |EMB3004, MEE32 &#124; dehydroquinate de
|
## |1828 |AT5G53580.1      |  1.00|t / c    |&#124; NAD(P)-linked oxidoreductase su
|
## |1847 |AT4G10060.1      |  1.00|t / c    |&#124; Beta-glucosidase, GBA2 type fam
|
## |1869 |AT1G75130.1      |  1.00|t-unique |CYP721A1 &#124; cytochrome P450, family
|
## |1873 |AT5G44410.1      |  1.00|t / c    |&#124; FAD-binding Berberine family pr
|
## |1900 |AT2G07360.1 (+1) |  1.00|t / c    |&#124; SH3 domain-containing protein &#124;
|
## |1906 |AT5G53140.1      |  1.00|t / c    |&#124; Protein phosphatase 2C family p
|
## |1912 |AT5G47700.1 (+1) |  1.00|t / c    |&#124; 60S acidic ribosomal protein fa
|
## |1930 |AT1G09270.1 (+1) |  1.00|t-unique |IMPA-4 &#124; importin alpha isoform 4
|
## |1968 |AT2G32160.2 (+1) |  1.00|t / c    |&#124; S-adenosyl-L-methionine-depende
|
## |2002 |AT4G12390.1      |  1.00|t / c    |PME1 &#124; pectin methylesterase inhib
|
## |2010 |AT4G03020.1 (+1) |  1.00|t / c    |&#124; transducin family protein / WD-
|
## |2026 |AT5G53650.1      |  1.00|t-unique |&#124; unknown protein; FUNCTIONS IN:
|
## |2048 |AT5G18660.1      |  1.00|t / c    |PCB2 &#124; NAD(P)-binding Rossmann-fol
|
## |2053 |AT1G01510.1      |  1.00|t / c    |AN &#124; NAD(P)-binding Rossmann-fold
|
## |2055 |AT5G51010.1      |  1.00|t / c    |&#124; Rubredoxin-like superfamily pro
|
## |2078 |AT5G64090.1      |  1.00|t / c    |&#124; FUNCTIONS IN: molecular_functio
|
## |2149 |AT3G63540.1      |  1.00|t / c    |&#124; Mog1/PsbP/DUF1795-like photosys
|
## |2155 |AT1G43710.1      |  1.00|t / c    |emb1075 &#124; Pyridoxal phosphate (PLP
|
## |2163 |AT5G40170.1      |  1.00|t / c    |AtRLP54, RLP54 &#124; receptor like pro
|
## |2164 |AT4G26070.2 (+1) |  1.00|t / c    |MEK1, NMAPKK, ATMEK1, MKK1 &#124; MAP k
|
## |2172 |AT2G43780.1 (+1) |  1.00|t-unique |&#124; unknown protein; Has 30 Blast h
|
## |2177 |AT2G01720.1      |  1.00|t / c    |&#124; Ribophorin I &#124; chr2:317193-3200
|
## |2225 |AT1G16890.2      |  1.00|t / c    |UBC36, UBC13B &#124; ubiquitin-conjugat
|
## |2227 |AT3G19810.1      |  1.00|t / c    |&#124; Protein of unknown function (DU
|
```

```
## |2260 |AT3G59990.1 (+3) |  1.00|t-unique |MAP2B &#124; methionine aminopeptidase
|
## |2288 |AT4G40050.1      |  1.00|t / c    |&#124; Protein of unknown function (DU
|
## |2308 |AT4G17300.1      |  1.00|t / c    |NS1, OVA8, ATNS1 &#124; Class II aminoa
|
## |1642 |AT5G07320.1      |  0.74|t-unique |&#124; Mitochondrial substrate carrier
|
## |1762 |AT5G16290.1 (+1) |  0.74|t-unique |VAT1 &#124; VALINE-TOLERANT 1 &#124; chr5:5
3 |
## |1837 |AT5G21326.1      |  0.74|t-unique |&#124; Ca2+regulated serine-threonine
|
## |2047 |AT1G22520.1      |  0.74|t-unique |&#124; Domain of unknown function (DUF
|
## |2077 |AT5G40440.1      |  0.74|t-unique |ATMKK3, MKK3 &#124; mitogen-activated p
|
## |2086 |AT3G51510.1      |  0.74|t-unique |&#124; unknown protein; FUNCTIONS IN:
|
## |2302 |AT3G18680.1      |  0.74|t-unique |&#124; Amino acid kinase family protei
|
## |2328 |AT2G43540.1      |  0.74|t-unique |&#124; unknown protein; FUNCTIONS IN:
|
## |1983 |AT5G16840.1 (+1) |  0.58|t-unique |BPA1 &#124; binding partner of acd11 1
|
## |2032 |AT2G37550.1 (+1) |  0.58|t-unique |ASP1, AGD7 &#124; ARF-GAP domain 7 &#124; c
h |
## |2079 |AT4G39090.1      |  0.58|t-unique |RD19, RD19A &#124; Papain family cystei
|
## |1914 |AT1G67310.1      |  0.42|t-unique |&#124; Calmodulin-binding transcriptio
|
## |1990 |AT1G70760.1      |  0.42|t-unique |CRR23 &#124; inorganic carbon transport
|
## |2130 |AT4G35230.1      |  0.42|t-unique |BSK1 &#124; BR-signaling kinase 1 &#124; ch
r |
## |2217 |AT3G51610.1      |  0.42|t-unique |&#124; unknown protein; FUNCTIONS IN:
|
## |1826 |AT5G54110.1      |  0.00|t-unique |ATMAMI, MAMI &#124; membrane-associated
|
## |2204 |AT1G22850.1      |  0.00|t-unique |&#124; SNARE associated Golgi protein
|
## |2224 |AT5G37360.1      |  0.00|t-unique |&#124; unknown protein; FUNCTIONS IN:
|
## |2403 |AT1G21900.1      |  0.00|t-unique |&#124; emp24/gp25L/p24 family/GOLD fam
|
## |2153 |AT5G04600.1      | -0.58|c-unique |&#124; RNA-binding (RRM/RBD/RNP motifs
|
## |2237 |AT1G51810.1      | -0.81|c-unique |&#124; Leucine-rich repeat protein kin
|
## |1185 |AT4G12060.1      | -1.00|t / c    |&#124; Double Clp-N motif protein &#124; ch
|
## |1490 |AT1G67280.1      | -1.00|t / c    |&#124; Glyoxalase/Bleomycin resistance
|
```

```
## |1740 |AT5G67590.1      | -1.00|t / c    |FRO1 &#124; NADH-ubiquinone oxidoreduct
|
## |1782 |AT3G62820.1      | -1.00|t / c    |&#124; Plant invertase/pectin methyles
|
## |1859 |AT4G13840.1      | -1.00|t / c    |&#124; HXXXD-type acyl-transferase fam
|
## |360  |AT3G05560.1 (+2) | -1.09|t / c    |&#124; Ribosomal L22e protein family &#124;
|
## |434  |AT3G55410.1      | -1.11|t / c    |&#124; 2-oxoglutarate dehydrogenase, E
|
## |907  |AT1G06110.1      | -1.12|t / c    |SKIP16 &#124; SKP1/ASK-interacting prot
|
## |878  |AT2G30870.1      | -1.13|t / c    |ATGSTF10, ERD13, ATGSTF4, GSTF10 &#124;
|
## |1012 |AT3G63190.1      | -1.15|t / c    |RRF, HFP108, cpRRF, AtcpRRF &#124; ribo
|
## |883  |AT3G60190.1      | -1.17|t / c    |ADL4, ADLP2, EDR3, DRP1E, ADL1E, D
|
## |1630 |AT5G17710.1 (+1) | -1.17|t / c    |EMB1241 &#124; Co-chaperone GrpE family
|
## |2006 |AT5G10560.1      | -1.17|t / c    |&#124; Glycosyl hydrolase family prote
|
## |1288 |AT5G63310.1      | -1.22|t / c    |NDPK2, NDPK1A, NDPK IA IA, NDPK IA
|
## |1017 |AT2G41475.1      | -1.25|t / c    |&#124; Embryo-specific protein 3, (ATS
|
## |1006 |AT5G51750.1      | -1.26|t / c    |ATSBT1.3, SBT1.3 &#124; subtilase 1.3 &#12
4; |
## |1397 |AT3G58570.1      | -1.26|t / c    |&#124; P-loop containing nucleoside tr
|
## |777  |AT2G25080.1      | -1.29|t / c    |ATGPX1, GPX1 &#124; glutathione peroxid
|
## |148  |AT1G52400.1 (+1) | -1.30|t / c    |BGL1, BGLU18, ATBG1 &#124; beta glucosi
|
## |795  |AT5G57655.2      | -1.32|t / c    |&#124; xylose isomerase family protein
|
## |1489 |AT5G55220.1      | -1.32|t / c    |&#124; trigger factor type chaperone f
|
## |1013 |AT2G33450.1      | -1.39|t / c    |&#124; Ribosomal L28 family &#124; chr2:141
|
## |1760 |AT5G14910.1      | -1.46|c-unique |&#124; Heavy metal transport/detoxific
|
## |288  |AT4G22010.1      | -1.46|t / c    |sks4 &#124; SKU5  similar 4 &#124; chr4:116
6 |
## |1261 |AT3G52990.1      | -1.58|t / c    |&#124; Pyruvate kinase family protein
|
## |2154 |AT2G04520.1 (+3) | -1.58|t / c    |&#124; Nucleic acid-binding, OB-fold-l
|
## |599  |AT5G11720.1      | -1.63|t / c    |&#124; Glycosyl hydrolases family 31
|
## |1057 |AT4G11010.1      | -1.70|t / c    |NDPK3 &#124; nucleoside diphosphate kin
|
```

```
## |1517 |AT4G21650.1      | -1.91|t / c    |&#124; Subtilase family protein &#124; chr4
|
## |1326 |AT3G43980.1 (+2) | -2.00|t / c    |&#124; Ribosomal protein S14p/S29e fam
|
## |1569 |AT3G08740.1      | -2.00|t / c    |&#124; elongation factor P (EF-P) fami
|
## |1352 |AT5G42980.1      | -2.17|t / c    |ATTRX3, ATH3, ATTRXH3, TRXH3, TRX3
|
## |1194 |AT5G55660.1      | -2.32|t / c    |&#124; DEK domain-containing chromatin
|
## |23   |AT4G35310.1      | -2.52|t / c    |CPK5, ATCPK5 &#124; calmodulin-domain p
|
## |14   |AT1G28380.1      | -8.79|t / c    |NSL1 &#124; MAC/Perforin domain-contain
|
##
##
## <!-- -->
```

### The end of script
